## SupplementaryMaterials for "Toward Robust Neuroanatomical Normative Models: Influence of Sample Size and Covariates Distributions"

### Supplementary Materials

### 1. Supplement to main analyses (OASIS-3)

#### 1.1. Model fit evaluation

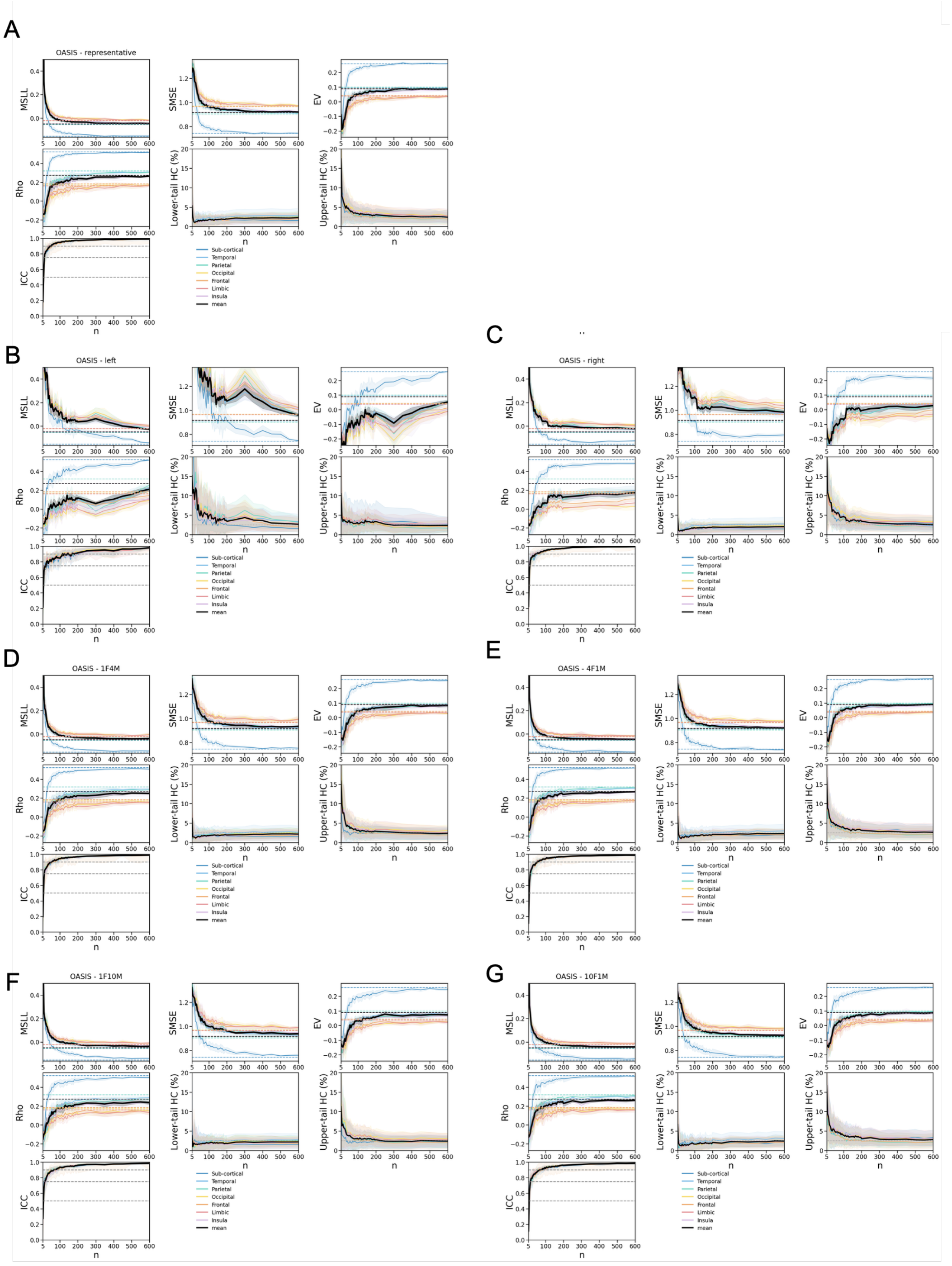

*Figure S1. Model fit evaluation of normative models trained within OASIS-3. Performance is assessed in Healthy Controls (HC) test set of OASIS-3 for different sampling strategies of the training set, including (A) Representative, (B) Left-skewed, and (C) Right-skewed age distributions, as well as (D, E, F, G) sex-imbalanced adaptations with female-to-male ratios of 1:4, 4:1, 1:10, and 10:1, respectively. Model performance is assessed as a function of the adaptation set size (n) using the HC test set from the OASIS-3 dataset. Each panel presents the following evaluation metrics: Mean Standardized Log Loss (MSLL), Standardized Mean Squared Error (SMSE), Explained Variance (EV), Pearson correlation coefficient (Rho), Intraclass Correlation Coefficient (ICC), lower-tail HC percentage (below the 2.5% bound), upper-tail HC percentage (above the 97.5% bound). Solid lines indicate the mean performance across all regions within each lobe (as well as the mean of subcortical regions), while shaded areas represent the standard deviation across 10 iterations*

per sample size. In ICC plots, grey dashed lines denote commonly accepted reliability thresholds: <0.5 (poor), 0.5–0.75 (moderate), 0.75–0.9 (good), and >0.9 (excellent) and shaded areas indicate standard deviation across ROI.

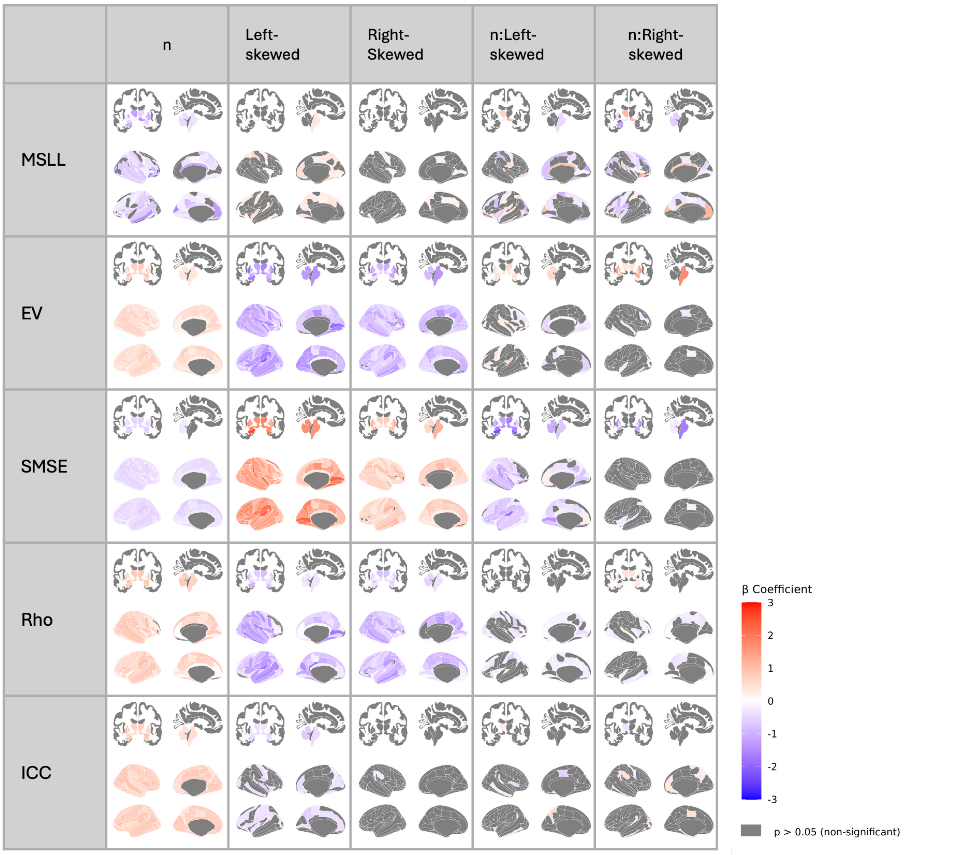

*Figure S2. Regional linear mixed-effects results for model-fit metrics in models trained in OASIS-3 with age-skewed samples. Standardized Betas (β) for each performance metric (MSLL, SMSE, EV, Rho, ICC; rows) are regressed on log-standardized sample size (n), age-distribution contrasts (left-skewed, right-skewed; representative = reference), and their interactions. Sex ratio is balanced (1:1). Grey shading denotes regions that did not survive FDR correction (p ≥ 0.05).*

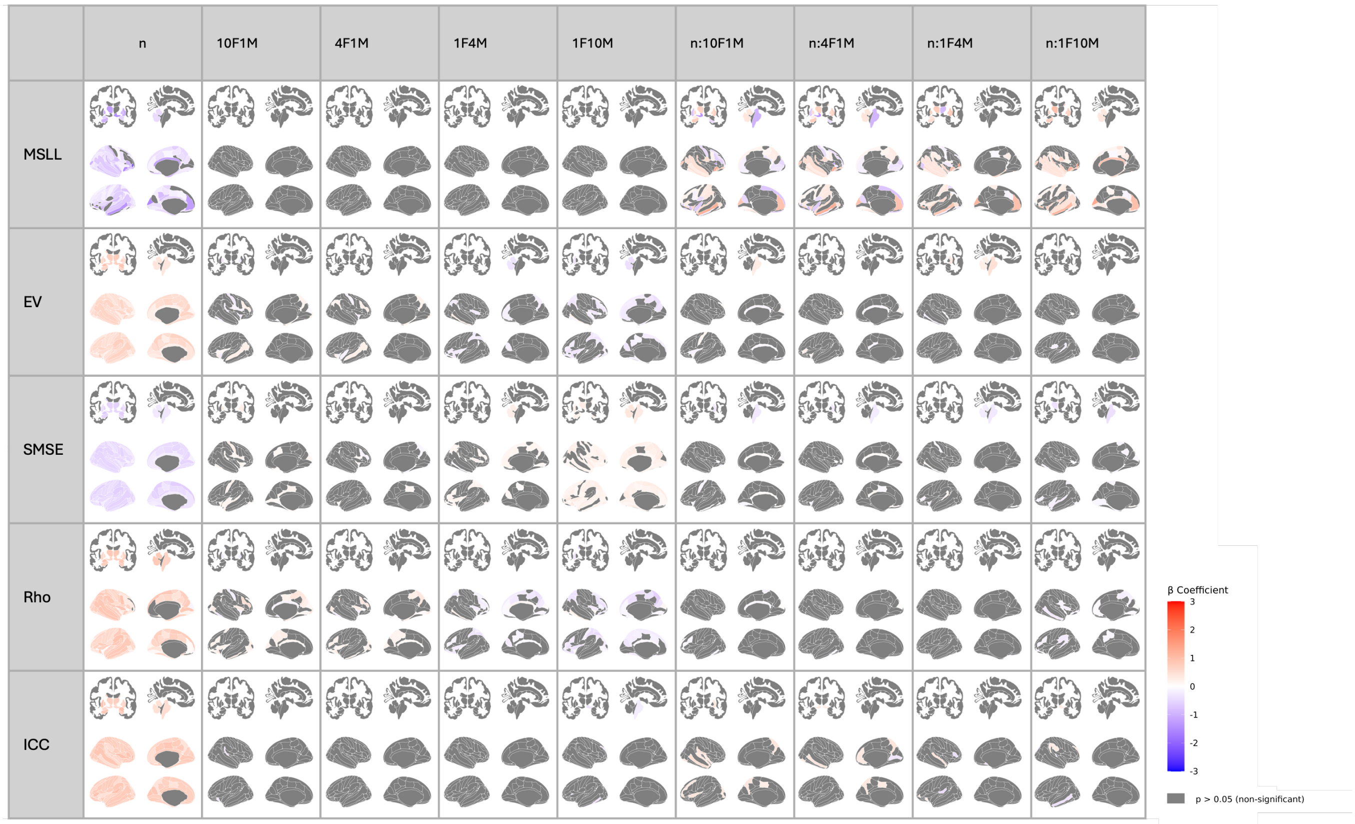

*Figure S3. Regional linear mixed-effects results for model-fit metrics in models trained in OASIS-3 with sex-imbalanced samples. Standardized Betas (β) for each performance metric (MSLL, SMSE, EV, Rho, ICC; rows) are regressed on log-standardized sample size (n), sex-ratio contrasts (female-to-male 10:1, 4:1, 1:1 [reference], 1:4, 1:10), and their interactions. Age distribution is representative. Grey shading denotes regions that did not survive FDR correction (p ≥ 0.05).*

#### 1.2. Clinical validation

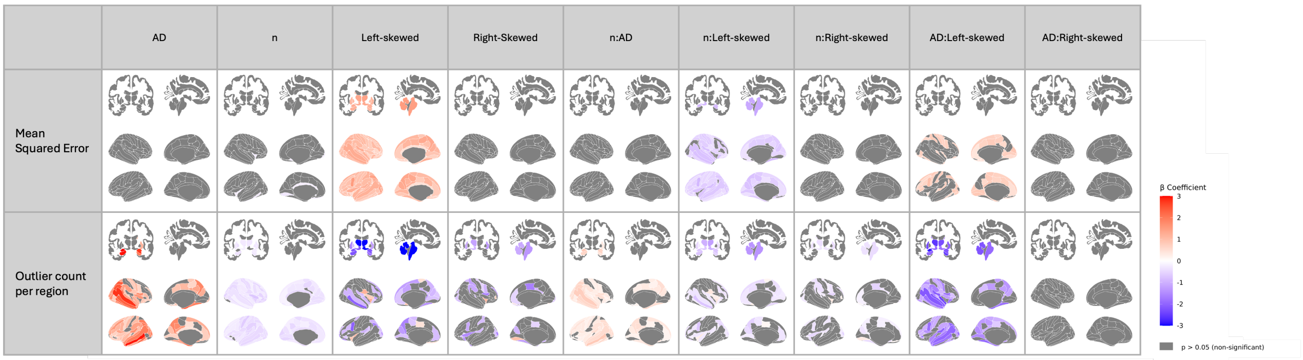

*Figure S4. Regional linear mixed-effects results for deviation-score metrics in models trained in OASIS-3 with age-skewed samples. Standardized Betas (β) are shown for the mean-squared error of z-scores relative to the full reference model and for the count of extreme z-score outliers per region (rows). Predictors include Alzheimer's disease diagnosis (AD), log-standardized sample size (n), age-distribution contrasts (left-skewed, right-skewed; representative = reference), and their two-way interactions with n and AD. Sex ratio is balanced (1:1). Grey shading denotes regions that did not survive FDR correction (p ≥ 0.05).*

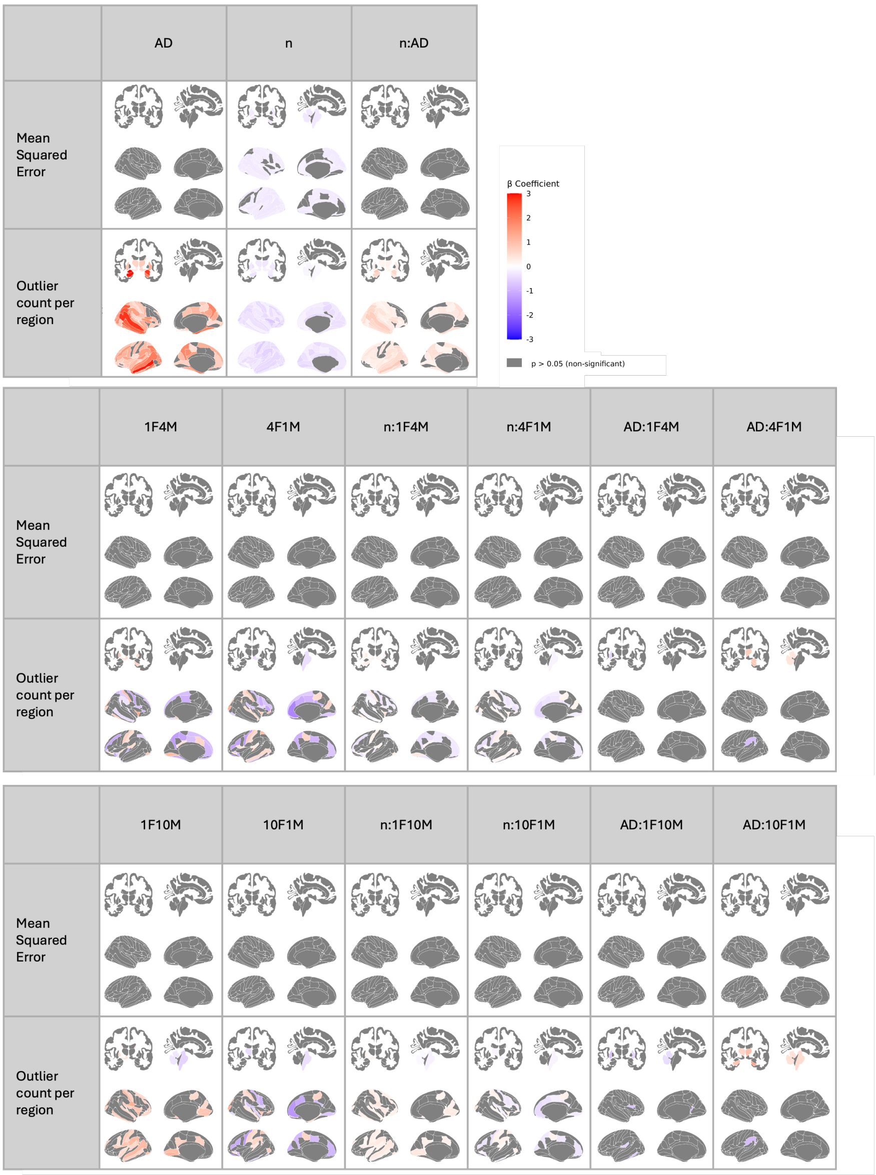

*Figure S5. Regional linear mixed-effects results for deviation-score metrics in models trained in OASIS-3 with sex-imbalanced samples. Standardized Betas (β) are shown for the mean-squared error of z-scores relative to the full reference model and for the count of extreme z-score outliers per region (rows). Predictors include Alzheimer's disease diagnosis (AD), log-standardized sample size (n), sex-ratio contrasts (female-to-male 10:1, 4:1, 1:1 [reference], 1:4, 1:10), and their two-way interactions with n and AD. Age distribution is representative. Grey shading denotes regions that did not survive FDR correction ($p \geq 0.05$).*

#### 1.3. Site effects

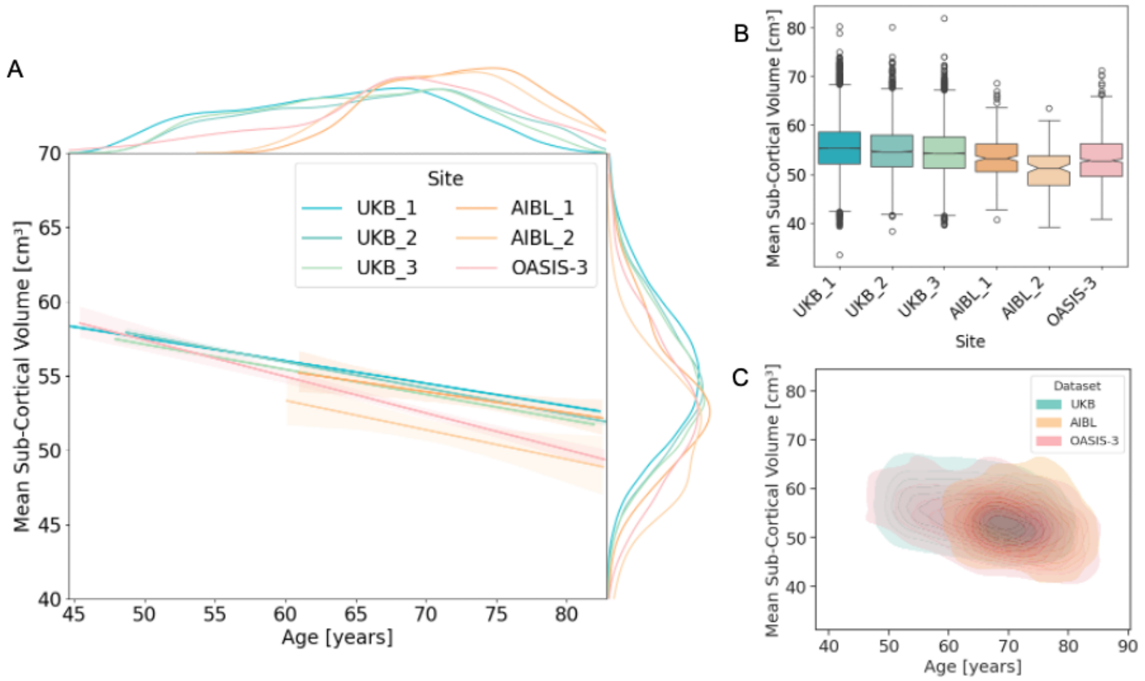

*Figure S6. Site Effects on Sub-Cortical Grey Matter Volume in the UK Biobank (UKB), AIBL, and OASIS-3 Datasets for Healthy Controls. A: Regression plots showing the relationship between age and mean sub-cortical volume across different datasets and imaging sites (UKB: 3 sites, AIBL: 2 sites, OASIS: 2 sites). Marginal density plots are displayed on the sides of each axis, illustrating the distribution of mean sub-cortical volume and age for each dataset's imaging sites. B: Boxplots representing the mean sub-cortical volume for participants in each dataset and imaging site. C: Bivariate kernel density estimates plots showing the joint distribution of age and mean sub-cortical volume for each dataset. The contour plots represent the density of data points, with filled areas reflecting higher concentrations of data.*

#### 1.4. Cohen's d effect sizes

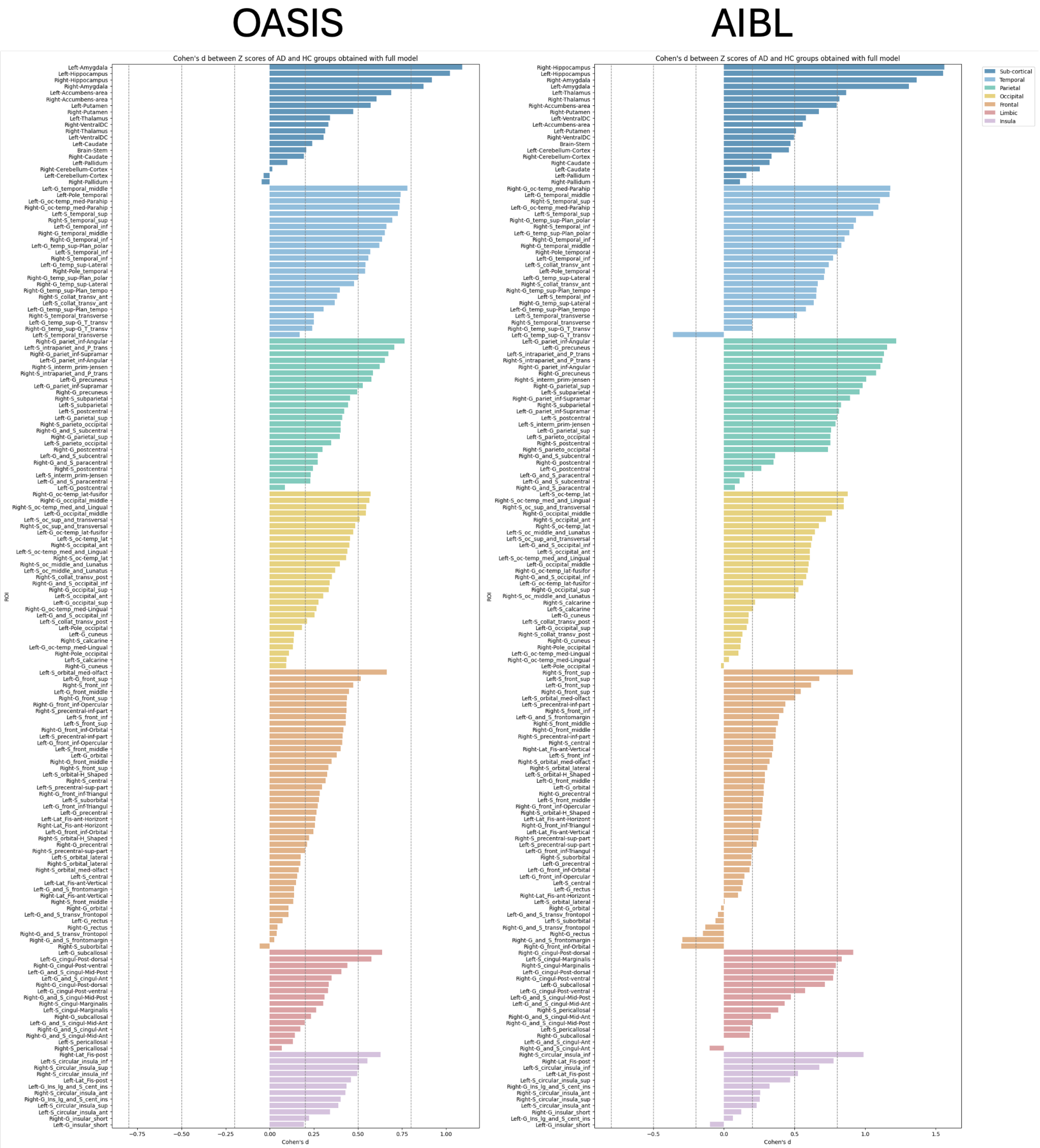

*Figure S7. Cohen's d effect sizes computed on full models' Z-scores for discriminating HC from AD groups for each ROI. Dashed lines correspond to Cohen's classification thresholds for effect sizes: small (d < 0.5), medium (0.5 ≤ d < 0.8), and large (d ≥ 0.8). Left and right panels show Cohen's d effect sizes for OASIS-3 and AIBL test sets, respectively.*

#### 1.5. Additional results

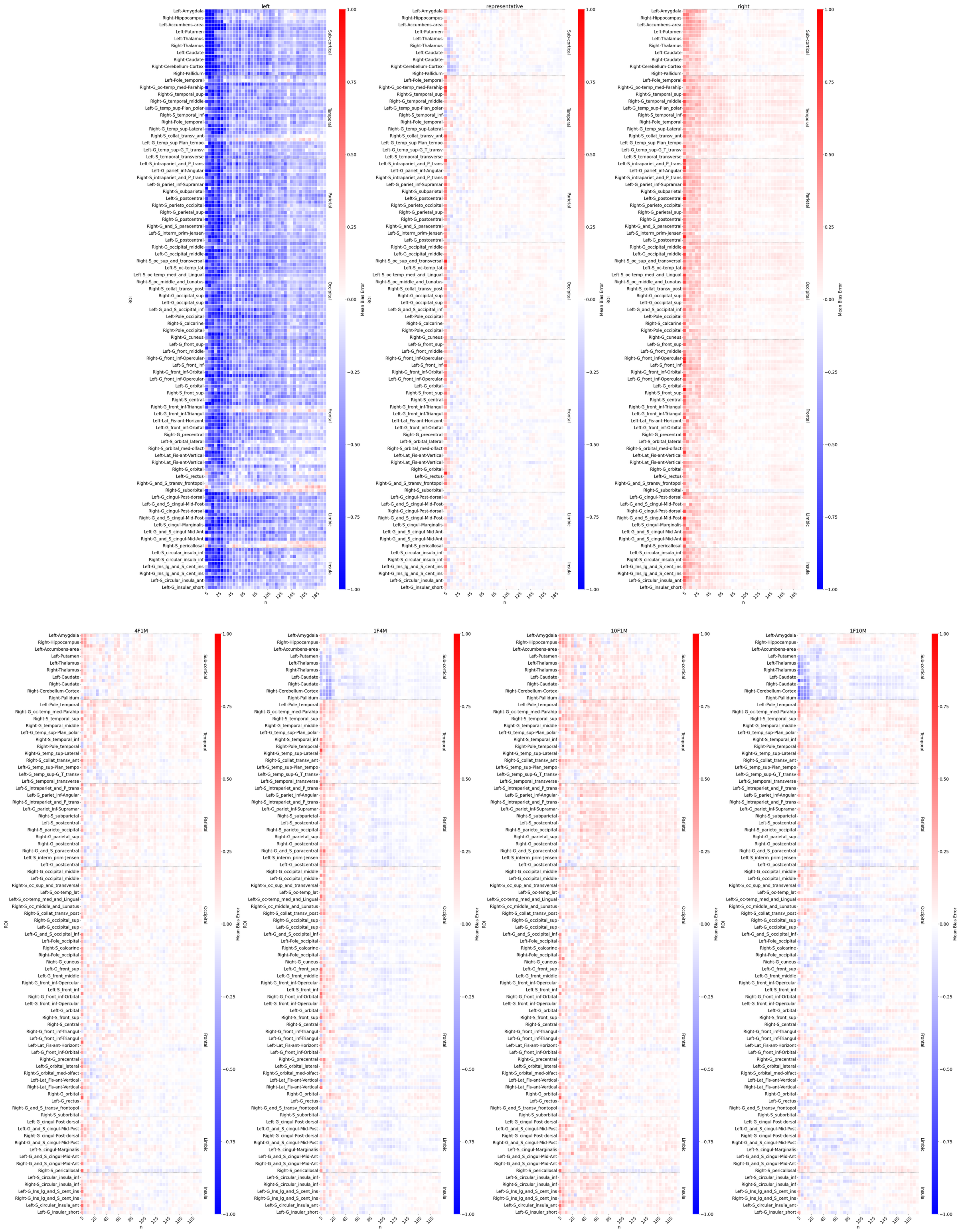

*Figure S8. Mean bias error on z-scores per region in the AD test set relative to the full models in the OASIS-3 dataset, averaged across iterations, as a function of n. Age-skewed sampling strategies include representative (matching the initial age distribution with balanced sex, 1F:1M), left-skewed (favoring younger individuals), and right-skewed (favoring older individuals), each with balanced sex distributions. Sex-imbalanced sampling strategies include female-to-male ratios of 1:1.1, 1:4, 1:10, 4:1, and 10:1, all with representative age distributions.*

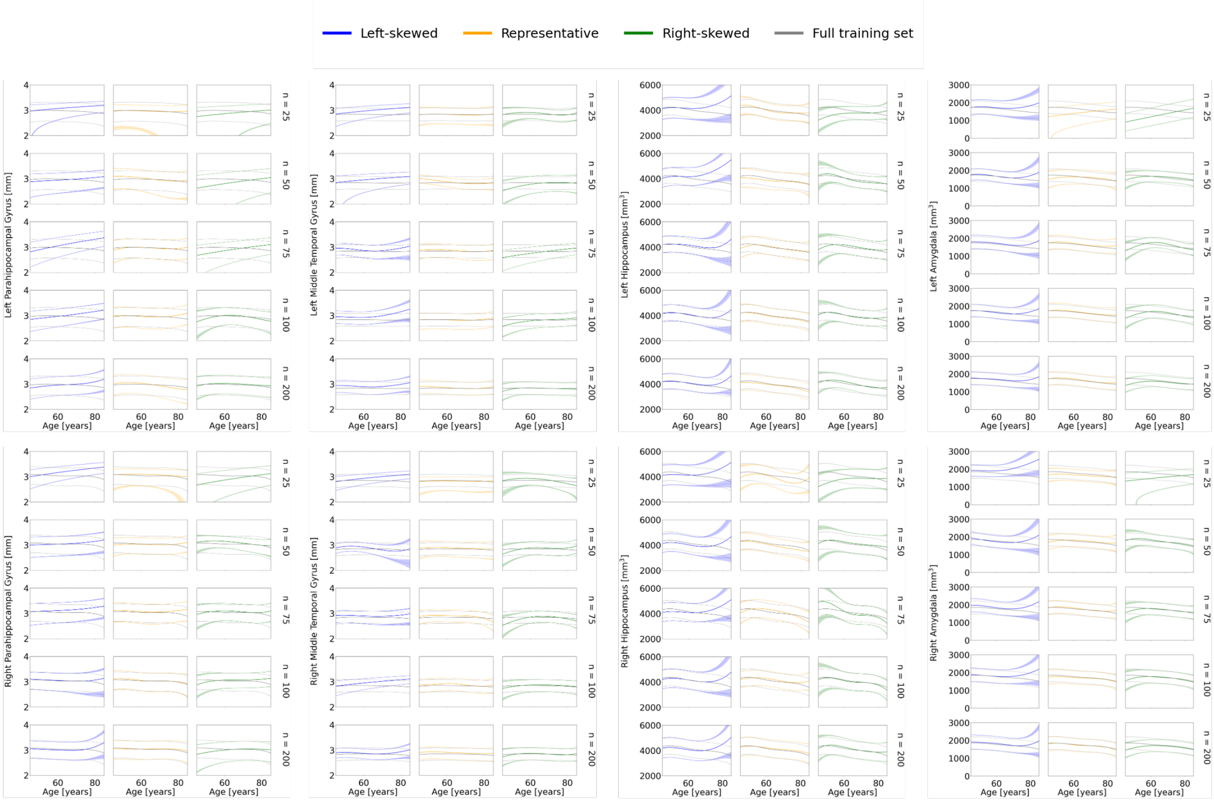

*Figure S9. Centile curve overlays for selected cortical and subcortical regions across sampling strategies and sample sizes. For each ROI, models trained using representative, left-skewed, and right-skewed sampling are shown for multiple sample sizes. Colored lines depict the 5th, 50th, and 95th percentiles estimated from each model, while grey lines indicate the corresponding centiles from the full training set. These overlays highlight how centile estimates diverge when age coverage is limited, particularly at the extreme of age ranges.*

### 2. Replication with Adaptive Transfer Learning (OASIS-3)

#### 2.1. Model fit evaluation

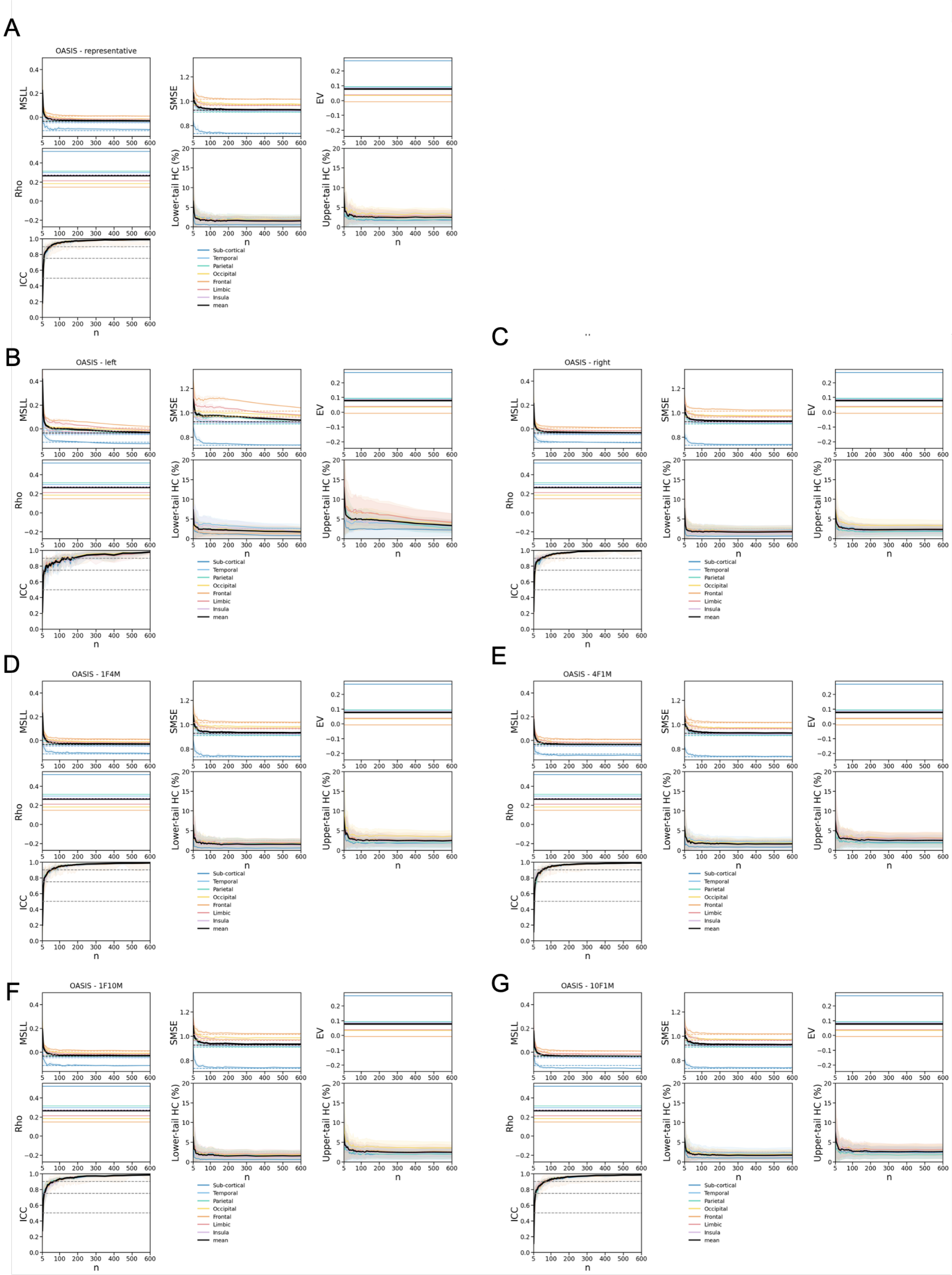

*Figure S10. Model fit evaluation of normative models pre-trained on the UKB and adapted to OASIS-3. Performance is assessed in Healthy Controls (HC) test set of OASIS-3 dataset for different sampling strategies of the adaptation set, including (A) Representative, (B) Left-skewed, and (C) Right-skewed age distributions, as well as (D, E, F, G) sex-imbalanced adaptations with female-to-male ratios of 1:4, 4:1, 1:10, and 10:1, respectively. Model performance is assessed as a function of the adaptation set size (n) using the HC test set from the OASIS-3 dataset. Each panel presents the following evaluation metrics: Mean Standardized Log Loss (MSLL), Standardized Mean Squared Error (SMSE), Explained Variance (EV), Pearson correlation coefficient (Rho), Intraclass Correlation Coefficient (ICC), lower-tail HC percentage (below the 2.5% bound), upper-tail HC percentage (above the 97.5% bound). Solid lines indicate the mean performance across all regions within each lobe (as well as the mean of subcortical regions), while shaded areas*

*represent the standard deviation across 10 iterations per sample size. In ICC plots, grey dashed lines denote commonly accepted reliability thresholds: <0.5 (poor), 0.5–0.75 (moderate), 0.75–0.9 (good), and >0.9 (excellent) and shaded areas indicate standard deviation across ROI.*

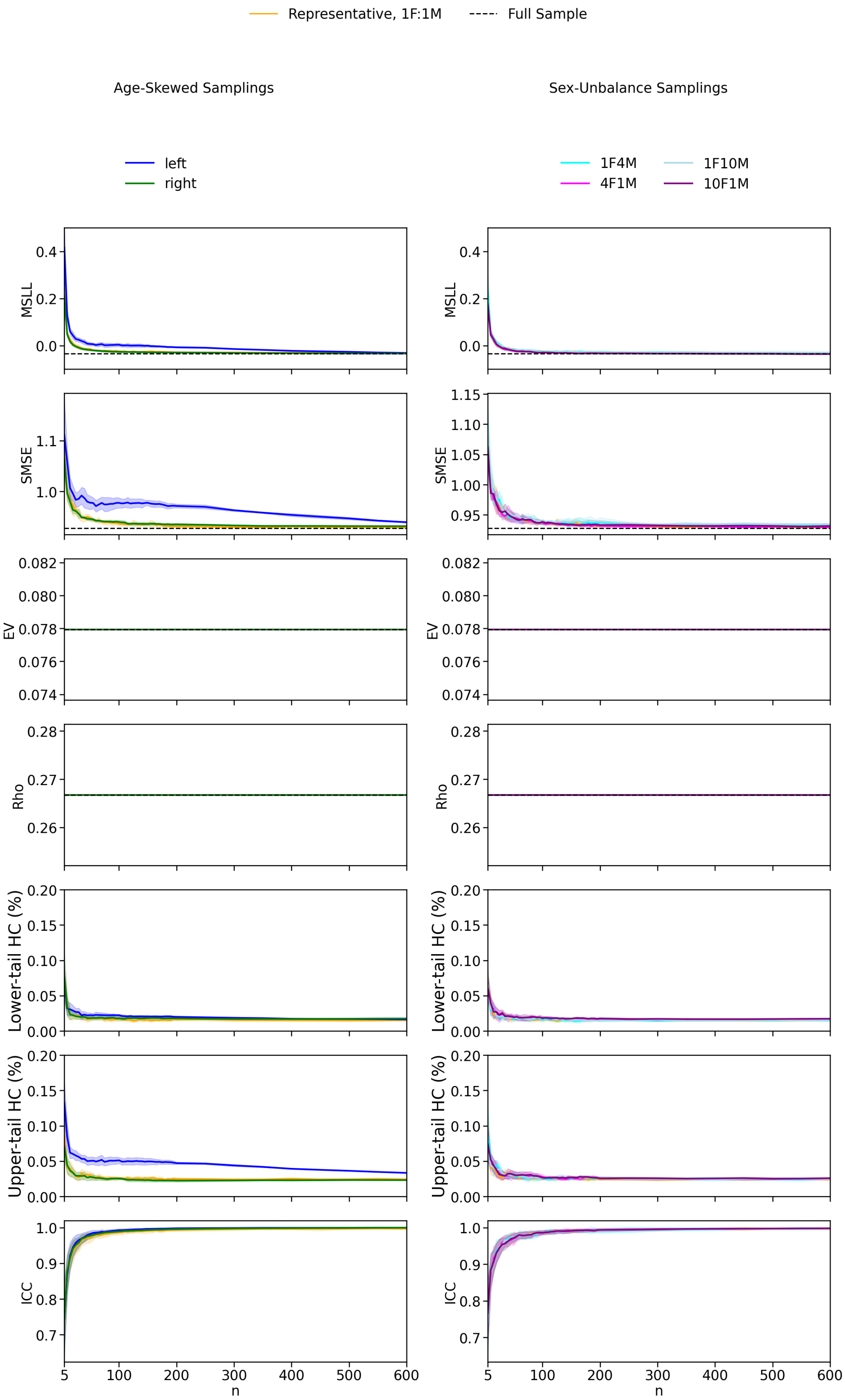

Representative, 1F:1M
Full Sample
Age-Skewed Samplings
Sex-Unbalance Samplings
left
right
1F4M
4F1M
1F10M
10F1M
MSLL
SMSE
EV
Rho
Lower-tail HC (%)
Upper-tail HC (%)
ICC
n

*Figure S11. Model fit evaluation of normative models pre-trained on UKB and adapted to OASIS-3 across different sampling strategies. Models were evaluated in the HC test set for various adaptation sample sizes (n). Age-skewed sampling strategies include representative (matching the initial age distribution with balanced sex, 1F:1M), left-skewed (favoring younger individuals), and right-skewed (favoring older individuals), each with balanced sex distributions. Sex-imbalanced sampling strategies include female-to-male ratios of 1:1.1, 1:4, 1:10, 4:1, and 10:1, all with representative age distributions. Solid lines represent the mean metric values across ROIs and iterations, with shaded areas indicating the standard deviation across iterations for MSLL, SMSE, EV, and Rho. For ICC, shaded areas represent variation across ROIs, as ICC already reflects variability across iteration. Dashed lines indicate the mean metric values for models trained with the full sample for MSLL, SMSE, EV, Rho, the lower and upper tail HC%.*

| | ***MSLL* β (p-value)** | ***SMSE* β (p-value)** | ***EV* β (p-value)** | ***Rho* β (p-value)** | ***ICC* β (p-value)** | ***Lower tail HC %* β (p-value)** | ***Upper tail HC %* β (p-value)** |
|---|---|---|---|---|---|---|---|
| ***Intercept (Representative)*** | -0.195 (p < .001) | -0.567 (p < .001) | 0.644 (p < .001) | 0.701 (p < .001) | 0.504 (p < .001) | -0.219 (p < .001) | -0.373 (p < .001) |
| ***Log(n)*** | -0.054 (p < .001) | -0.102 (p < .001) | 0.000 (p = 1.000) | 0.000 (p = 1.000) | 0.245 (p < .001) | -0.228 (p < .001) | -0.210 (p < .001) |
| ***Left-skewed*** | 0.062 (p < .001) | 0.220 (p < .001) | 0.000 (p = 1.000) | -0.000 (p = 1.000) | 0.041 (p < .001) | 0.186 (p < .001) | 0.778 (p < .001) |
| ***Right-skewed*** | -0.005 (p < .001) | 0.006 (p = .049) | -0.000 (p = 1.000) | -0.000 (p = 1.000) | 0.020 (p < .001) | 0.076 (p < .001) | -0.057 (p < .001) |
| ***Log(n):Left-skewed*** | -0.037 (p < .001) | -0.019 (p < .001) | 0.000 (p = 1.000) | 0.000 (p = 1.000) | -0.002 (p = .645) | -0.010 (p = .133) | -0.161 (p < .001) |
| ***Log(n):Right-skewed*** | 0.003 (p = .087) | 0.008 (p = .010) | -0.000 (p = 1.000) | 0.000 (p = 1.000) | -0.001 (p = .686) | 0.008 (p = .242) | 0.012 (p = .187) |

*Table S1. Linear mixed model results for evaluation metrics under age-skewed sampling conditions using normative models pre-trained on the UKB and adapted to the OASIS-3 dataset. Models assess the influence of standardized and log-transformed sample size (n) and age sampling strategy (Representative, Left-skewed, Right-skewed) on model performance metrics (MSLL, SMSE, EV, Rho, ICC, the lower and upper tail HC%). All variables were standardized to allow comparison of effect sizes. Representative sampling serves as the reference level. Reported β coefficients and corresponding p-values indicate the direction and significance of each effect.*

| | ***MSLL* β (p-value)** | ***SMSE* β (p-value)** | ***EV* β (p-value)** | ***Rho* β (p-value)** | ***ICC* β (p-value)** | ***Lower tail HC %* β (p-value)** | ***Upper tail HC %* β (p-value)** |
|---|---|---|---|---|---|---|---|
| ***Intercept (1F1M)*** | -0.195 (p < .001) | -0.567 (p < .001) | 0.644 (p < .001) | 0.701 (p < .001) | 0.504 (p < .001) | -0.219 (p < .001) | -0.373 (p < .001) |
| ***Log(n)*** | -0.054 (p < .001) | -0.102 (p < .001) | 0.000 (p = 1.000) | 0.000 (p = 1.000) | 0.245 (p < .001) | -0.228 (p < .001) | -0.210 (p < .001) |
| ***10F1M*** | -0.011 (p < .001) | -0.005 (p = .002) | -0.000 (p = 1.000) | -0.000 (p = 1.000) | -0.000 (p = .919) | 0.088 (p < .001) | 0.059 (p < .001) |
| ***4F1M*** | -0.009 (p < .001) | -0.006 (p < .001) | 0.000 (p = 1.000) | 0.000 (p = 1.000) | -0.000 (p = .995) | 0.050 (p < .001) | 0.036 (p < .001) |
| ***1F4M*** | 0.002 (p = .031) | 0.013 (p < .001) | 0.000 (p = 1.000) | 0.000 (p = 1.000) | -0.002 (p = .595) | -0.006 (p = .266) | 0.036 (p < .001) |
| ***1F10M*** | -0.001 (p = .593) | 0.020 (p < .001) | -0.000 (p = 1.000) | -0.000 (p = 1.000) | -0.005 (p = .138) | 0.011 (p = .032) | 0.036 (p < .001) |
| ***Log(n):10F1M*** | 0.004 (p < .001) | 0.006 (p < .001) | -0.000 (p = 1.000) | -0.000 (p = 1.000) | -0.004 (p = .280) | 0.006 (p = .267) | 0.001 (p = .855) |
| ***Log(n):4F1M*** | 0.004 (p < .001) | 0.004 (p = .014) | -0.000 (p = 1.000) | -0.000 (p = 1.000) | -0.003 (p = .366) | 0.008 (p = .133) | 0.000 (p = .970) |
| ***Log(n):1F4M*** | 0.000 (p = .975) | -0.001 (p = .479) | -0.000 (p = 1.000) | 0.000 (p = 1.000) | 0.001 (p = .825) | 0.007 (p = .208) | -0.046 (p < .001) |
| ***Log(n):1F10M*** | 0.006 (p < .001) | -0.002 (p = .175) | -0.000 (p = 1.000) | -0.000 (p = 1.000) | 0.004 (p = .192) | 0.001 (p = .793) | 0.015 (p = .001) |

*Table S2. Linear mixed model results for evaluation metrics under sex-imbalanced sampling conditions using normative models pre-trained on the UKB and adapted to the OASIS-3 dataset. Models assess the influence of standardized and log-transformed sample size (n) and sex ratio in the training set (1F:1M, 1F:4M, 1F:10M, 4F:1M, 10F:1M; F = female, M = male) on model performance metrics (MSLL, SMSE, EV, Rho, ICC, the lower and upper tail HC%). All variables were standardized to allow comparison of effect sizes. Representative sampling (1F:1M) serves as the reference level. Reported β coefficients and corresponding p-values indicate the direction and significance of each effect.*

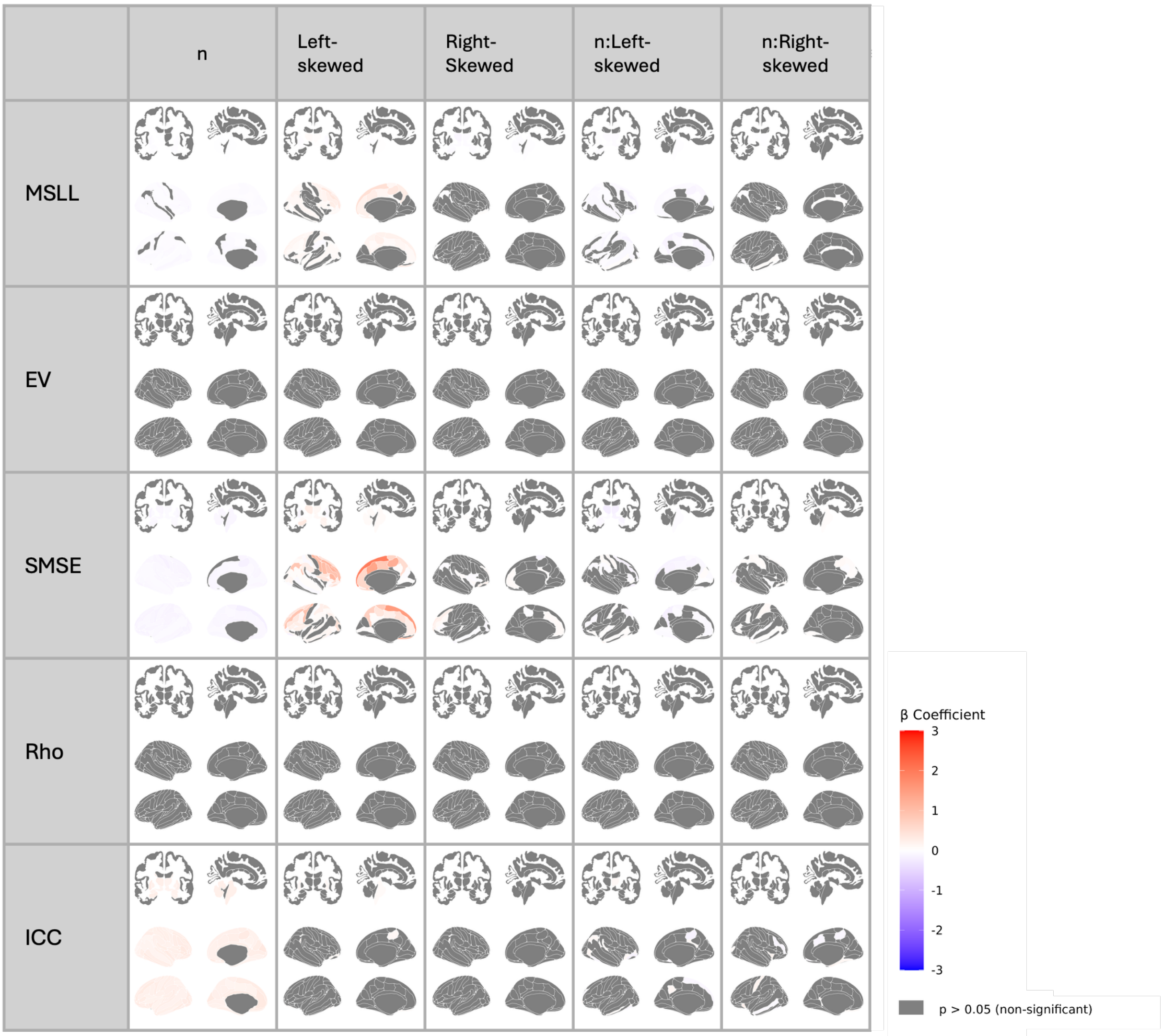

*Figure S12. Regional linear mixed-effects results for model-fit metrics in models pre-fitted in the UKB and adapted to OASIS-3 with age-skewed samples. Standardized Betas (β) for each performance metric (MSLL, SMSE, EV, Rho, ICC; rows) are regressed on log-standardized sample size (n), age-distribution contrasts (left-skewed, right-skewed; representative = reference), and their interactions. Sex ratio is balanced (1:1). Grey shading denotes regions that did not survive FDR correction (p ≥ 0.05).*

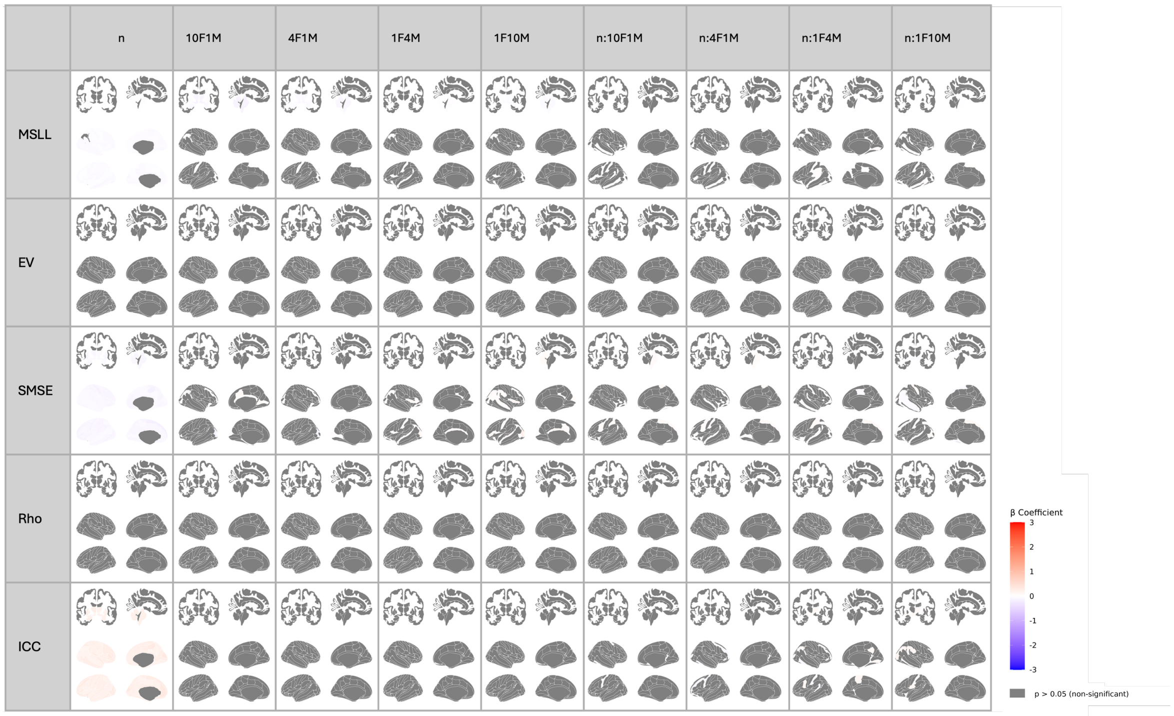

*Figure S13. Regional linear mixed-effects results for model-fit metrics in in models pre-fitted in the UKB and adapted to OASIS-3 with sex-imbalanced samples. Standardized Betas (β) for each performance metric (MSLL, SMSE, EV, Rho, ICC; rows) are regressed on log-standardized sample size (n), sex-ratio contrasts (female-to-male 10:1, 4:1, 1:1 [reference], 1:4, 1:10), and their interactions. Age distribution is representative. Grey shading denotes regions that did not survive FDR correction (p ≥ 0.05).*

#### 2.2. Z-scores errors

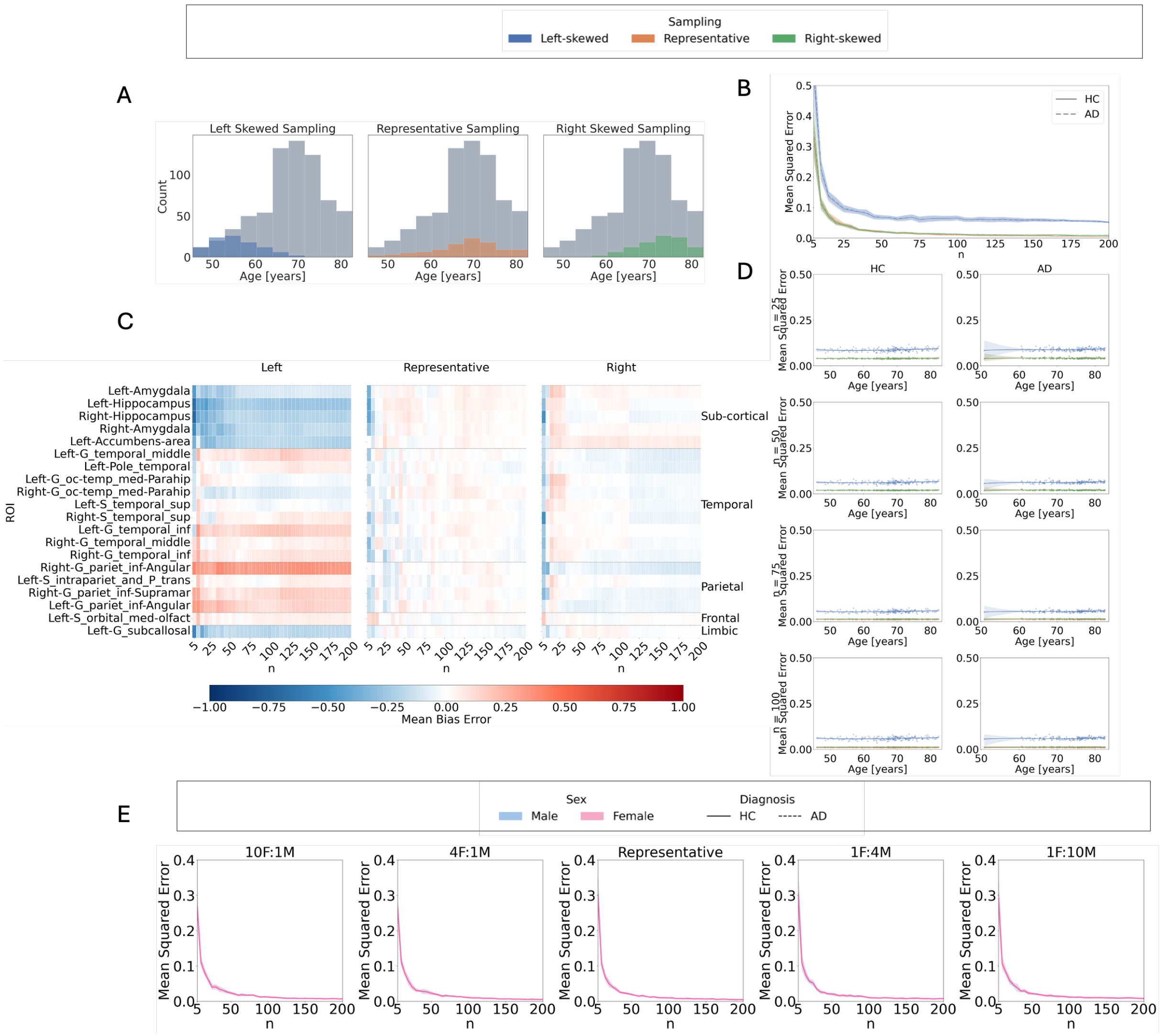

*Figure S14. Figure 5. Z-score errors in OASIS-3 using models trained in UKB and adapted to OASIS-3: Influence of sample size, age distribution, and sex imbalance on normative model outcomes. A: Age distributions for left-skewed (younger-biased), representative, and right-skewed (older-biased) sampling strategies, compared to the full training set in the OASIS-3 dataset. B: Mean squared error (MSE) of Z-scores relative to the full training set model across sample sizes, age distributions, and diagnostic groups. C: Mean bias error (MBE) per region across sample sizes and age distributions in the test set Shown are the 20 brain regions with the highness Cohen's d effect sizes based on models using the full training set. From left to right: results for left-skewed, representative, and right-skewed training sets. Blue indicates negative MBE (underestimation), red indicates positive MBE (overestimation), and white indicates close alignment with the full training set model. D: Cubic regression of MSE as a function of age, across sample sizes and sampling strategies. Left-skewed sampling shows increased errors in older individuals; right-skewed sampling shows increased errors in younger individuals. E: Centile curves for the Left Hippocampus as a function of age (females only), derived from models trained on left-skewed, representative, and right-skewed sampling (n = 100). Colored lines represent the 5th, 50th, and 95th percentiles; grey lines show centiles from the full training set model. F: MSE across sample sizes and test set sex, obtained using sex-imbalanced training sets (female-to-male ratios: 10:1, 4:1, 1:1, 1:4, 1:10), all with representative age distributions.*

| | ***MSE*** <br> ***β (p-value)*** | ***tOC*** <br> ***β (p-value)*** |
|---|---|---|
| ***Intercept (HC, Representative)*** | -0.313 (p < .001) | -0.462 (p < .001) |
| ***Log(n)*** | -0.451 (p < .001) | 0.026 (p = 0.015) |
| ***AD*** | 0.070 (p = 0.243) | 0.909 (p < .001) |
| ***Left-skewed*** | 0.930 (p < .001) | 0.281 (p < .001) |
| ***Right-skewed*** | 0.560 (p < .001) | -0.068 (p < .001) |
| ***Log(n):AD*** | -0.073 (p = 0.039) | 0.144 (p < .001) |
| ***Log(n):Left-skewed*** | -0.359 (p < .001) | -0.142 (p < .001) |
| ***Log(n):Right-skewed*** | -0.521 (p < .001) | 0.012 (p = 0.427) |
| ***Left-skewed:AD*** | 0.211 (p < .001) | 0.670 (p < .001) |
| ***Right-skewed:AD*** | -0.061 (p = 0.224) | -0.192 (p < .001) |
| ***Log(n):Left-skewed:AD*** | -0.030 (p = 0.555) | -0.187 (p < .001) |
| ***Log(n):Right-skewed:AD*** | 0.056 (p = 0.264) | 0.005 (p = 0.819) |

*Table S3. Linear mixed model results for MSE and total outlier count (tOC) under age-skewed sampling for models adapted from the UK Biobank to OASIS-3. Models evaluate the influence of diagnosis (HC, AD), log-transformed and standardized sample size (n), and age sampling strategy (Representative, Left-skewed, Right-skewed) on standardized deviation score outcomes. Continuous variables were standardized to allow comparison of effect sizes. Representative sampling and HC are used as reference levels. Reported β coefficients and corresponding p-values indicate the direction and significance of the effects.*

| | ***MSE*** <br> ***β (p-value)*** | ***tOC*** <br> ***β (p-value)*** |
|---|---|---|
| ***Intercept (HC, 1F 1M)*** | NA | -0.443 (p < .001) |
| ***Log(n)*** | 0.010 (p < .001) | 0.029 (p < .001) |
| ***AD*** | 0.003 (p = 0.168) | 0.016 (p < .001) |
| ***10F1M*** | 0.010 (p < .001) | -0.003 (p = 0.202) |
| ***4F1M*** | 0.014 (p < .001) | 0.003 (p = 0.119) |
| ***1F4M*** | NA | 0.576 (p < .001) |
| ***1F10M*** | 0.002 (p = 0.576) | 0.055 (p < .001) |
| ***Log(n):AD*** | 0.001 (p = 0.809) | 0.033 (p < .001) |
| ***Log(n):10F1M*** | -0.000 (p = 0.964) | -0.012 (p < .001) |
| ***Log(n):4F1M*** | 0.001 (p = 0.701) | -0.007 (p = 0.011) |
| ***Log(n):1F4M*** | -0.142 (p < .001) | -0.075 (p < .001) |
| ***Log(n):1F10M*** | 0.004 (p = 0.052) | 0.003 (p = 0.185) |
| ***10F1M:AD*** | 0.005 (p = 0.010) | 0.003 (p = 0.100) |
| ***4F1M:AD*** | -0.003 (p = 0.106) | 0.004 (p = 0.060) |
| ***1F4M:AD*** | 0.001 (p = 0.628) | 0.002 (p = 0.404) |
| ***1F10M:AD*** | -0.005 (p = 0.014) | -0.044 (p < .001) |
| ***Log(n):10F1M:AD*** | -0.002 (p = 0.604) | 0.002 (p = 0.558) |
| ***Log(n):4F1M:AD*** | -0.001 (p = 0.749) | 0.003 (p = 0.260) |
| ***Log(n):1F4M:AD*** | 0.002 (p = 0.560) | 0.006 (p = 0.025) |
| ***Log(n):1F10M:AD*** | -0.001 (p = 0.817) | 0.002 (p = 0.514) |

*Table S4. Linear mixed model results for MSE and total outlier count (tOC) under sex-imbalanced sampling for models adapted from the UK Biobank to OASIS-3. Models evaluate the influence of diagnosis (HC, AD), log-transformed and standardized sample size (n), and sex ratio in the training set (1:1, 1F:4M, 1F:10M, 4F:1M, 10F:1M) on standardized deviation score outcomes. Continuous variables were standardized to allow comparison of effect sizes. The 1:1 ratio and HC are used as reference levels. Reported β coefficients and corresponding p-values indicate the direction and significance of the effects.*

#### 2.3. Clinical validation

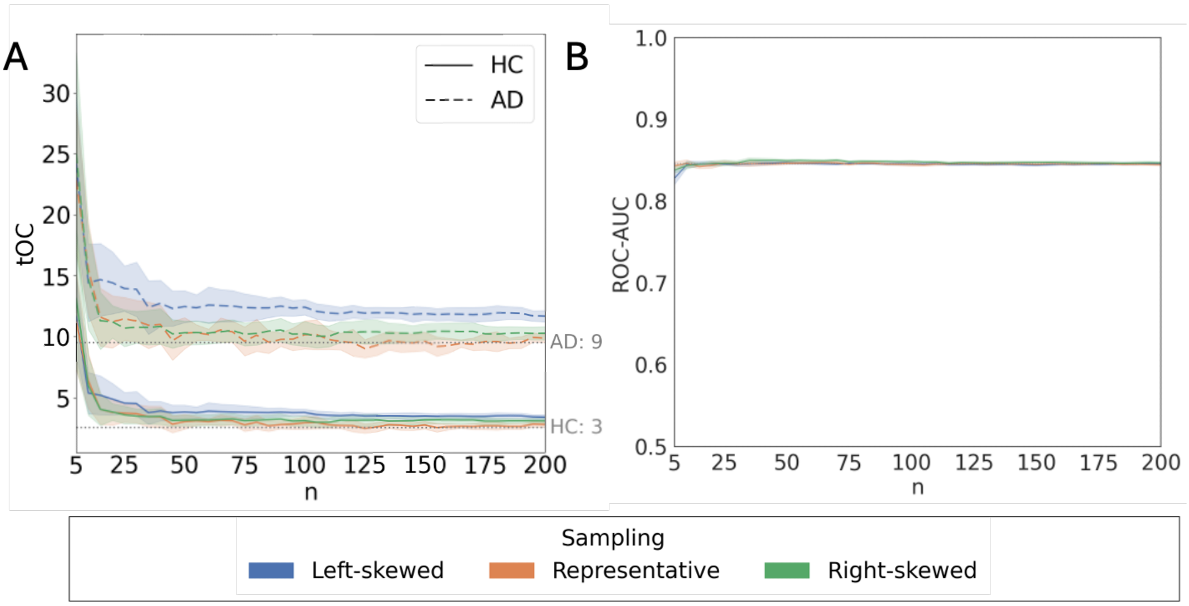

*Figure S15. Clinical validation for models pre-fitted in UKB and adapted to OASIS-3: effect of sample size and age distributions in outlier detection and classification performance. A: Average total Outlier Count (tOC) in the AD and HC are represented with dashed and solid lines respectively as a function of sample size (n). The shaded areas indicate the standard deviation across iterations. Dotted grey lines correspond to the estimations of tOC for HC and AD groups obtained with full sample size. B: ROC-AUC of Support Vector Classifier with 10-fold cross-validation as a function of sample size in the training set. The solid lines represent the average AUC across iterations for each sampling strategy. The dotted line shows the performance of the models trained with full sample size.*

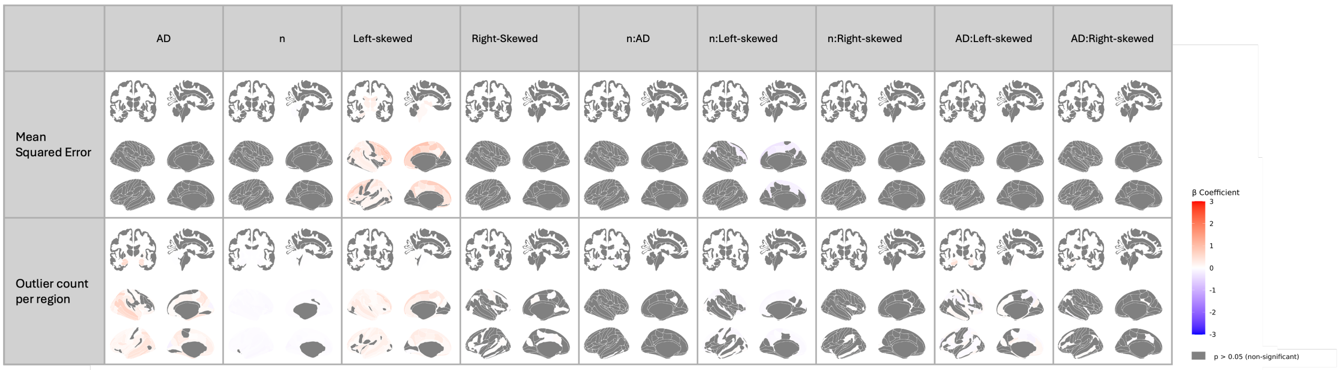

*Figure S16. Regional linear mixed-effects results for deviation-score metrics in models pre-fitted in the UKB and adapted to OASIS-3 with age-skewed samples. Standardized Betas (β) are shown for the mean-squared error of z-scores relative to the full reference model and for the count of extreme z-score outliers per region (rows). Predictors include Alzheimer's disease diagnosis (AD), log-standardized sample size (n), age-distribution contrasts (left-skewed, right-skewed; representative = reference), and their two-way interactions with n and AD. Sex ratio is balanced (1:1). Grey shading denotes regions that did not survive FDR correction (p ≥ 0.05).*

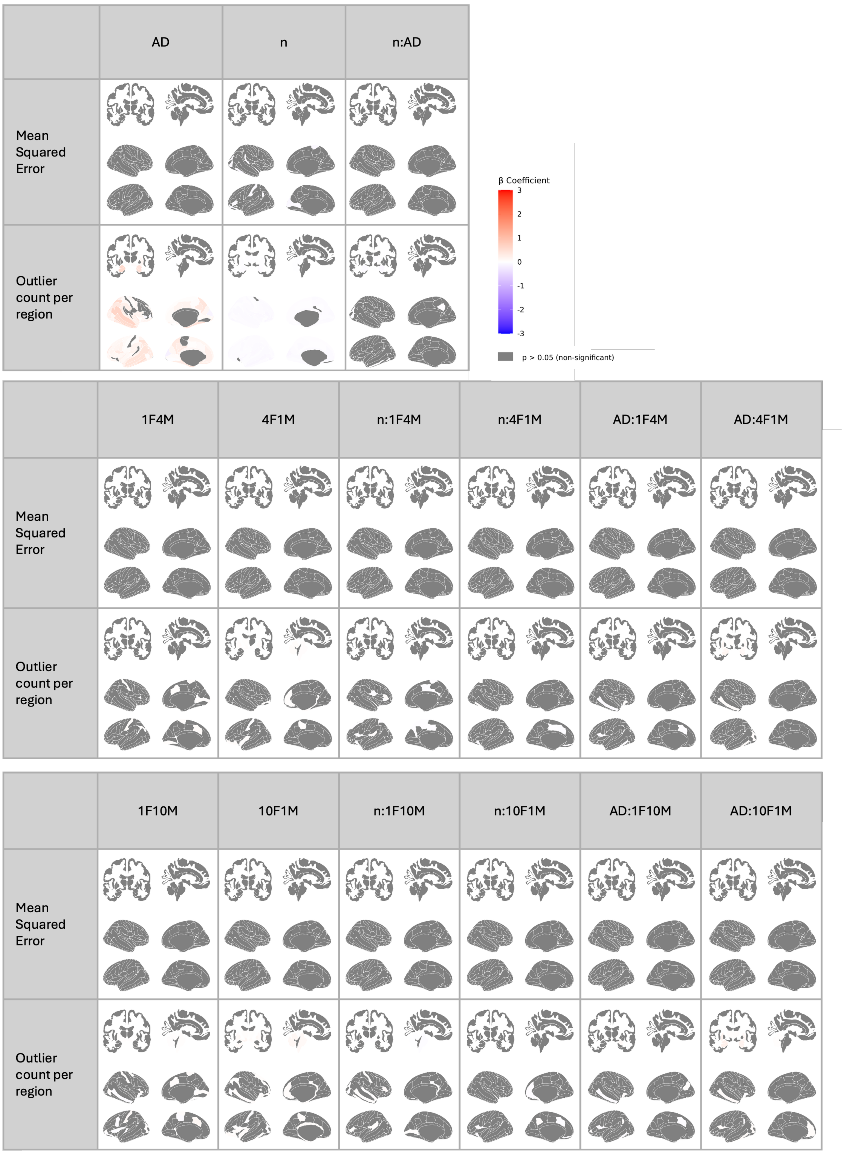

*Figure S17. Regional linear mixed-effects results for deviation-score metrics in models pre-fitted in the UKB and adapted to OASIS-3 with sex-imbalanced samples. Standardized Betas (β) are shown for the mean-squared error of z-scores relative to the full reference model and for the count of extreme z-score outliers per region (rows). Predictors include Alzheimer's disease diagnosis (AD), log-standardized sample size (n), sex-ratio contrasts (female-to-male 10:1, 4:1, 1:1 [reference], 1:4, 1:10), and their two-way interactions with n and AD. Age distribution is representative. Grey shading denotes regions that did not survive FDR correction (p ≥ 0.05).*

### 3. Replication of direct training in independent dataset (AIBL)

#### 3.1. Model fit evaluation

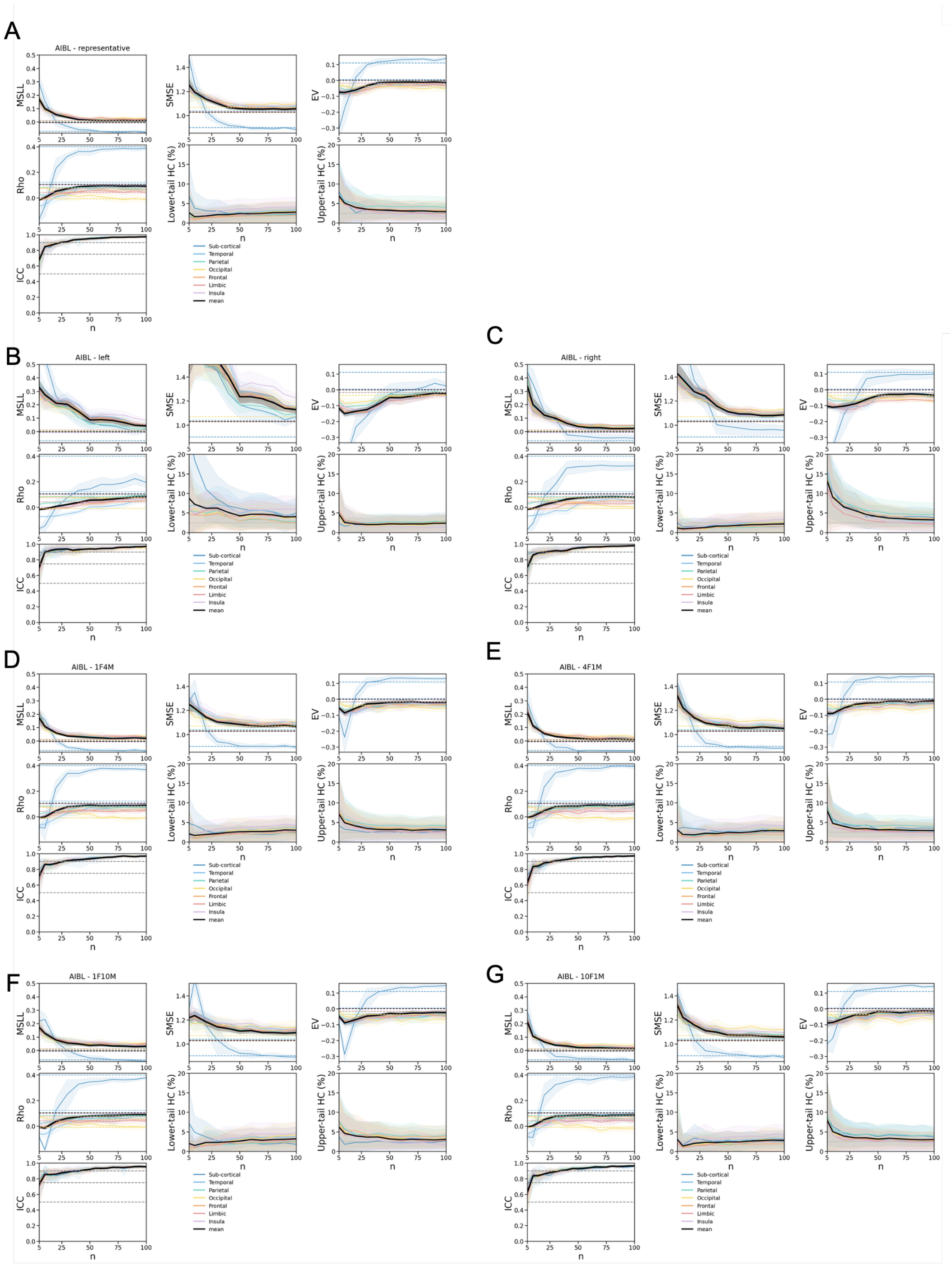

*Figure S18. Model fit evaluation of normative models trained within AIBL. Performance is assessed in Healthy Controls (HC) test set of AIBL for different sampling strategies of the training set, including (A) Representative, (B) Left-skewed, and (C) Right-skewed age distributions, as well as (D, E, F, G) sex-imbalanced adaptations with female-to-male ratios of 1:4, 4:1, 1:10, and 10:1, respectively. Model performance is assessed as a function of the adaptation set size (n) using the HC test set from the OASIS-3 dataset. Each panel presents the following evaluation metrics: Mean Standardized Log Loss (MSLL), Standardized Mean Squared Error (SMSE), Explained Variance (EV), Pearson correlation coefficient*

(Rho), Intraclass Correlation Coefficient (ICC), lower-tail HC percentage (below the 2.5% bound), upper-tail HC percentage (above the 97.5% bound). Solid lines indicate the mean performance across all regions within each lobe (as well as the mean of subcortical regions), while shaded areas represent the standard deviation across 10 iterations per sample size. In ICC plots, grey dashed lines denote commonly accepted reliability thresholds: <0.5 (poor), 0.5–0.75 (moderate), 0.75–0.9 (good), and >0.9 (excellent) and shaded areas indicate standard deviation across ROI.*

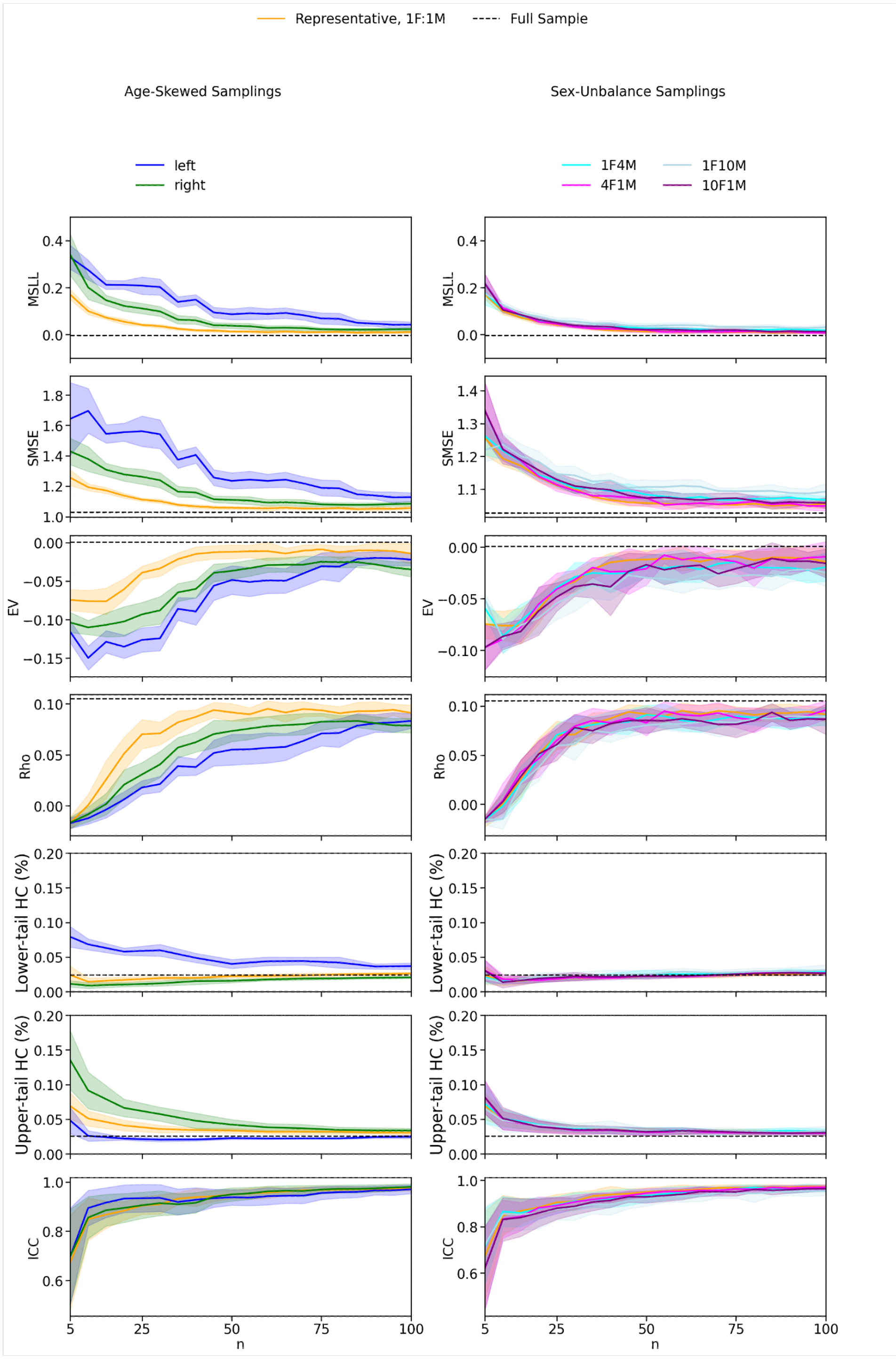

*Figure S19. Model fit evaluation of normative models trained on AIBL across different sampling strategies. Models were evaluated in the HC test set for various training sample sizes (n). Age-skewed sampling strategies include representative (matching the initial age distribution with balanced sex, 1F:1M), left-skewed (favoring younger*

*individuals), and right-skewed (favoring older individuals), each with balanced sex distributions. Sex-imbalanced sampling strategies include female-to-male ratios of 1:1.1, 1:4, 1:10, 4:1, and 10:1, all with representative age distributions. Solid lines represent the mean metric values across ROIs and iterations, with shaded areas indicating the standard deviation across iterations for MSLL, SMSE, EV, Rho, the lower and upper tail HC%. For ICC, shaded areas represent variation across ROIs, as ICC already reflects variability across iteration. Dashed lines indicate the mean metric values for models trained with the full sample for MSLL, SMSE, EV, Rho, and the lower and upper tail HC%.*

| | ***MSLL β (p-value)*** | ***EV β (p-value)*** | ***SMSE β (p-value)*** | ***Rho β (p-value)*** | ***ICC β (p-value)*** | ***Lower tail HC % β (p-value)*** | ***Upper tail HC % β (p-value)*** |
|---|---|---|---|---|---|---|---|
| ***Intercept (Representative)*** | -0.293 (p < .001) | -0.332 (p < .001) | 0.175 (p < .001) | 0.067 (p = .181) | 0.100 (p < .001) | -0.163 (p = .004) | -0.026 (p = .626) |
| ***Log(n)*** | -0.426 (p < .001) | -0.308 (p < .001) | 0.262 (p < .001) | 0.219 (p < .001) | 0.755 (p < .001) | 0.103 (p < .001) | -0.307 (p < .001) |
| ***Left-skewed*** | 1.080 (p < .001) | 1.382 (p < .001) | -0.517 (p < .001) | -0.212 (p < .001) | 0.043 (p = .011) | 1.096 (p < .001) | -0.457 (p < .001) |
| ***Right-skewed*** | 0.465 (p < .001) | 0.438 (p < .001) | -0.338 (p < .001) | -0.127 (p < .001) | -0.052 (p = .002) | -0.265 (p < .001) | 0.562 (p < .001) |
| ***Log(n):Left-skewed*** | -0.484 (p < .001) | -0.670 (p < .001) | 0.196 (p < .001) | 0.002 (p = .897) | -0.208 (p < .001) | -0.555 (p < .001) | 0.182 (p < .001) |
| ***Log(n):Right-skewed*** | -0.411 (p < .001) | -0.284 (p < .001) | 0.072 (p < .001) | 0.006 (p = .703) | 0.002 (p = .912) | 0.047 (p = .010) | -0.572 (p < .001) |

*Table S5. Linear mixed model results for evaluation metrics under age-skewed sampling conditions using normative models trained on AIBL dataset. Models assess the influence of standardized and log-transformed sample size (n) and age sampling strategy (Representative, Left-skewed, Right-skewed) on model performance metrics (MSLL, SMSE, EV, Rho, ICC, the lower and upper tail HC%). All variables were standardized to allow comparison of effect sizes. Representative sampling serves as the reference level. Reported β coefficients and corresponding p-values indicate the direction and significance of each effect.*

| | ***MSLL β (p-value)*** | ***SMSE β (p-value)*** | ***EV β (p-value)*** | ***Rho β (p-value)*** | ***ICC β (p-value)*** | ***Lower tail HC % β (p-value)*** | ***Upper tail HC % β (p-value)*** |
|---|---|---|---|---|---|---|---|
| ***Intercept (1F1M)*** | -0.294 (p < .001) | -0.333 (p < .001) | 0.177 (p = .001) | 0.067 (p = .285) | 0.100 (p < .001) | -0.163 (p < .001) | -0.026 (p = .635) |
| ***Log(n)*** | -0.423 (p < .001) | -0.306 (p < .001) | 0.258 (p < .001) | 0.219 (p < .001) | 0.755 (p < .001) | 0.103 (p < .001) | -0.307 (p < .001) |
| ***10F1M*** | 0.107 (p < .001) | 0.118 (p < .001) | -0.111 (p < .001) | -0.035 (p = .021) | -0.223 (p < .001) | 0.041 (p < .001) | 0.015 (p = .157) |
| ***4F1M*** | 0.056 (p < .001) | 0.054 (p < .001) | -0.042 (p = .004) | -0.010 (p = .490) | -0.113 (p < .001) | 0.048 (p < .001) | -0.005 (p = .651) |
| ***1F4M*** | 0.101 (p < .001) | 0.100 (p < .001) | -0.054 (p < .001) | -0.026 (p = .086) | -0.081 (p < .001) | 0.076 (p < .001) | 0.037 (p < .001) |
| ***1F10M*** | 0.248 (p < .001) | 0.234 (p < .001) | -0.164 (p < .001) | -0.062 (p < .001) | -0.272 (p < .001) | 0.141 (p < .001) | 0.033 (p = .002) |
| ***Log(n):10F1M*** | -0.056 (p < .001) | -0.069 (p < .001) | 0.029 (p = .048) | -0.022 (p = .146) | 0.095 (p < .001) | -0.040 (p < .001) | -0.067 (p < .001) |
| ***Log(n):4F1M*** | -0.076 (p < .001) | -0.077 (p < .001) | 0.037 (p = .011) | -0.013 (p = .400) | 0.105 (p < .001) | -0.059 (p < .001) | -0.074 (p < .001) |
| ***Log(n):1F4M*** | 0.041 (p < .001) | 0.026 (p = .005) | -0.073 (p < .001) | -0.011 (p = .461) | -0.069 (p < .001) | 0.040 (p < .001) | -0.007 (p = .501) |
| ***Log(n):1F10M*** | 0.036 (p = .001) | 0.060 (p < .001) | -0.076 (p < .001) | 0.002 (p = .914) | -0.112 (p < .001) | 0.080 (p < .001) | 0.018 (p = .083) |

*Table S6. Linear mixed model results for evaluation metrics under sex-imbalanced sampling conditions using normative models trained on AIBL dataset. Models assess the influence of standardized and log-transformed sample size (n) and sex ratio in the training set (1F:1M, 1F:4M, 1F:10M, 4F:1M, 10F:1M; F = female, M = male) on model performance metrics (MSLL, SMSE, EV, Rho, ICC, the lower and upper tail HC%). All variables were standardized to allow comparison of effect sizes. Representative sampling (1F:1M) serves as the reference level. Reported β coefficients and corresponding p-values indicate the direction and significance of each effect.*

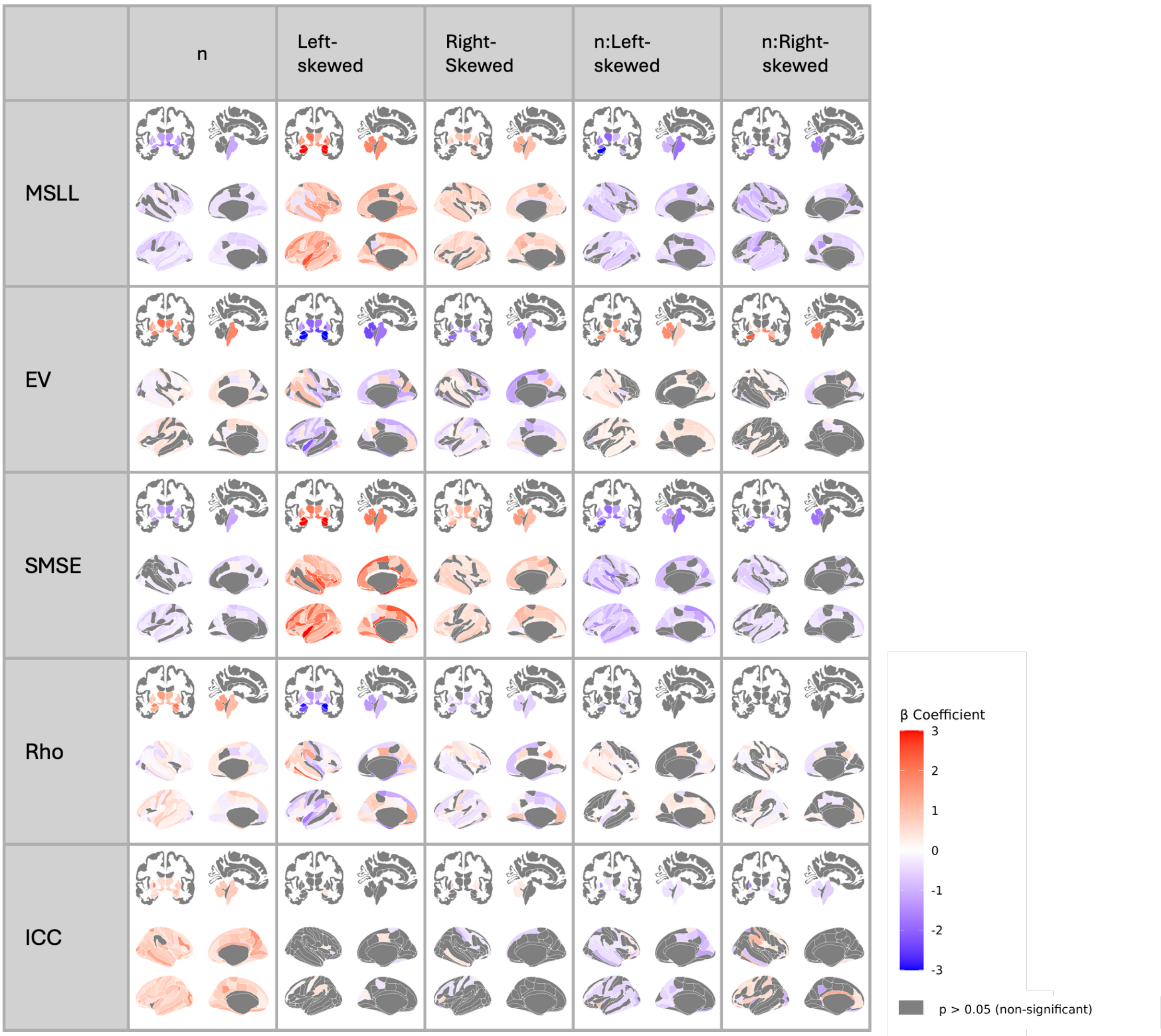

*Figure S20. Regional linear mixed-effects results for model-fit metrics in models trained in AIBL with age-skewed samples. Standardized Betas (β) for each performance metric (MSLL, SMSE, EV, Rho, ICC; rows) are regressed on log-standardized sample size (n), age-distribution contrasts (left-skewed, right-skewed; representative = reference), and their interactions. Sex ratio is balanced (1:1). Grey shading denotes regions that did not survive FDR correction (p ≥ 0.05).*

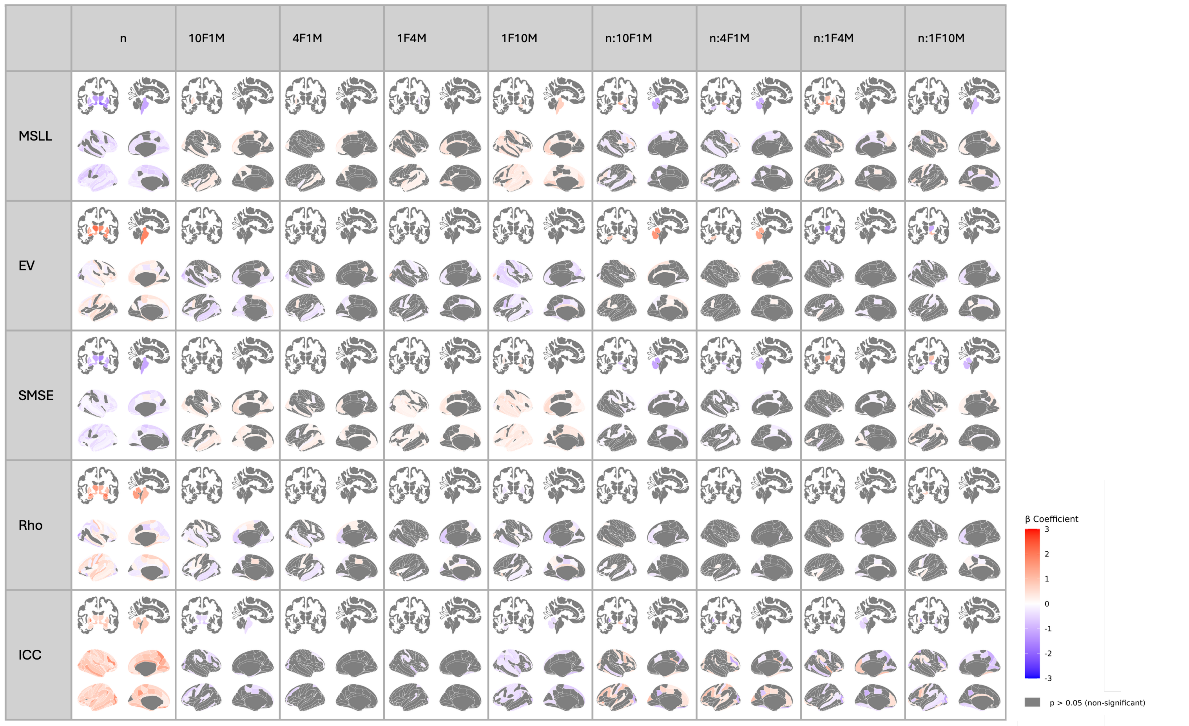

*Figure S21. Regional linear mixed-effects results for model-fit metrics in models trained in AIBL with sex-imbalanced samples. Standardized Betas (β) for each performance metric (MSLL, SMSE, EV, Rho, ICC; rows) are regressed on log-standardized sample size (n), sex-ratio contrasts (female-to-male 10:1, 4:1, 1:1 [reference], 1:4, 1:10), and their interactions. Age distribution is representative. Grey shading denotes regions that did not survive FDR correction (p ≥ 0.05).*

#### 3.2. Z-scores errors

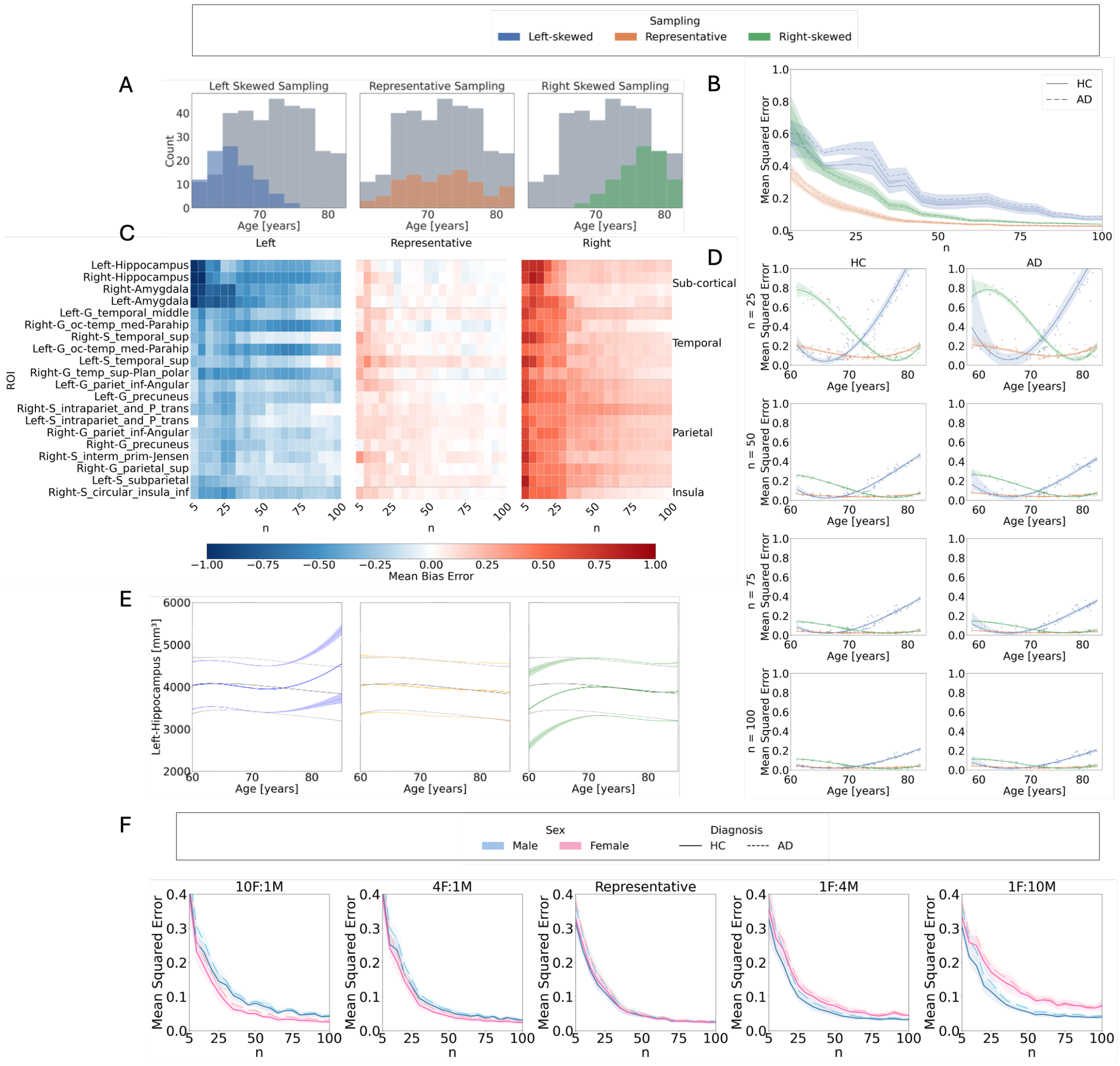

*Figure S22. Z-score errors in AIBL using models trained directly on the dataset: Influence of sample size, age distribution, and sex imbalance on normative model outcomes. A: Age distributions for left-skewed (younger-biased), representative, and right-skewed (older-biased) sampling strategies, compared to the full training set in the OASIS-3 dataset. B: Mean squared error (MSE) of Z-scores relative to the full training set model across sample sizes, age distributions, and diagnostic groups. C: Mean bias error (MBE) per region across sample sizes and age distributions in the test set Shown are the 20 brain regions with the highness Cohen's d effect sizes based on models using the full training set. From left to right: results for left-skewed, representative, and right-skewed training sets. Blue indicates negative MBE (underestimation), red indicates positive MBE (overestimation), and white indicates close alignment with the full training set model. D: Cubic regression of MSE as a function of age, across sample sizes and sampling strategies. Left-skewed sampling shows increased errors in older individuals; right-skewed sampling shows increased errors in younger individuals. E: Centile curves for the Left Hippocampus as a function of age (females only), derived from models trained on left-skewed, representative, and right-skewed sampling (n = 100). Colored lines represent the 5th, 50th, and 95th percentiles; grey lines show centiles from the full training set model. F: MSE across sample sizes and test set sex, obtained using sex-imbalanced training sets (female-to-male ratios: 10:1, 4:1, 1:1, 1:4, 1:10), all with representative age distributions.*

| | MSE<br>β (p-value) | tOC<br>β (p-value) |
|---|---|---|
| ***Intercept (HC, Representative)*** | -0.313 (p < .001) | -0.462 (p < .001) |
| ***Log(n)*** | -0.451 (p < .001) | 0.026 (p = 0.015) |
| ***AD*** | 0.070 (p = 0.243) | 0.909 (p < .001) |
| ***Left-skewed*** | 0.930 (p < .001) | 0.281 (p < .001) |
| ***Right-skewed*** | 0.560 (p < .001) | -0.068 (p < .001) |
| ***Log(n):AD*** | -0.073 (p = 0.039) | 0.144 (p < .001) |
| ***Log(n):Left-skewed*** | -0.359 (p < .001) | -0.142 (p < .001) |
| ***Log(n):Right-skewed*** | -0.521 (p < .001) | 0.012 (p = 0.427) |
| ***Left-skewed:AD*** | 0.211 (p < .001) | 0.670 (p < .001) |
| ***Right-skewed:AD*** | -0.061 (p = 0.224) | -0.192 (p < .001) |
| ***Log(n):Left-skewed:AD*** | -0.030 (p = 0.555) | -0.187 (p < .001) |
| ***Log(n):Right-skewed:AD*** | 0.056 (p = 0.264) | 0.005 (p = 0.819) |

*Table S7. Linear mixed model results for MSE and total outlier count (tOC) under age-skewed sampling for models trained in AIBL. Models evaluate the influence of diagnosis (HC, AD), log-transformed and standardized sample size (n), and age sampling strategy (Representative, Left-skewed, Right-skewed) on standardized deviation score outcomes. Continuous variables were standardized to allow comparison of effect sizes. Representative sampling and HC are used as reference levels. Reported β coefficients and corresponding p-values indicate the direction and significance of the effects.*

| | MSE<br>β (p-value) | tOC<br>β (p-value) |
|---|---|---|
| ***Intercept (HC, 1F 1M)*** | 0.052 (p < .001) | -0.462 (p < .001) |
| ***Log(n)*** | 0.093 (p < .001) | 0.011 (p = 0.207) |
| ***AD*** | 0.183 (p < .001) | 0.012 (p = 0.141) |
| ***10F1M*** | 0.070 (p = 0.009) | 0.020 (p = 0.020) |
| ***4F1M*** | 0.011 (p = 0.404) | 0.036 (p < .001) |
| ***1F4M*** | 0.007 (p = 0.614) | 0.909 (p < .001) |
| ***1F10M*** | 0.002 (p = 0.896) | 0.019 (p = 0.148) |
| ***Log(n):AD*** | 0.003 (p = 0.823) | 0.031 (p = 0.017) |
| ***Log(n):10F1M*** | -0.451 (p < .001) | 0.012 (p = 0.346) |
| ***Log(n):4F1M*** | -0.055 (p < .001) | 0.006 (p = 0.667) |
| ***Log(n):1F4M*** | -0.070 (p < .001) | 0.026 (p < .001) |
| ***Log(n):1F10M*** | -0.010 (p = 0.264) | -0.010 (p = 0.221) |
| ***10F1M:AD*** | 0.042 (p < .001) | -0.015 (p = 0.073) |
| ***4F1M:AD*** | -0.073 (p < .001) | 0.010 (p = 0.225) |
| ***1F4M:AD*** | -0.000 (p = 0.980) | 0.021 (p = 0.014) |
| ***1F10M:AD*** | -0.004 (p = 0.771) | 0.144 (p < .001) |
| ***Log(n):10F1M:AD*** | 0.001 (p = 0.949) | -0.005 (p = 0.719) |
| ***Log(n):4F1M:AD*** | 0.004 (p = 0.765) | -0.017 (p = 0.188) |
| ***Log(n):1F4M:AD*** | 0.052 (p < .001) | -0.006 (p = 0.657) |
| ***Log(n):1F10M:AD*** | 0.093 (p < .001) | 0.021 (p = 0.101) |

*Table S8. Linear mixed model results for MSE and total outlier count (tOC) under sex-imbalanced sampling for model trained in AIBL. Models evaluate the influence of diagnosis (HC, AD), log-transformed and standardized sample size (n), and sex ratio in the training set (1:1, 1F:4M, 1F:10M, 4F:1M, 10F:1M) on standardized deviation score outcomes. Continuous variables were standardized to allow comparison of effect sizes. The 1:1 ratio and HC are used as reference levels. Reported β coefficients and corresponding p-values indicate the direction and significance of the effects.*

#### 3.3. Clinical validation

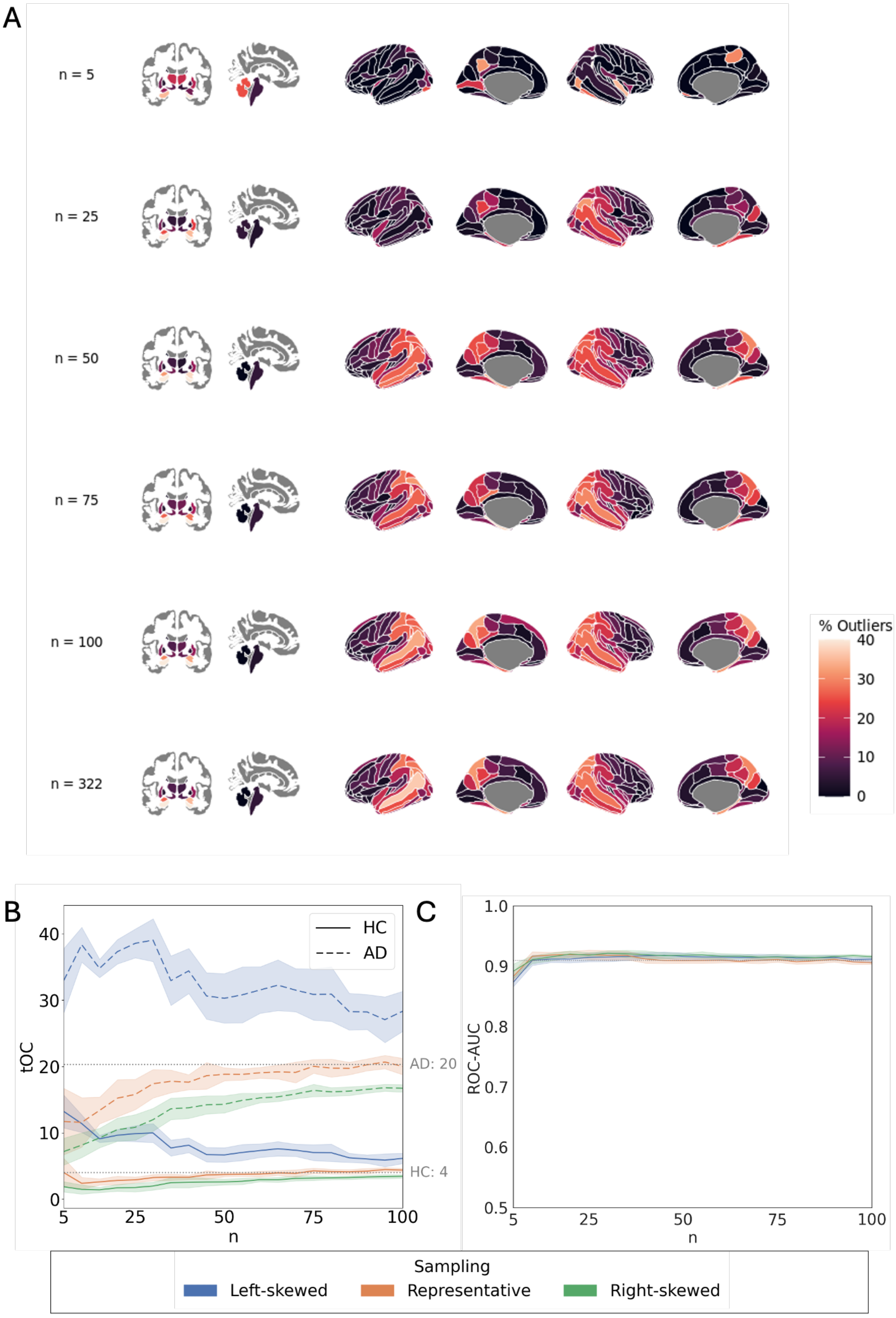

*Figure S23. Clinical validation for models trained in AIBL: effect of sample size and age distributions in outlier detection and classification performance. A: Percentage of participants with extreme negative deviation (Z < -1.96) in each brain region in the AD group, shown for independent example iterations at different sample sizes with representative age distribution. B: Average total Outlier Count (tOC) in the AD and HC are represented with dashed and solid lines respectively as a function of sample size (n). The shaded areas indicate the standard deviation across iterations. Dotted grey lines correspond to the estimations of tOC for HC and AD groups obtained with full sample size. C: ROC-AUC of Support Vector Classifier with 10-fold cross-validation as a function of sample size in the training set. The solid lines represent the average AUC across iterations for each sampling strategy. The dotted line shows the performance of the models trained with full sample size.*

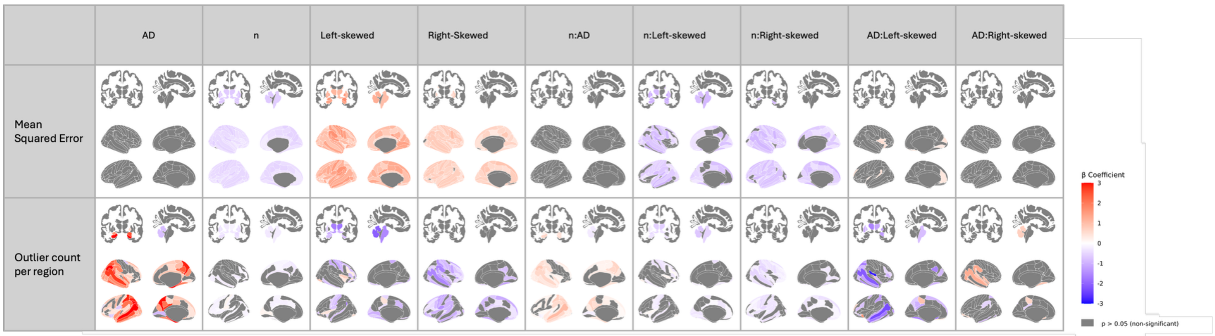

*Figure S24. Regional linear mixed-effects results for deviation-score metrics in models trained in AIBL with age-skewed samples. Standardized Betas (β) are shown for the mean-squared error of z-scores relative to the full reference model and for the count of extreme z-score outliers per region (rows). Predictors include Alzheimer's disease diagnosis (AD), log-standardized sample size (n), age-distribution contrasts (left-skewed, right-skewed; representative = reference), and their two-way interactions with n and AD. Sex ratio is balanced (1:1). Grey shading denotes regions that did not survive FDR correction (p ≥ 0.05).*

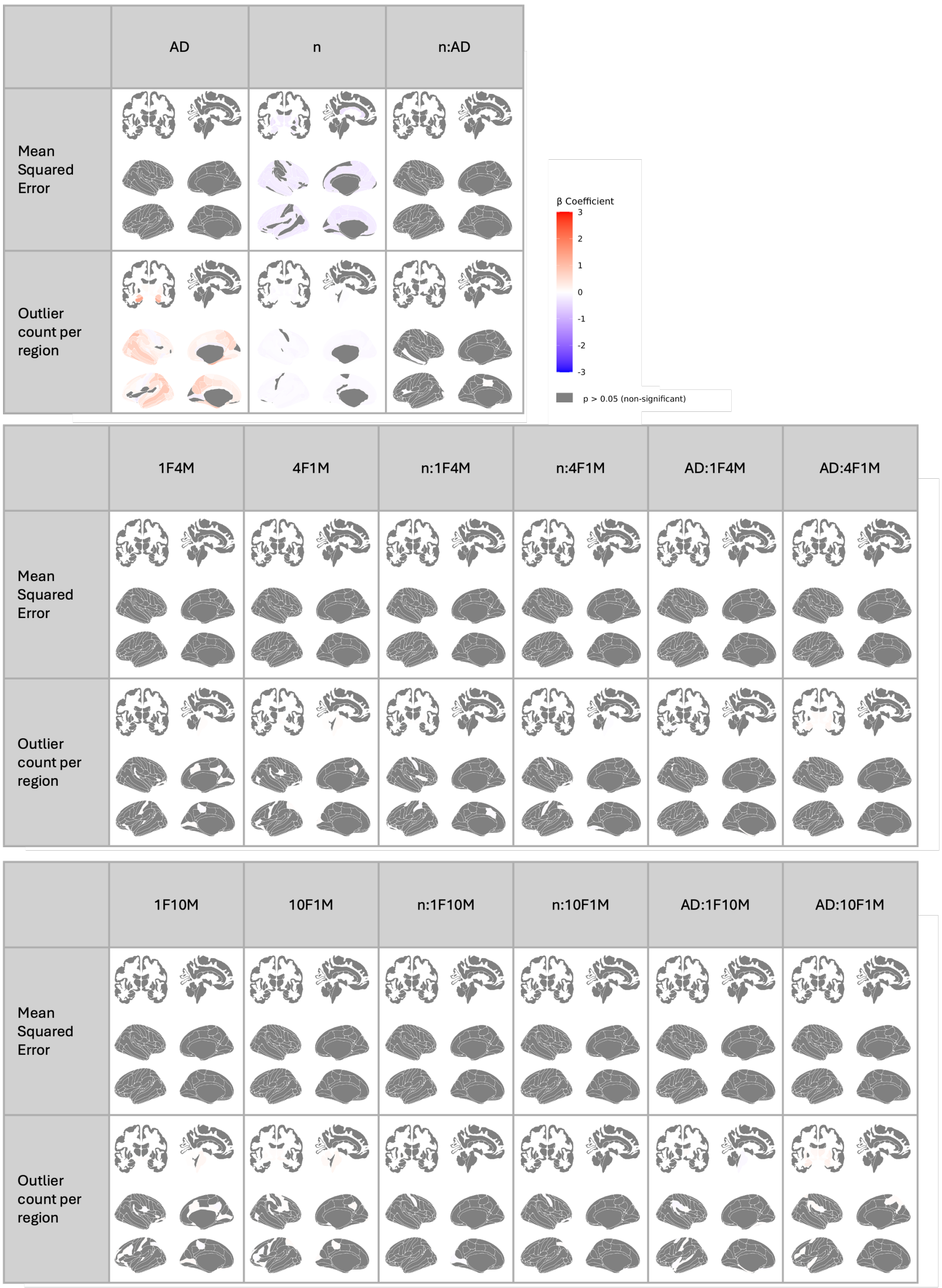

*Figure S25. Regional linear mixed-effects results for deviation-score metrics in models trained in AIBL with sex-imbalanced samples. Standardized Betas (β) are shown for the mean-squared error of z-scores relative to the full reference model and for the count of extreme z-score outliers per region (rows). Predictors include Alzheimer's disease diagnosis (AD), log-standardized sample size (n), sex-ratio contrasts (female-to-male 10:1, 4:1, 1:1 [reference], 1:4, 1:10), and their two-way interactions with n and AD. Age distribution is representative. Grey shading denotes regions that did not survive FDR correction (p ≥ 0.05).*

#### 3.4. Adaptation from large dataset

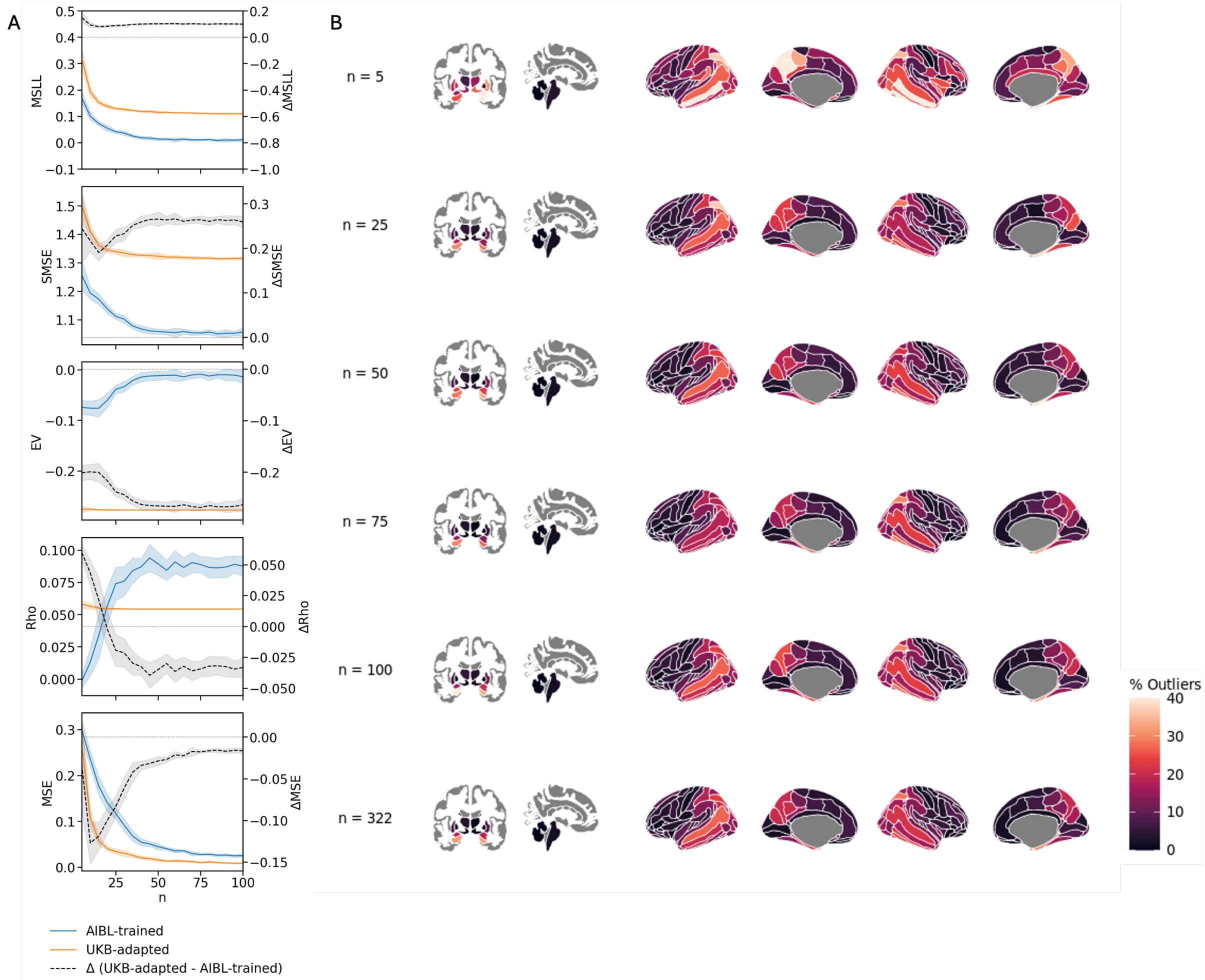

*Figure S26. Adaptive transfer learning evaluation in AIBL. A. Comparison of model performance between direct training models (AIBL trained) and models pretrained on UKB then adapted to AIBL (UKB-adapted). The grey curve shows the difference in performance (within-cohort minus adapted), plotted on a separate y-axis (right). For MSLL, SMSE, and MSE, lower values indicate better performance; therefore, positive differences indicate better performance of the within-cohort models. For EV and Rho, higher values indicate better performance; hence, positive differences reflect better performance of the adapted models. EV and Rho values remain constant across models, as the adaptation procedure only modifies mean of models and does not affect the shape. B. Example adaptation iteration showing the percentage of individuals identified as outliers per region. Last row shows the outlier detection obtained using the full adaptation sample (n = 692).*

#### 3.5. Additional results

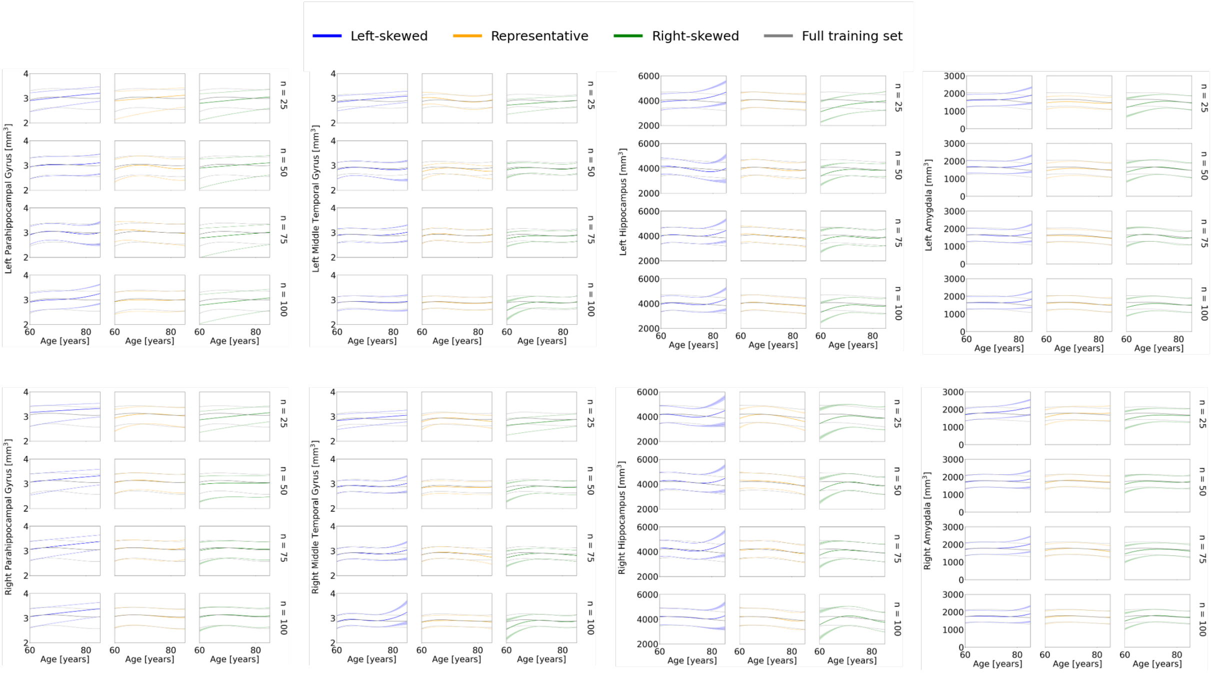

*Figure S27. Centile curve overlays for selected cortical and subcortical regions across sampling strategies and sample sizes. For each ROI, models trained using representative, left-skewed, and right-skewed sampling are shown for multiple sample sizes. Colored lines depict the 5th, 50th, and 95th percentiles estimated from each model, while grey lines indicate the corresponding centiles from the full training set. These overlays highlight how centile estimates diverge when age coverage is limited, particularly at the extreme of age ranges.*

### 4. Replication with Adaptive Transfer Learning in independent dataset (AIBL)

#### 4.1. Model fit evaluation

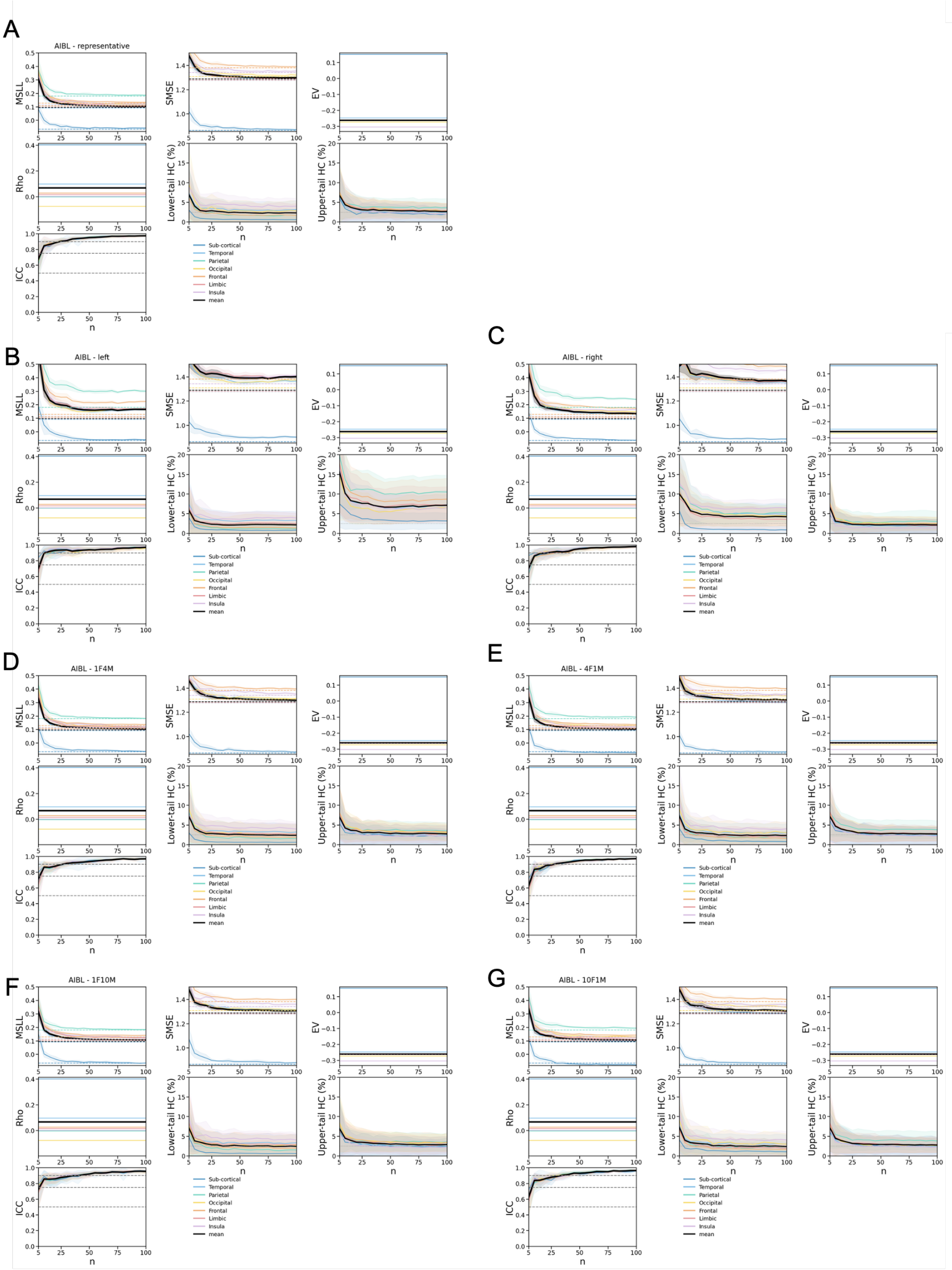

*Figure S28.Model fit evaluation of normative models pre-trained on the UKB and adapted to AIBL. Performance is assessed in Healthy Controls (HC) test set of AIBL dataset for different sampling strategies of the adaptation set, including (A) Representative, (B) Left-skewed, and (C) Right-skewed age distributions, as well as (D, E, F, G) sex-imbalanced adaptations with female-to-male ratios of 1:4, 4:1, 1:10, and 10:1, respectively. Model performance is assessed as a function of the adaptation set size (n) using the HC test set from the OASIS-3 dataset. Each panel presents the following evaluation metrics: Mean Standardized Log Loss (MSLL), Standardized Mean Squared Error (SMSE), Explained Variance (EV), Pearson correlation coefficient (Rho), Intraclass Correlation Coefficient (ICC), lower-tail HC percentage (below the 2.5% bound), upper-tail HC percentage (above the 97.5% bound). Solid lines indicate the mean*

*performance across all regions within each lobe (as well as the mean of subcortical regions), while shaded areas represent the standard deviation across 10 iterations per sample size. In ICC plots, grey dashed lines denote commonly accepted reliability thresholds: <0.5 (poor), 0.5–0.75 (moderate), 0.75–0.9 (good), and >0.9 (excellent) and shaded areas indicate standard deviation across ROI.*

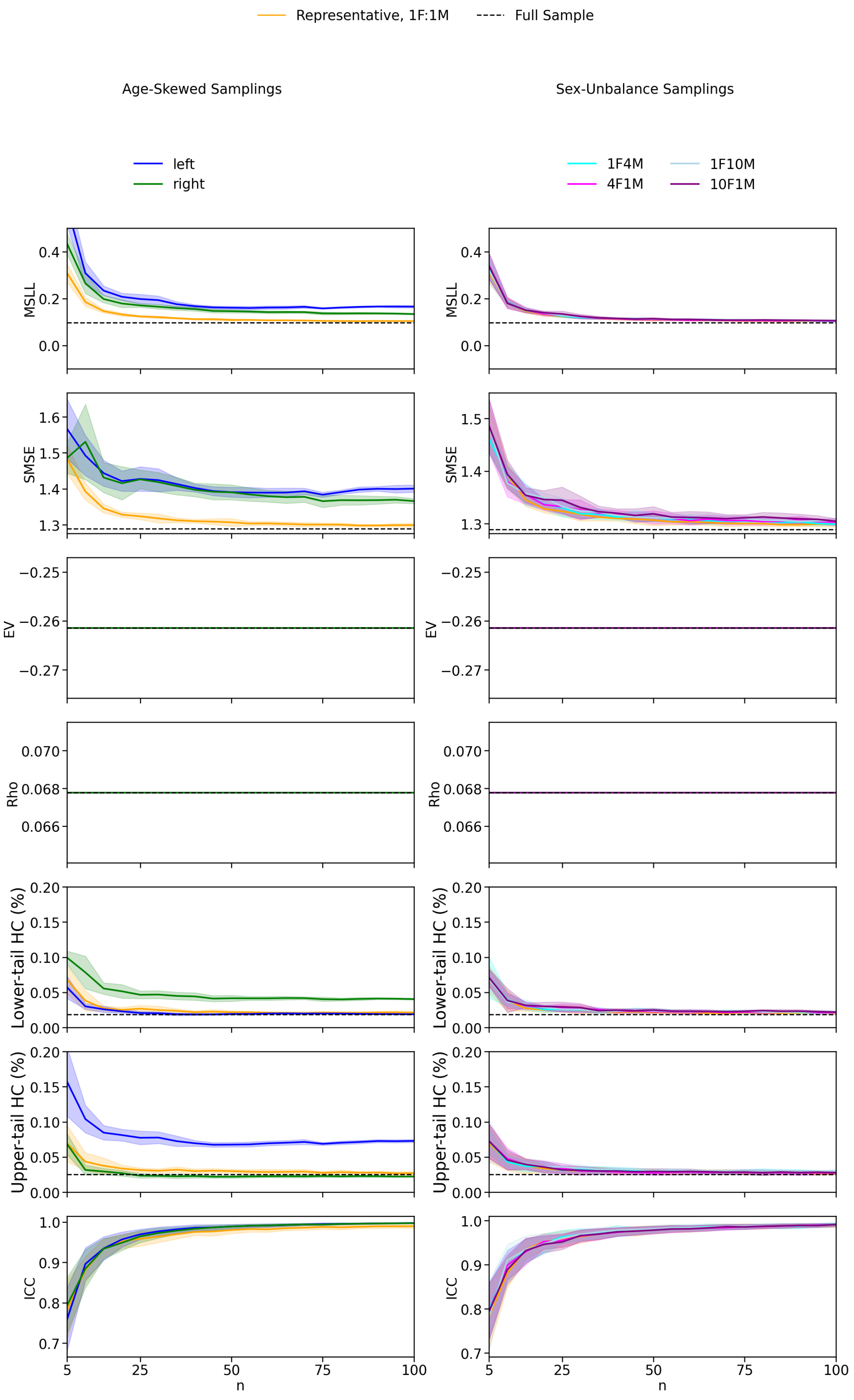

*Figure S29. Model fit evaluation of normative models pre-trained on the UKB and adapted to AIBL across different sampling strategies. Models were evaluated in the HC test set for various adaptation sample sizes (n). Age-skewed sampling strategies include representative (matching the initial age distribution with balanced sex, 1F:1M), left-skewed (favoring younger individuals), and right-skewed (favoring older individuals), each with balanced sex distributions. Sex-imbalanced sampling strategies include female-to-male ratios of 1:1.1, 1:4, 1:10, 4:1, and 10:1, all with representative age distributions. Solid lines represent the mean metric values across ROIs and iterations, with shaded areas indicating the standard deviation across iterations for MSLL, SMSE, EV, Rho, the lower and upper tail HC%). For ICC, shaded areas represent variation across ROIs, as ICC already reflects variability across iteration. Dashed lines indicate the mean metric values for models trained with the full sample for MSLL, SMSE, EV, and Rho.*

| | ***MSLL* β (p-value)** | ***SMSE* β (p-value)** | ***EV* β (p-value)** | ***Rho* β (p-value)** | ***ICC* β (p-value)** | ***Lower tail HC %* β (p-value)** | ***Upper tail HC %* β (p-value)** |
|---|---|---|---|---|---|---|---|
| ***Intercept (Representative)*** | 0.792 (p < .001) | 1.040 (p < .001) | -2.562 (p < .001) | -0.076 (p < .001) | 0.502 (p < .001) | -0.010 (p = .869) | -0.168 (p = .009) |
| ***Log(n)*** | -0.436 (p < .001) | -0.211 (p < .001) | -0.000 (p = 1.000) | -0.000 (p = 1.000) | 0.515 (p < .001) | -0.364 (p < .001) | -0.295 (p < .001) |
| ***Left-skewed*** | 0.862 (p < .001) | 0.550 (p < .001) | 0.000 (p = 1.000) | 0.000 (p = 1.000) | 0.078 (p < .001) | -0.130 (p < .001) | 1.701 (p < .001) |
| ***Right-skewed*** | 0.505 (p < .001) | 0.461 (p < .001) | 0.000 (p = 1.000) | 0.000 (p = 1.000) | 0.093 (p < .001) | 0.946 (p < .001) | -0.244 (p < .001) |
| ***Log(n):Left-skewed*** | -0.445 (p < .001) | -0.012 (p = .261) | 0.000 (p = 1.000) | 0.000 (p = 1.000) | 0.062 (p < .001) | 0.087 (p < .001) | -0.303 (p < .001) |
| ***Log(n):Right-skewed*** | -0.214 (p < .001) | -0.009 (p = .374) | 0.000 (p = 1.000) | 0.000 (p = 1.000) | -0.009 (p = .267) | -0.180 (p < .001) | 0.013 (p = .537) |

*Table S9. Linear mixed model results for evaluation metrics under age-skewed sampling conditions using normative models pre-trained on the UKB and adapted to the AIBL dataset. Models assess the influence of standardized and log-transformed sample size (n) and age sampling strategy (Representative, Left-skewed, Right-skewed) on model performance metrics (MSLL, SMSE, EV, Rho, ICC, the lower and upper tail HC%). All variables were standardized to allow comparison of effect sizes. Representative sampling serves as the reference level. Reported β coefficients and corresponding p-values indicate the direction and significance of each effect.*

| | ***MSLL* β (p-value)** | ***EV* β (p-value)** | ***SMSE* β (p-value)** | ***Rho* β (p-value)** | ***ICC* β (p-value)** | ***Lower tail HC %* β (p-value)** | ***Upper tail HC %* β (p-value)** |
|---|---|---|---|---|---|---|---|
| ***Intercept (1F1M)*** | 0.792 (p < .001) | 1.040 (p < .001) | -2.561 (p < .001) | -0.075 (p < .001) | 0.502 (p < .001) | -0.010 (p = .864) | -0.168 (p = .002) |
| ***Log(n)*** | -0.436 (p < .001) | -0.211 (p < .001) | -0.000 (p = 1.000) | -0.000 (p = 1.000) | 0.515 (p < .001) | -0.364 (p < .001) | -0.295 (p < .001) |
| ***10F1M*** | 0.056 (p < .001) | 0.060 (p < .001) | -0.000 (p = 1.000) | -0.000 (p = 1.000) | 0.002 (p = .781) | 0.097 (p < .001) | 0.021 (p = .018) |
| ***4F1M*** | 0.034 (p < .001) | 0.032 (p < .001) | 0.000 (p = 1.000) | -0.000 (p = 1.000) | 0.012 (p = .105) | 0.056 (p < .001) | 0.009 (p = .319) |
| ***1F4M*** | 0.025 (p = .011) | 0.017 (p < .001) | -0.000 (p = 1.000) | 0.000 (p = 1.000) | 0.031 (p < .001) | 0.056 (p < .001) | 0.033 (p < .001) |
| ***1F10M*** | 0.030 (p = .002) | 0.034 (p < .001) | -0.000 (p = 1.000) | -0.000 (p = 1.000) | 0.036 (p < .001) | 0.063 (p < .001) | 0.085 (p < .001) |
| ***Log(n):10F1M*** | -0.032 (p = .001) | 0.003 (p = .462) | 0.000 (p = 1.000) | -0.000 (p = 1.000) | -0.025 (p < .001) | -0.014 (p = .189) | -0.027 (p = .002) |
| ***Log(n):4F1M*** | -0.037 (p < .001) | 0.007 (p = .128) | -0.000 (p = 1.000) | -0.000 (p = 1.000) | -0.039 (p < .001) | -0.023 (p = .029) | -0.046 (p < .001) |
| ***Log(n):1F4M*** | -0.021 (p = .030) | 0.017 (p < .001) | 0.000 (p = 1.000) | 0.000 (p = 1.000) | -0.049 (p < .001) | -0.016 (p = .130) | -0.002 (p = .836) |

| | | | | | | | |
|---|---|---|---|---|---|---|---|
| ***Log(n):1F10M*** | 0.008 (p = .410) | 0.016 (p < .001) | 0.000 (p = 1.000) | 0.000 (p = 1.000) | -0.037 (p < .001) | 0.007 (p = .538) | -0.014 (p = .113) |

*Table S10. Linear mixed model results for evaluation metrics under sex-imbalanced sampling conditions using normative models pre-trained on the UKB and adapted to the AIBL dataset. Models assess the influence of standardized and log-transformed sample size (n) and sex ratio in the training set (1F:1M, 1F:4M, 1F:10M, 4F:1M, 10F:1M; F = female, M = male) on model performance metrics (MSLL, SMSE, EV, Rho, ICC, the lower and upper tail HC%). All variables were standardized to allow comparison of effect sizes. Representative sampling (1F:1M) serves as the reference level. Reported β coefficients and corresponding p-values indicate the direction and significance of each effect.*

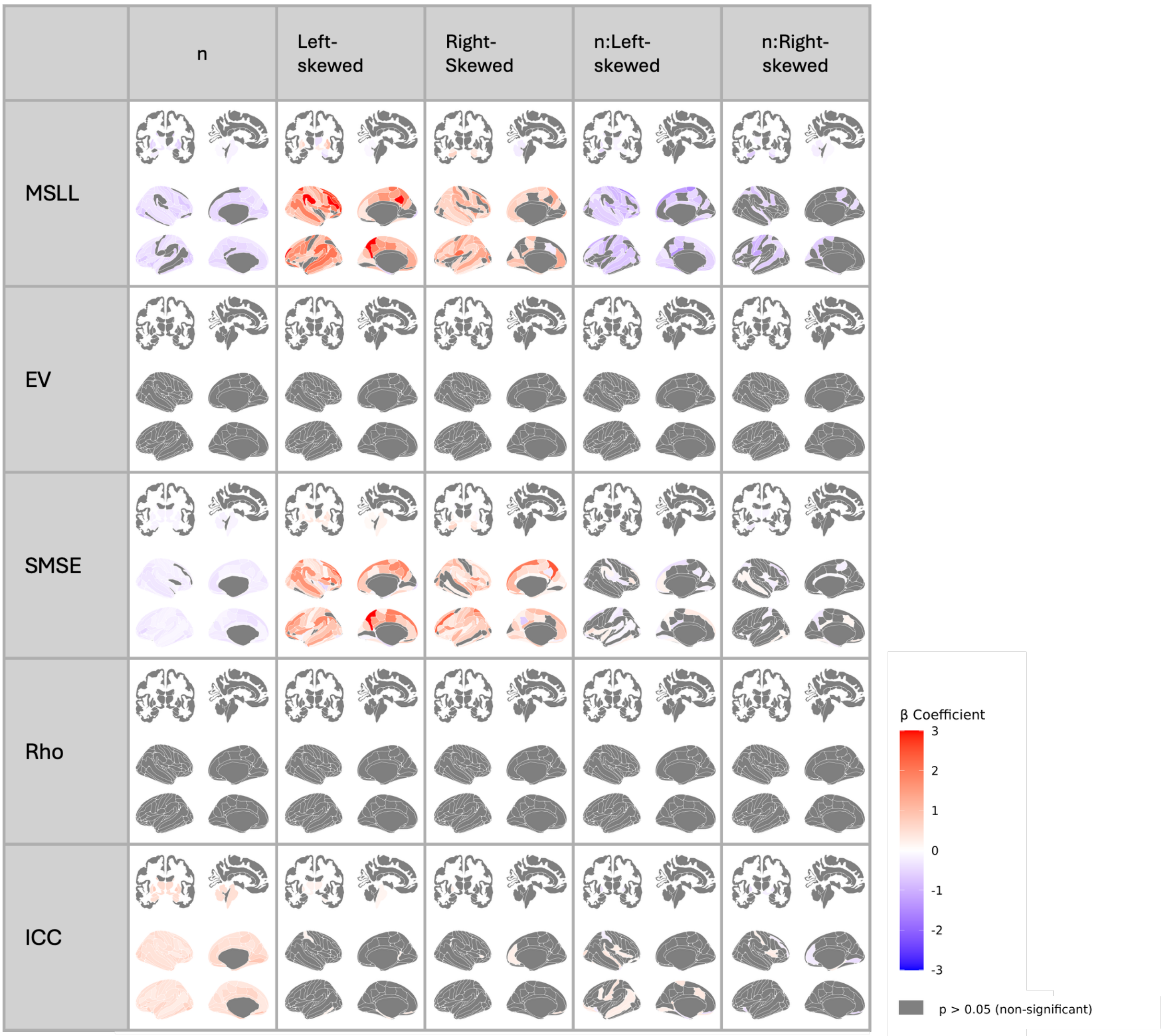

*Figure S30. Regional linear mixed-effects results for model-fit metrics in models pre-fitted in the UKB and adapted to AIBL with age-skewed samples. Standardized Betas (β) for each performance metric (MSLL, SMSE, EV, Rho, ICC; rows) are regressed on log-standardized sample size (n), age-distribution contrasts (left-skewed, right-skewed; representative = reference), and their interactions. Sex ratio is balanced (1:1). Grey shading denotes regions that did not survive FDR correction (p ≥ 0.05).*

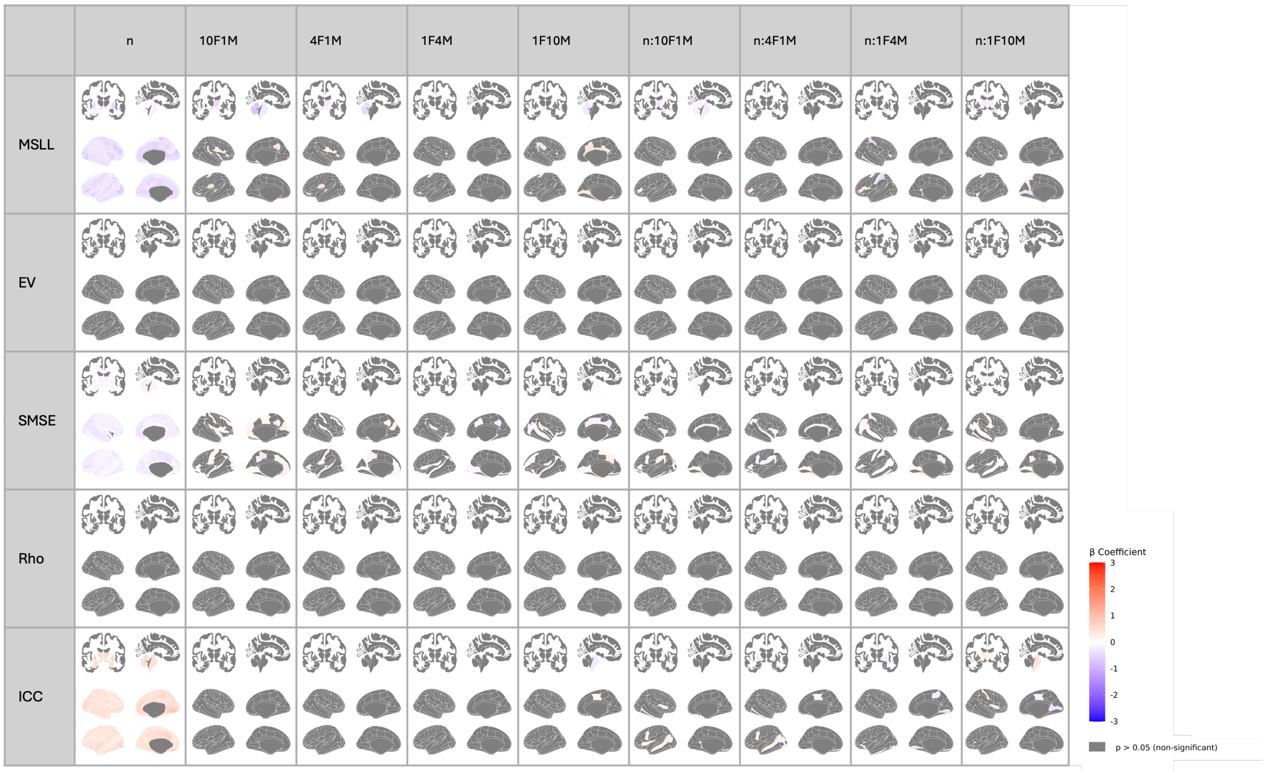

*Figure S31. Regional linear mixed-effects results for model-fit metrics in models pre-fitted in the UKB and adapted to AIBL with sex-imbalanced samples. Standardized Betas (β) for each performance metric (MSLL, SMSE, EV, Rho, ICC; rows) are regressed on log-standardized sample size (n), sex-ratio contrasts (female-to-male 10:1, 4:1, 1:1 [reference], 1:4, 1:10), and their interactions. Age distribution is representative. Grey shading denotes regions that did not survive FDR correction (p ≥ 0.05).*

#### 4.2. Z-scores errors

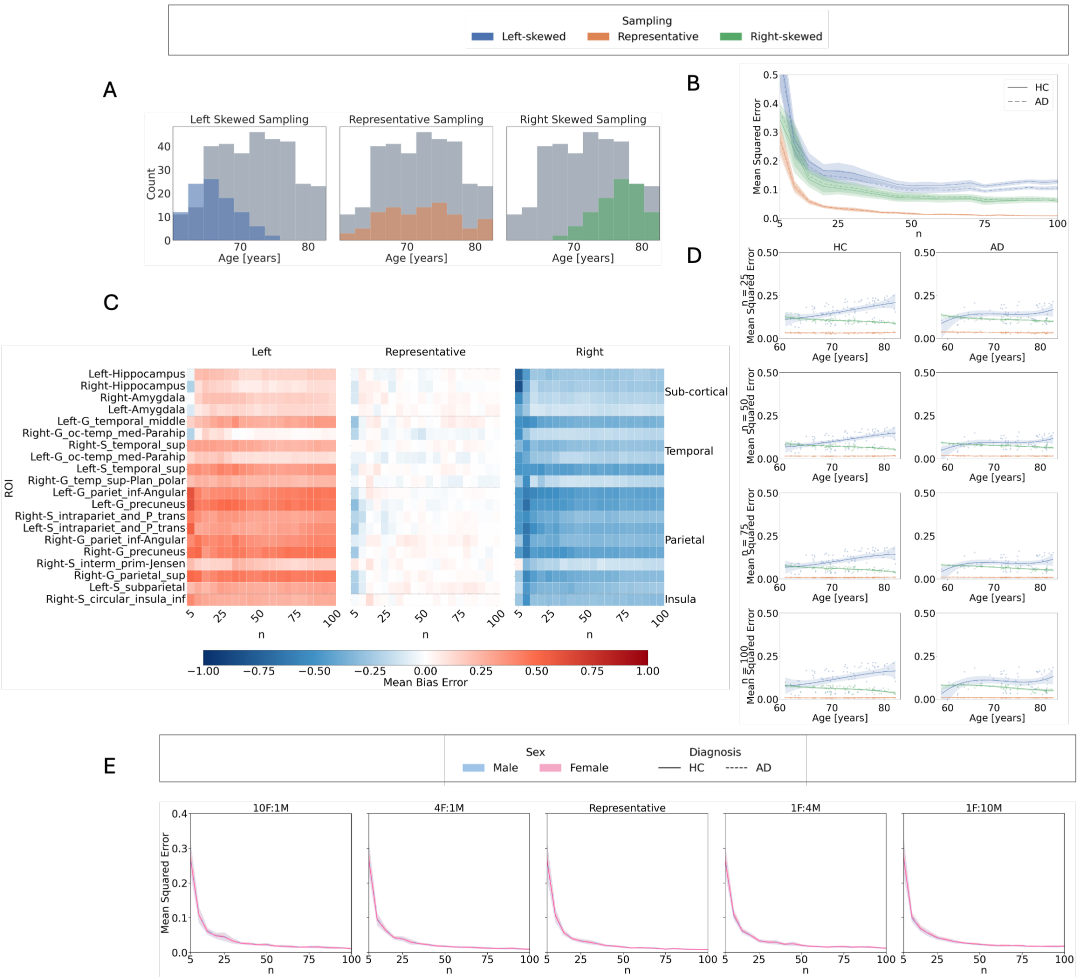

*Figure S32. Z-score errors in AIBL using models pretrained in UKB and adapted to AIBL: Influence of sample size, age distribution, and sex imbalance on normative model outcomes. A: Age distributions for left-skewed (younger-biased), representative, and right-skewed (older-biased) sampling strategies, compared to the full training set in the AIBL dataset. B: Mean squared error (MSE) of Z-scores relative to the full training set model across sample sizes, age distributions, and diagnostic groups. C: Mean bias error (MBE) per region across sample sizes and age distributions in the test set Shown are the 20 brain regions with the highness Cohen's d effect sizes based on models using the full training set. From left to right: results for left-skewed, representative, and right-skewed training sets. Blue indicates negative MBE (underestimation), red indicates positive MBE (overestimation), and white indicates close alignment with the full training set model. D: Cubic regression of MSE as a function of age, across sample sizes and sampling strategies. Left-skewed sampling shows increased errors in older individuals; right-skewed sampling shows increased errors in younger individuals. E: MSE across sample sizes and test set sex, obtained using sex-imbalanced training sets (female-to-male ratios: 10:1, 4:1, 1:1, 1:4, 1:10), all with representative age distributions.*

| | *MSE*<br>*β (p-value)* | *tOC*<br>*β (p-value)* |
|---|---|---|
| ***Intercept (HC, Representative)*** | -0.572 (p < .001) | -0.415 (p < .001) |
| ***Log(n)*** | -0.277 (p < .001) | -0.096 (p < .001) |
| ***AD*** | 0.009 (p = 0.613) | 0.792 (p < .001) |
| ***Left-skewed*** | 0.712 (p < .001) | -0.039 (p < .001) |
| ***Right-skewed*** | 0.356 (p < .001) | 0.252 (p < .001) |
| ***Log(n):AD*** | -0.011 (p = 0.305) | -0.036 (p < .001) |
| ***Log(n):Left-skewed*** | -0.193 (p < .001) | 0.026 (p < .001) |
| ***Log(n):Right-skewed*** | -0.070 (p < .001) | -0.048 (p < .001) |
| ***Left-skewed:AD*** | -0.111 (p < .001) | -0.019 (p = 0.113) |
| ***Right-skewed:AD*** | 0.036 (p = 0.020) | 0.213 (p < .001) |
| ***Log(n):Left-skewed:AD*** | 0.002 (p = 0.878) | 0.005 (p = 0.682) |
| ***Log(n):Right-skewed:AD*** | -0.014 (p = 0.368) | 0.005 (p = 0.650) |

*Table S11. Linear mixed model results for MSE and total outlier count (tOC) under age-skewed sampling for models adapted from the UK Biobank to AIBL. Models evaluate the influence of diagnosis (HC, AD), log-transformed and standardized sample size (n), and age sampling strategy (Representative, Left-skewed, Right-skewed) on standardized deviation score outcomes. Continuous variables were standardized to allow comparison of effect sizes. Representative sampling and HC are used as reference levels. Reported β coefficients and corresponding p-values indicate the direction and significance of the effects.*

| | *MSE*<br>*β (p-value)* | *tOC*<br>*β (p-value)* |
|---|---|---|
| ***Intercept (HC, 1F 1M)*** | -0.572 (p < .001) | -0.415 (p < .001) |
| ***Log(n)*** | 0.027 (p < .001) | 0.025 (p < .001) |
| ***AD*** | 0.010 (p = 0.132) | 0.014 (p = 0.002) |
| ***10F1M*** | 0.025 (p < .001) | 0.014 (p = 0.002) |
| ***4F1M*** | 0.046 (p < .001) | 0.016 (p < .001) |
| ***1F4M*** | 0.009 (p = 0.201) | 0.792 (p < .001) |
| ***1F10M*** | 0.003 (p = 0.746) | 0.049 (p < .001) |
| ***Log(n):AD*** | 0.002 (p = 0.838) | 0.031 (p < .001) |
| ***Log(n):10F1M*** | 0.002 (p = 0.844) | 0.009 (p = 0.188) |
| ***Log(n):4F1M*** | 0.002 (p = 0.817) | 0.015 (p = 0.031) |
| ***Log(n):1F4M*** | -0.277 (p < .001) | -0.096 (p < .001) |
| ***Log(n):1F10M*** | -0.001 (p = 0.923) | -0.003 (p = 0.554) |
| ***10F1M:AD*** | 0.002 (p = 0.750) | -0.005 (p = 0.281) |
| ***4F1M:AD*** | 0.006 (p = 0.334) | -0.003 (p = 0.560) |
| ***1F4M:AD*** | 0.002 (p = 0.699) | 0.003 (p = 0.477) |
| ***1F10M:AD*** | -0.011 (p = 0.111) | -0.036 (p < .001) |
| ***Log(n):10F1M:AD*** | -0.002 (p = 0.816) | -0.001 (p = 0.911) |
| ***Log(n):4F1M:AD*** | -0.003 (p = 0.778) | -0.008 (p = 0.278) |
| ***Log(n):1F4M:AD*** | -0.002 (p = 0.872) | -0.007 (p = 0.293) |
| ***Log(n):1F10M:AD*** | 0.000 (p = 0.996) | -0.006 (p = 0.370) |

*Table S12. Linear mixed model results for MSE and total outlier count (tOC) under sex-imbalanced sampling for models adapted from the UK Biobank to AIBL. Models evaluate the influence of diagnosis (HC, AD), log-transformed and standardized sample size (n), and sex ratio in the training set (1:1, 1F:4M, 1F:10M, 4F:1M, 10F:1M) on standardized deviation score outcomes. Continuous variables were standardized to allow comparison of effect sizes. The 1:1 ratio and HC are used as reference levels. Reported β coefficients and corresponding p-values indicate the direction and significance of the effects.*

#### 4.3. Clinical validation

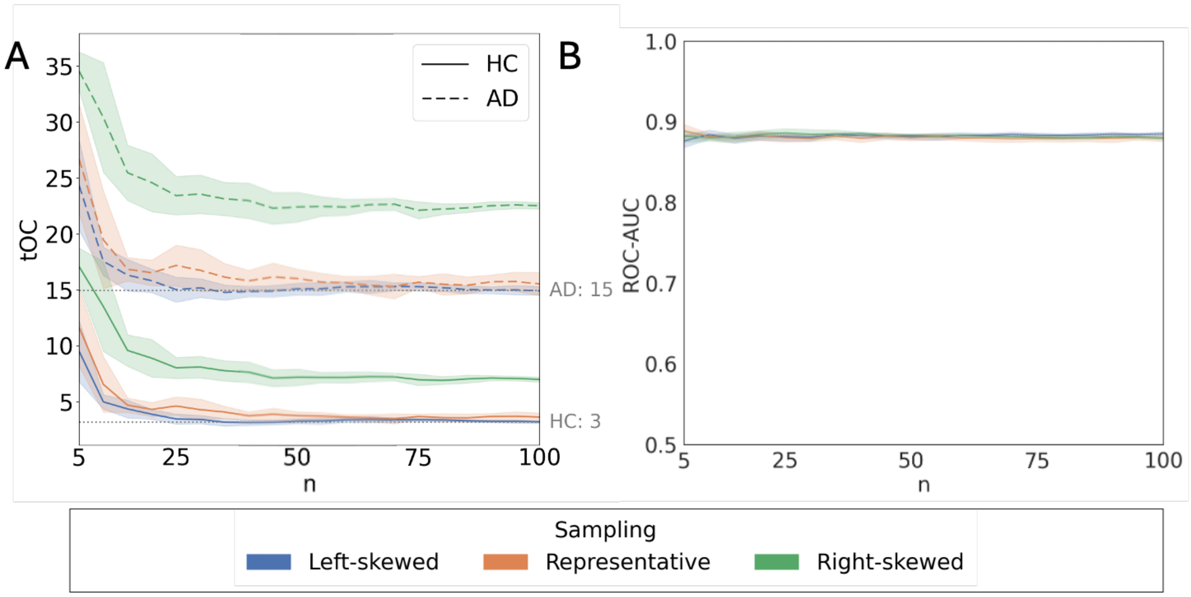

*Figure S33. Clinical validation for models pre-fitted in UKB and adapted to AIBL: effect of sample size and age distributions in outlier detection and classification performance. A: Average total Outlier Count (tOC) in the AD and HC are represented with dashed and solid lines respectively as a function of sample size (n). The shaded areas indicate the standard deviation across iterations. Dotted grey lines correspond to the estimations of tOC for HC and AD groups obtained with full sample size. B: ROC-AUC of Support Vector Classifier with 10-fold cross-validation as a function of sample size in the training set. The solid lines represent the average AUC across iterations for each sampling strategy. The dotted line shows the performance of the models trained with full sample size*

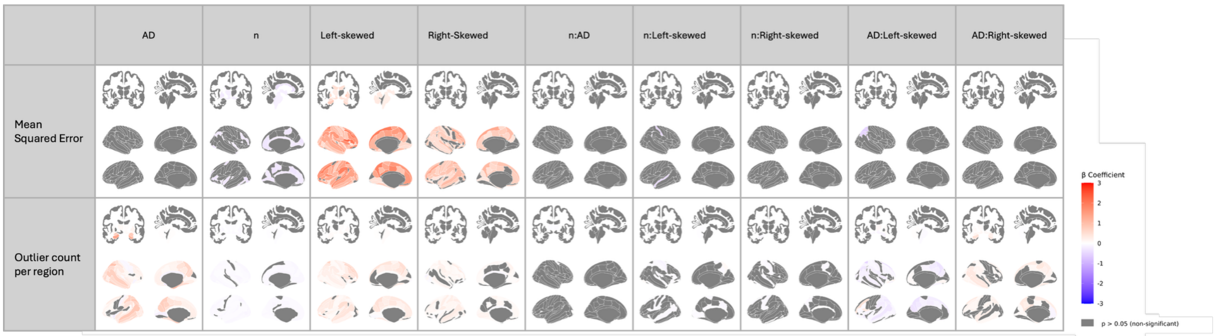

*Figure S34. Regional linear mixed-effects results for deviation-score in models pre-fitted in the UKB and adapted to AIBL with age-skewed samples. Standardized Betas (β) are shown for the mean-squared error of z-scores relative to the full reference model and for the count of extreme z-score outliers per region (rows). Predictors include Alzheimer's disease diagnosis (AD), log-standardized sample size (n), age-distribution contrasts (left-skewed, right-skewed; representative = reference), and their two-way interactions with n and AD. Sex ratio is balanced (1:1). Grey shading denotes regions that did not survive FDR correction (p ≥ 0.05).*

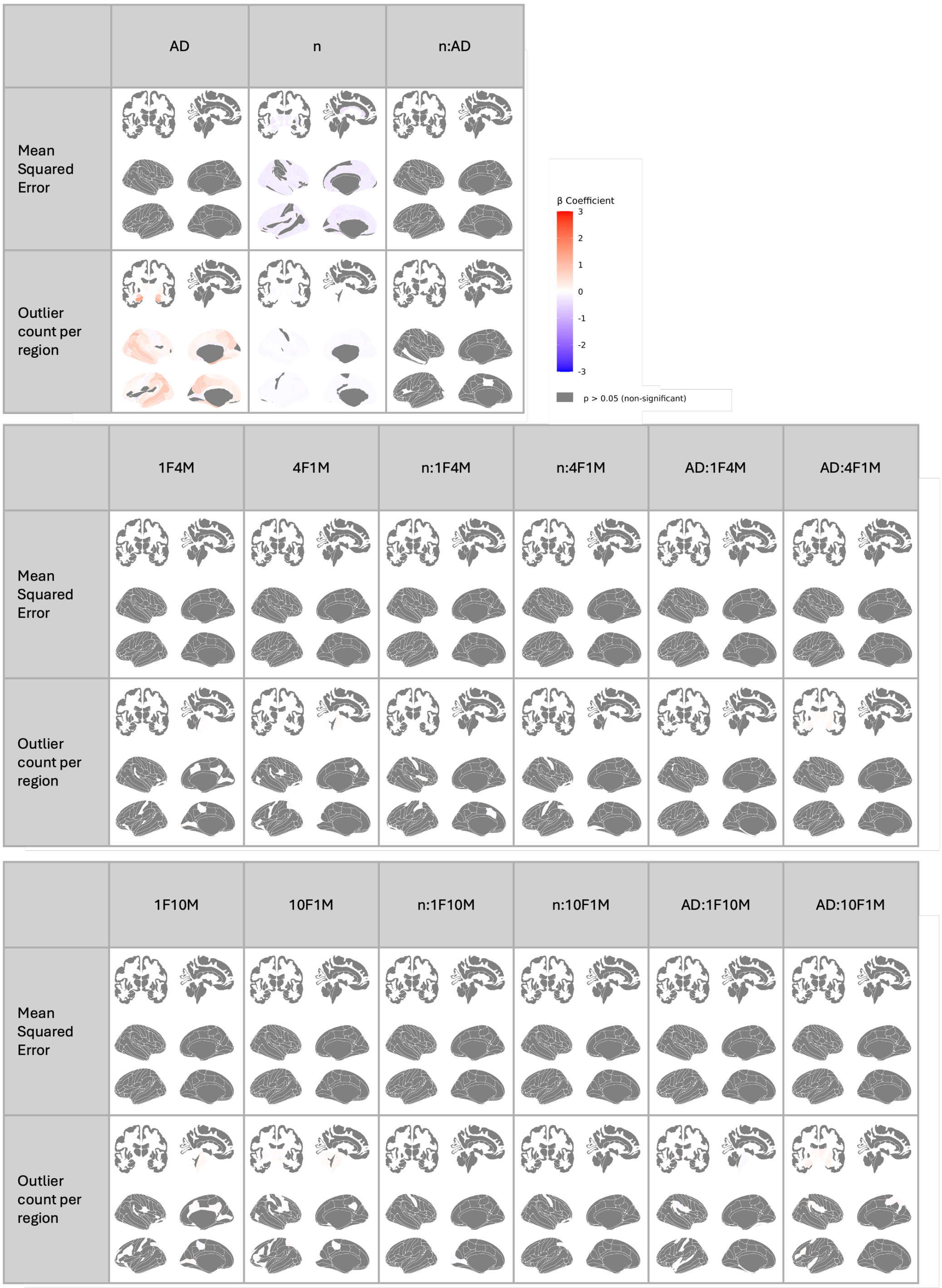

*Figure S35. Regional linear mixed-effects results for deviation-score metrics in models pre-fitted in the UKB and adapted to AIBL with sex-imbalanced samples. Standardized Betas (β) are shown for the mean-squared error of z-scores relative to the full reference model and for the count of extreme z-score outliers per region (rows). Predictors include Alzheimer's disease diagnosis (AD), log-standardized sample size (n), sex-ratio contrasts (female-to-male 10:1, 4:1, 1:1 [reference], 1:4, 1:10), and their two-way interactions with n and AD. Age distribution is representative. Grey shading denotes regions that did not survive FDR correction (p ≥ 0.05).*

#### 5. Control analyses

##### 5.1. Age and sex distributions and expected sampling effects

Age and sex distributions were first summarized for each dataset and split (train HC, test HC, test AD) separately for males and females and are reported as mean ± SD and interquartile range (P25–P75).

To characterize the expected age distributions induced by the sampling procedures, we simulated the sampling used in the main analyses by repeatedly drawing samples from the training set. Because the sampling strategy enforces the same quantile-based age stratification independently of sample size, the expected age distribution is theoretically invariant with respect to n, with smaller n only introducing additional sampling noise. We therefore performed the simulation at a single representative sample size (n = 200). For each ratio, 1000 independent samples were drawn. Ages were pooled across repetitions within each ratio × sex combination and summarized as mean ± SD and P25–P75.

| | *Split* | *Sex* | *Mean ± sd* | *p25_p75* |
|---|---|---|---|---|
| *OASIS-3* | Train HC | Male | 68.99 ± 7.55 | 65.6–74.1 |
| | | Female | 67.66 ± 7.97 | 63.2–73.7 |
| | Test HC | Male | 69.05 ± 7.67 | 66.1–73.6 |
| | | Female | 67.69 ± 7.97 | 63.9–73.3 |
| | Test AD | Male | 73.69 ± 6.06 | 69.6–78.6 |
| | | Female | 73.91 ± 5.93 | 70.4–77.9 |
| *AIBL* | Train HC | Male | 72.43 ± 5.19 | 67.9–76.0 |
| | | Female | 72.02 ± 5.56 | 67.7–76.2 |
| | Test HC | Male | 72.38 ± 5.46 | 69.0–77.0 |
| | | Female | 72.47 ± 5.42 | 68.8–77.3 |
| | Test AD | Male | 73.51 ± 6.59 | 69.3–79.0 |
| | | Female | 73.65 ± 6.75 | 71.5–79.8 |
| *UKB* | Train HC | Male | 65.22 ± 7.80 | 59.3–71.3 |
| | | Female | 63.87 ± 7.52 | 58.0–69.7 |
| | Test HC | Male | 65.20 ± 7.72 | 59.3–71.2 |
| | | Female | 63.85 ± 7.51 | 58.0–69.6 |

*Table S13. Empirical age distributions by sex for the training and test splits of OASIS-3, AIBL, and UK Biobank. Age is reported as mean ± SD and interquartile range (P25–P75) separately for males and females in each split (Train HC, Test HC, Test AD where available).*

| | *Sampling* | *Sex* | *Mean ± sd* | *p25_p75* |
|---|---|---|---|---|
| *OASIS-3* | Representative | Male | 68.24 ± 7.75 | 64.4–73.8 |
| | | Female | 68.20 ± 7.82 | 64.1–73.8 |
| | Left | Male | 57.58 ± 5.23 | 54.0–60.0 |
| | | Female | 57.05 ± 5.40 | 53.3–60.1 |
| | Right | Male | 72.10 ± 6.08 | 67.9–76.8 |
| | | Female | 71.94 ± 5.89 | 68.2–75.8 |
| *AIBL* | Representative | Male | 72.19 ± 5.34 | 67.7–76.0 |
| | | Female | 72.18 ± 5.39 | 68.0–76.2 |
| | Left | Male | 67.54 ± 3.16 | 65.5–69.1 |
| | | Female | 67.29 ± 3.19 | 65.4–69.0 |
| | Right | Male | 76.04 ± 3.49 | 74.0–78.4 |
| | | Female | 75.66 ± 3.21 | 73.9–77.4 |

*Table S14. Expected age distributions under representative, left-skewed, and right-skewed age sampling for OASIS-3 and AIBL. Expected distributions were estimated by repeated simulation of the age-skewed sampling procedure at n = 200 across 1000 independent samples and are reported as mean ± SD and P25–P75 separately for males and females.*

| | *Sampling* | *Sex* | *Mean ± sd* | *p25_p75* |
|---|---|---|---|---|
| *OASIS-3* | 1F1M | Male | 68.24 ± 7.74 | 64.4–73.8 |
| | | Female | 68.19 ± 7.83 | 64.1–73.8 |
| | 1F4M | Male | 68.24 ± 7.74 | 64.4–73.8 |
| | | Female | 68.19 ± 7.83 | 64.1–73.8 |
| | 1F10M | Male | 68.23 ± 7.74 | 64.4–73.8 |
| | | Female | 68.2 ± 7.85 | 64.1–73.8 |

| | | | | |
|---|---|---|---|---|
| | 4F1M | Male | 68.24 ± 7.73 | 64.4–73.8 |
| | | Female | 68.19 ± 7.83 | 64.1–73.8 |
| | 10F1M | Male | 68.22 ± 7.78 | 64.2–73.9 |
| | | Female | 68.19 ± 7.84 | 64.1–73.8 |
| ***AIBL*** | 1F1M | Male | 72.19 ± 5.34 | 67.6–76.0 |
| | | Female | 72.19 ± 5.39 | 68.1–76.2 |
| | 1F4M | Male | 72.19 ± 5.34 | 67.6–76.0 |
| | | Female | 72.18 ± 5.39 | 68.0–76.2 |
| | 1F10M | Male | 72.19 ± 5.34 | 67.6–76.0 |
| | | Female | 72.19 ± 5.39 | 68.0–76.2 |
| | 4F1M | Male | 72.2 ± 5.33 | 67.5–76.0 |
| | | Female | 72.18 ± 5.39 | 68.1–76.2 |
| | 10F1M | Male | 72.18 ± 5.33 | 67.6–76.0 |
| | | Female | 72.18 ± 5.39 | 68.0–76.2 |

*Table S15. Expected age distributions under sex-imbalance sampling for OASIS-3 and AIBL. Sex ratios include 1F:1M (representative), 1F:4M, 1F:10M, 4F:1M, and 10F:1M. Expected distributions were estimated by repeated simulation of the sampling procedure at n = 200 across 1000 independent samples and are reported as mean ± SD and P25–P75 separately for males and females.*

#### 5.2. Statistical models including age and sex

| | ***OASIS-3*** | | ***AIBL*** | |
|---|---|---|---|---|
| | **MSE** β **(p-value)** | **tOC** β **(p-value)** | **MSE** β **(p-value)** | **tOC** β **(p-value)** |
| ***Intercept*** | 0.113 (p = 0.824) | 0.202 (p = 0.916) | 0.245 (p = 0.876) | 0.614 (p = 0.874) |
| ***n*** | -1.237 (p < .001) | 0.539 (p < .001) | -0.972 (p = 0.096) | 1.112 (p = 0.003) |
| ***AD*** | -0.964 (p = 0.242) | -0.795 (p = 0.798) | -0.132 (p = 0.952) | -7.493 (p = 0.165) |
| ***left*** | -6.233 (p < .001) | -3.919 (p < .001) | -12.217 (p < .001) | -5.242 (p < .001) |
| ***right*** | 1.270 (p < .001) | -0.604 (p < .001) | 8.295 (p < .001) | -1.852 (p < .001) |
| ***n:AD*** | 1.197 (p < .001) | 0.480 (p = 0.017) | -0.285 (p = 0.726) | -0.914 (p = 0.079) |
| ***n:left*** | 4.473 (p < .001) | 2.006 (p < .001) | 6.366 (p < .001) | 1.041 (p = 0.049) |
| ***n:right*** | -1.482 (p < .001) | 0.077 (p = 0.662) | -6.562 (p < .001) | -0.793 (p = 0.134) |
| ***left:AD*** | -2.744 (p < .001) | -4.194 (p < .001) | 0.434 (p = 0.705) | -4.832 (p < .001) |
| ***right:AD*** | -0.142 (p = 0.730) | 0.550 (p = 0.054) | -0.003 (p = 0.998) | 1.009 (p = 0.171) |
| ***n:left:AD*** | 0.558 (p = 0.175) | 0.389 (p = 0.172) | -1.107 (p = 0.335) | 0.962 (p = 0.192) |
| ***n:right:AD*** | -0.415 (p = 0.313) | 0.253 (p = 0.375) | -1.304 (p = 0.256) | 0.368 (p = 0.618) |
| ***age*** | -0.005 (p = 0.477) | -0.008 (p = 0.786) | -0.008 (p = 0.706) | -0.014 (p = 0.795) |
| ***left:age*** | 0.104 (p < .001) | 0.065 (p < .001) | 0.183 (p < .001) | 0.076 (p < .001) |
| ***right:age*** | -0.017 (p < .001) | 0.009 (p < .001) | -0.108 (p < .001) | 0.024 (p < .001) |
| ***AD:age*** | 0.014 (p = 0.228) | 0.017 (p = 0.685) | 0.003 (p = 0.922) | 0.108 (p = 0.141) |
| ***left:AD:age*** | 0.037 (p < .001) | 0.057 (p < .001) | -0.007 (p = 0.650) | 0.072 (p < .001) |
| ***right:AD:age*** | 0.002 (p = 0.686) | -0.007 (p = 0.064) | 0.002 (p = 0.884) | -0.014 (p = 0.167) |
| ***n:age*** | 0.012 (p < .001) | -0.007 (p < .001) | 0.008 (p = 0.330) | -0.014 (p = 0.006) |
| ***n:left:age*** | -0.075 (p < .001) | -0.034 (p < .001) | -0.094 (p < .001) | -0.016 (p = 0.030) |
| ***n:right:age*** | 0.019 (p < .001) | -0.001 (p = 0.690) | 0.084 (p < .001) | 0.010 (p = 0.152) |
| ***n:AD:age*** | -0.017 (p < .001) | -0.005 (p = 0.073) | 0.003 (p = 0.820) | 0.013 (p = 0.068) |
| ***n:left:AD:age*** | -0.006 (p = 0.299) | -0.005 (p = 0.166) | 0.016 (p = 0.294) | -0.015 (p = 0.132) |
| ***n:right:AD:age*** | 0.005 (p = 0.362) | -0.003 (p = 0.383) | 0.016 (p = 0.318) | -0.005 (p = 0.612) |
| ***sex*** | 0.073 (p = 0.809) | -0.521 (p = 0.647) | -0.290 (p = 0.759) | -0.660 (p = 0.778) |
| ***left:sex*** | 0.002 (p = 0.990) | 0.722 (p < .001) | 0.057 (p = 0.909) | 1.052 (p = 0.001) |
| ***right:sex*** | 0.018 (p = 0.907) | 0.228 (p = 0.029) | 0.288 (p = 0.563) | 0.854 (p = 0.008) |
| ***AD:sex*** | 0.009 (p = 0.987) | 1.404 (p = 0.475) | 0.323 (p = 0.806) | 5.076 (p = 0.120) |
| ***left:AD:sex*** | 0.086 (p = 0.741) | 0.347 (p = 0.055) | 1.359 (p = 0.050) | 0.541 (p = 0.224) |
| ***right:AD:sex*** | -0.106 (p = 0.685) | -0.438 (p = 0.015) | -0.694 (p = 0.317) | -1.900 (p < .001) |
| ***n:sex*** | -0.034 (p = 0.751) | -0.200 (p = 0.007) | 0.187 (p = 0.595) | -0.302 (p = 0.182) |
| ***n:left:sex*** | -0.222 (p = 0.142) | -0.285 (p = 0.006) | -0.394 (p = 0.430) | 0.061 (p = 0.850) |
| ***n:right:sex*** | -0.089 (p = 0.554) | -0.024 (p = 0.817) | -0.555 (p = 0.266) | 0.240 (p = 0.453) |
| ***n:AD:sex*** | -0.067 (p = 0.716) | 0.259 (p = 0.042) | -0.178 (p = 0.717) | 1.452 (p < .001) |
| ***n:left:AD:sex*** | -0.176 (p = 0.499) | -0.067 (p = 0.712) | -0.170 (p = 0.806) | -0.605 (p = 0.174) |
| ***n:right:AD:sex*** | 0.433 (p = 0.096) | 0.070 (p = 0.699) | 1.254 (p = 0.071) | -0.708 (p = 0.112) |
| ***age:sex*** | -0.001 (p = 0.803) | 0.006 (p = 0.696) | 0.004 (p = 0.743) | 0.008 (p = 0.793) |
| ***left:age:sex*** | 0.002 (p = 0.466) | -0.011 (p < .001) | -0.002 (p = 0.767) | -0.015 (p < .001) |
| ***right:age:sex*** | -0.000 (p = 0.884) | -0.003 (p = 0.028) | -0.003 (p = 0.635) | -0.012 (p = 0.009) |
| ***AD:age:sex*** | -0.000 (p = 0.993) | -0.018 (p = 0.517) | -0.004 (p = 0.803) | -0.065 (p = 0.143) |
| ***left:AD:age:sex*** | -0.001 (p = 0.826) | -0.001 (p = 0.757) | -0.018 (p = 0.062) | -0.006 (p = 0.310) |
| ***right:AD:age:sex*** | 0.001 (p = 0.709) | 0.006 (p = 0.020) | 0.009 (p = 0.364) | 0.024 (p < .001) |
| ***n:age:sex*** | 0.000 (p = 0.787) | 0.002 (p = 0.026) | -0.003 (p = 0.538) | 0.004 (p = 0.245) |
| ***n:left:age:sex*** | 0.002 (p = 0.330) | 0.004 (p = 0.004) | 0.006 (p = 0.373) | -0.001 (p = 0.789) |
| ***n:right:age:sex*** | 0.001 (p = 0.540) | 0.000 (p = 0.764) | 0.007 (p = 0.300) | -0.003 (p = 0.510) |
| ***n:AD:age:sex*** | 0.001 (p = 0.717) | -0.004 (p = 0.047) | 0.003 (p = 0.697) | -0.019 (p < .001) |
| ***n:left:AD:age:sex*** | 0.003 (p = 0.460) | -0.000 (p = 0.931) | 0.002 (p = 0.834) | 0.008 (p = 0.183) |

| n:right:AD:age:sex | -0.006 (p = 0.111) | -0.001 (p = 0.707) | -0.016 (p = 0.088) | 0.010 (p = 0.111) |
|---|---|---|---|---|

*Table S16. Linear mixed model results for MSE and total outlier count (tOC) under age-skewed sampling for models trained with OASIS-3 or AIBL. Models evaluate the influence of diagnosis (HC, AD), log-transformed and standardized sample size (n), age sampling strategy (Representative, Left-skewed, Right-skewed), age, sex, and their interactions on standardized deviation score outcomes. Continuous variables were standardized to allow comparison of effect sizes. Representative sampling, HC, and female sex are used as reference levels. Reported β coefficients and corresponding p-values indicate the direction and significance of the effects.*

| | ***OASIS-3*** | | ***AIBL*** | |
|---|---|---|---|---|
| | **MSE β (p-value)** | **tOC β (p-value)** | **MSE β (p-value)** | **tOC β (p-value)** |
| ***Intercept*** | 0.113 (p = 0.674) | 0.202 (p = 0.905) | 0.245 (p = 0.754) | 0.614 (p = 0.868) |
| ***n*** | -1.237 (p < .001) | 0.539 (p < .001) | -0.972 (p < .001) | 1.112 (p < .001) |
| ***AD*** | -0.964 (p = 0.027) | -0.795 (p = 0.771) | -0.132 (p = 0.903) | -7.493 (p = 0.144) |
| ***10F1M*** | 0.461 (p = 0.002) | 0.052 (p = 0.618) | 0.649 (p = 0.086) | -0.027 (p = 0.934) |
| ***4F1M*** | 0.129 (p = 0.383) | 0.055 (p = 0.595) | 0.381 (p = 0.313) | -0.040 (p = 0.901) |
| ***1F4M*** | 0.080 (p = 0.587) | -0.155 (p = 0.135) | -0.033 (p = 0.931) | 0.295 (p = 0.361) |
| ***1F10M*** | -0.061 (p = 0.677) | -0.101 (p = 0.331) | -0.097 (p = 0.797) | 0.474 (p = 0.142) |
| ***n:AD*** | 1.197 (p < .001) | 0.480 (p < .001) | -0.285 (p = 0.443) | -0.914 (p = 0.004) |
| ***n:10F1M*** | -0.150 (p = 0.308) | 0.035 (p = 0.732) | -0.587 (p = 0.120) | 0.067 (p = 0.836) |
| ***n:4F1M*** | 0.032 (p = 0.826) | 0.066 (p = 0.523) | -0.452 (p = 0.231) | 0.183 (p = 0.571) |
| ***n:1F4M*** | -0.178 (p = 0.228) | 0.020 (p = 0.844) | 0.285 (p = 0.450) | 0.062 (p = 0.849) |
| ***n:1F10M*** | 0.338 (p = 0.022) | -0.157 (p = 0.129) | 0.469 (p = 0.214) | -0.165 (p = 0.610) |
| ***10F1M:AD*** | 0.729 (p = 0.002) | -0.558 (p < .001) | -0.102 (p = 0.845) | -0.071 (p = 0.874) |
| ***4F1M:AD*** | 0.351 (p = 0.142) | -0.404 (p = 0.016) | 0.050 (p = 0.925) | -0.140 (p = 0.755) |
| ***1F4M:AD*** | -0.535 (p = 0.025) | 0.190 (p = 0.257) | 0.102 (p = 0.846) | -0.201 (p = 0.654) |
| ***1F10M:AD*** | -0.890 (p < .001) | 0.407 (p = 0.015) | -0.028 (p = 0.958) | -0.067 (p = 0.881) |
| ***n:10F1M:AD*** | -0.462 (p = 0.053) | -0.078 (p = 0.641) | -0.020 (p = 0.969) | 0.000 (p = 0.999) |
| ***n:4F1M:AD*** | -0.239 (p = 0.318) | -0.022 (p = 0.897) | -0.053 (p = 0.919) | 0.105 (p = 0.814) |
| ***n:1F4M:AD*** | 0.291 (p = 0.223) | -0.056 (p = 0.739) | -0.548 (p = 0.297) | 0.156 (p = 0.728) |
| ***n:1F10M:AD*** | 0.302 (p = 0.206) | -0.198 (p = 0.239) | -0.492 (p = 0.349) | -0.264 (p = 0.556) |
| ***age*** | -0.005 (p = 0.180) | -0.008 (p = 0.757) | -0.008 (p = 0.450) | -0.014 (p = 0.785) |
| ***10F1M:age*** | -0.000 (p = 0.819) | -0.001 (p = 0.667) | -0.004 (p = 0.463) | 0.000 (p = 0.931) |
| ***4F1M:age*** | 0.001 (p = 0.692) | -0.000 (p = 0.789) | -0.002 (p = 0.659) | 0.001 (p = 0.855) |
| ***1F4M:age*** | -0.002 (p = 0.245) | 0.002 (p = 0.158) | -0.001 (p = 0.889) | -0.004 (p = 0.336) |
| ***1F10M:age*** | -0.002 (p = 0.426) | 0.001 (p = 0.441) | -0.001 (p = 0.802) | -0.007 (p = 0.121) |
| ***AD:age*** | 0.014 (p = 0.023) | 0.017 (p = 0.645) | 0.003 (p = 0.844) | 0.108 (p = 0.122) |
| ***10F1M:AD:age*** | -0.010 (p = 0.002) | 0.006 (p = 0.011) | 0.001 (p = 0.850) | -0.001 (p = 0.826) |
| ***4F1M:AD:age*** | -0.005 (p = 0.137) | 0.005 (p = 0.035) | -0.001 (p = 0.928) | 0.001 (p = 0.906) |
| ***1F4M:AD:age*** | 0.008 (p = 0.023) | -0.003 (p = 0.283) | -0.001 (p = 0.906) | 0.003 (p = 0.616) |
| ***1F10M:AD:age*** | 0.013 (p < .001) | -0.005 (p = 0.032) | 0.001 (p = 0.879) | 0.002 (p = 0.802) |
| ***n:age*** | 0.012 (p < .001) | -0.007 (p < .001) | 0.008 (p = 0.033) | -0.014 (p < .001) |
| ***n:10F1M:age*** | -0.000 (p = 0.817) | -0.001 (p = 0.375) | 0.006 (p = 0.230) | -0.002 (p = 0.718) |
| ***n:4F1M:age*** | -0.002 (p = 0.328) | -0.002 (p = 0.250) | 0.004 (p = 0.432) | -0.003 (p = 0.469) |
| ***n:1F4M:age*** | 0.003 (p = 0.121) | -0.001 (p = 0.644) | -0.004 (p = 0.481) | -0.001 (p = 0.837) |
| ***n:1F10M:age*** | -0.003 (p = 0.130) | 0.002 (p = 0.162) | -0.006 (p = 0.259) | 0.002 (p = 0.679) |
| ***n:AD:age*** | -0.017 (p < .001) | -0.005 (p = 0.002) | 0.003 (p = 0.619) | 0.013 (p = 0.003) |
| ***n:10F1M:AD:age*** | 0.006 (p = 0.052) | 0.002 (p = 0.503) | 0.000 (p = 0.985) | -0.000 (p = 0.943) |
| ***n:4F1M:AD:age*** | 0.003 (p = 0.314) | 0.001 (p = 0.795) | 0.001 (p = 0.935) | -0.002 (p = 0.748) |
| ***n:1F4M:AD:age*** | -0.004 (p = 0.226) | 0.001 (p = 0.772) | 0.007 (p = 0.340) | -0.002 (p = 0.705) |
| ***n:1F10M:AD:age*** | -0.004 (p = 0.222) | 0.002 (p = 0.350) | 0.006 (p = 0.371) | 0.003 (p = 0.609) |
| ***sex*** | 0.073 (p = 0.648) | -0.521 (p = 0.603) | -0.290 (p = 0.539) | -0.660 (p = 0.767) |
| ***10F1M:sex*** | -0.134 (p = 0.125) | -0.003 (p = 0.960) | -0.308 (p = 0.177) | 0.083 (p = 0.672) |
| ***4F1M:sex*** | -0.019 (p = 0.826) | -0.012 (p = 0.849) | -0.135 (p = 0.554) | 0.071 (p = 0.716) |
| ***1F4M:sex*** | 0.007 (p = 0.939) | 0.072 (p = 0.244) | 0.071 (p = 0.755) | -0.101 (p = 0.605) |
| ***1F10M:sex*** | 0.042 (p = 0.629) | 0.046 (p = 0.455) | 0.219 (p = 0.337) | -0.158 (p = 0.419) |
| ***AD:sex*** | 0.009 (p = 0.975) | 1.404 (p = 0.417) | 0.323 (p = 0.623) | 5.076 (p = 0.102) |
| ***10F1M:AD:sex*** | -0.519 (p < .001) | 0.369 (p < .001) | 0.092 (p = 0.773) | 0.438 (p = 0.107) |
| ***4F1M:AD:sex*** | -0.267 (p = 0.078) | 0.241 (p = 0.024) | -0.026 (p = 0.935) | 0.340 (p = 0.211) |
| ***1F4M:AD:sex*** | 0.300 (p = 0.047) | -0.194 (p = 0.068) | -0.036 (p = 0.909) | -0.024 (p = 0.930) |
| ***1F10M:AD:sex*** | 0.552 (p < .001) | -0.333 (p = 0.002) | -0.002 (p = 0.995) | -0.136 (p = 0.616) |
| ***n:sex*** | -0.034 (p = 0.584) | -0.200 (p < .001) | 0.187 (p = 0.246) | -0.302 (p = 0.029) |
| ***n:10F1M:sex*** | 0.009 (p = 0.920) | -0.068 (p = 0.273) | 0.082 (p = 0.721) | -0.082 (p = 0.673) |
| ***n:4F1M:sex*** | -0.063 (p = 0.469) | -0.074 (p = 0.230) | -0.003 (p = 0.991) | -0.118 (p = 0.546) |
| ***n:1F4M:sex*** | 0.061 (p = 0.487) | -0.016 (p = 0.795) | -0.217 (p = 0.342) | -0.010 (p = 0.959) |
| ***n:1F10M:sex*** | -0.018 (p = 0.834) | 0.090 (p = 0.145) | -0.286 (p = 0.210) | 0.147 (p = 0.451) |
| ***n:AD:sex*** | -0.067 (p = 0.531) | 0.259 (p < .001) | -0.178 (p = 0.428) | 1.452 (p < .001) |
| ***n:10F1M:AD:sex*** | 0.305 (p = 0.044) | 0.044 (p = 0.680) | 0.011 (p = 0.972) | 0.162 (p = 0.551) |
| ***n:4F1M:AD:sex*** | 0.199 (p = 0.188) | 0.052 (p = 0.628) | 0.042 (p = 0.895) | 0.083 (p = 0.760) |
| ***n:1F4M:AD:sex*** | -0.188 (p = 0.214) | 0.061 (p = 0.566) | 0.305 (p = 0.337) | -0.073 (p = 0.787) |
| ***n:1F10M:AD:sex*** | -0.302 (p = 0.046) | 0.128 (p = 0.229) | 0.337 (p = 0.288) | 0.088 (p = 0.747) |
| ***age:sex*** | -0.001 (p = 0.637) | 0.006 (p = 0.658) | 0.004 (p = 0.510) | 0.008 (p = 0.782) |
| ***10F1M:age:sex*** | -0.001 (p = 0.360) | 0.000 (p = 0.998) | 0.002 (p = 0.570) | -0.001 (p = 0.693) |
| ***4F1M:age:sex*** | -0.001 (p = 0.363) | -0.000 (p = 0.964) | 0.000 (p = 0.887) | -0.001 (p = 0.700) |

| | | | | |
|---|---|---|---|---|
| ***1F4M:age:sex*** | 0.001 (p = 0.365) | -0.001 (p = 0.317) | 0.001 (p = 0.856) | 0.002 (p = 0.526) |
| ***1F10M:age:sex*** | 0.002 (p = 0.159) | -0.000 (p = 0.676) | 0.000 (p = 0.939) | 0.003 (p = 0.308) |
| ***AD:age:sex*** | -0.000 (p = 0.987) | -0.018 (p = 0.461) | -0.004 (p = 0.617) | -0.065 (p = 0.124) |
| ***10F1M:AD:age:sex*** | 0.007 (p < .001) | -0.004 (p = 0.008) | -0.001 (p = 0.794) | -0.004 (p = 0.246) |
| ***4F1M:AD:age:sex*** | 0.004 (p = 0.071) | -0.003 (p = 0.059) | 0.000 (p = 0.926) | -0.004 (p = 0.333) |
| ***1F4M:AD:age:sex*** | -0.004 (p = 0.044) | 0.003 (p = 0.088) | 0.000 (p = 0.968) | 0.000 (p = 0.952) |
| ***1F10M:AD:age:sex*** | -0.008 (p < .001) | 0.004 (p = 0.007) | -0.000 (p = 0.933) | 0.002 (p = 0.684) |
| ***n:age:sex*** | 0.000 (p = 0.642) | 0.002 (p < .001) | -0.003 (p = 0.178) | 0.004 (p = 0.057) |
| ***n:10F1M:age:sex*** | 0.001 (p = 0.275) | 0.001 (p = 0.112) | -0.000 (p = 0.893) | 0.001 (p = 0.583) |
| ***n:4F1M:age:sex*** | 0.002 (p = 0.148) | 0.001 (p = 0.102) | 0.001 (p = 0.803) | 0.002 (p = 0.471) |
| ***n:1F4M:age:sex*** | -0.001 (p = 0.246) | 0.000 (p = 0.641) | 0.003 (p = 0.385) | 0.000 (p = 0.924) |
| ***n:1F10M:age:sex*** | -0.000 (p = 0.708) | -0.001 (p = 0.225) | 0.004 (p = 0.210) | -0.002 (p = 0.552) |
| ***n:AD:age:sex*** | 0.001 (p = 0.532) | -0.004 (p < .001) | 0.003 (p = 0.395) | -0.019 (p < .001) |
| ***n:10F1M:AD:age:sex*** | -0.004 (p = 0.040) | -0.001 (p = 0.555) | -0.000 (p = 0.979) | -0.002 (p = 0.596) |
| ***n:4F1M:AD:age:sex*** | -0.003 (p = 0.174) | -0.001 (p = 0.539) | -0.001 (p = 0.896) | -0.001 (p = 0.803) |
| ***n:1F4M:AD:age:sex*** | 0.003 (p = 0.214) | -0.001 (p = 0.589) | -0.004 (p = 0.386) | 0.001 (p = 0.769) |
| ***n:1F10M:AD:age:sex*** | 0.004 (p = 0.046) | -0.001 (p = 0.352) | -0.004 (p = 0.312) | -0.001 (p = 0.853) |

*Table S17. Linear mixed model results for MSE and total outlier count (tOC) under sex-imbalanced sampling for models trained with OASIS-3 or AIBL. Models evaluate the influence of diagnosis (HC, AD), log-transformed and standardized sample size (n), sex ratio in the training set (1:1, 1F:4M, 1F:10M, 4F:1M, 10F:1M), age, sex, and their interactions on standardized deviation score outcomes. Continuous variables were standardized to allow comparison of effect sizes. The 1:1 ratio, HC, and female sex are used as reference levels. Reported β coefficients and corresponding p-values indicate the direction and significance of the effects.*

| | ***OASIS-3*** | | ***AIBL*** | |
|---|---|---|---|---|
| | **MSE β (p-value)** | **tOC β (p-value)** | **MSE β (p-value)** | **tOC β (p-value)** |
| ***Intercept*** | -0.475 (p < .001) | 1.421 (p = 0.448) | -0.475 (p = 0.345) | 7.329 (p = 0.072) |
| ***n*** | -0.172 (p = 0.004) | -0.261 (p < .001) | -0.439 (p = 0.154) | -1.094 (p < .001) |
| ***AD*** | -0.013 (p = 0.893) | 0.355 (p = 0.907) | -0.061 (p = 0.930) | -8.922 (p = 0.116) |
| ***left*** | 0.187 (p = 0.025) | 0.215 (p = 0.002) | -1.989 (p < .001) | -0.726 (p = 0.011) |
| ***right*** | 0.009 (p = 0.918) | 0.397 (p < .001) | 1.258 (p = 0.004) | 5.751 (p < .001) |
| ***n:AD*** | 0.013 (p = 0.889) | 0.110 (p = 0.167) | 0.146 (p = 0.733) | 0.843 (p = 0.003) |
| ***n:left*** | -0.063 (p = 0.447) | 0.019 (p = 0.783) | 0.138 (p = 0.752) | 0.206 (p = 0.469) |
| ***n:right*** | 0.001 (p = 0.990) | 0.109 (p = 0.117) | -0.404 (p = 0.354) | 0.013 (p = 0.964) |
| ***left:AD*** | 0.031 (p = 0.818) | -0.213 (p = 0.058) | 4.092 (p < .001) | 0.250 (p = 0.528) |
| ***right:AD*** | -0.000 (p = 0.999) | 0.058 (p = 0.608) | -0.663 (p = 0.274) | -5.204 (p < .001) |
| ***n:left:AD*** | -0.002 (p = 0.987) | 0.041 (p = 0.712) | 0.040 (p = 0.947) | -0.094 (p = 0.812) |
| ***n:right:AD*** | 0.012 (p = 0.930) | 0.016 (p = 0.889) | 0.556 (p = 0.359) | -0.058 (p = 0.884) |
| ***age*** | -0.000 (p = 0.743) | -0.024 (p = 0.366) | -0.001 (p = 0.848) | -0.104 (p = 0.064) |
| ***left:age*** | 0.000 (p = 0.907) | -0.001 (p = 0.160) | 0.037 (p < .001) | 0.009 (p = 0.021) |
| ***right:age*** | 0.000 (p = 0.994) | -0.005 (p < .001) | -0.012 (p = 0.041) | -0.074 (p < .001) |
| ***AD:age*** | 0.000 (p = 0.850) | 0.002 (p = 0.955) | 0.001 (p = 0.930) | 0.122 (p = 0.117) |
| ***left:AD:age*** | -0.000 (p = 0.940) | 0.004 (p = 0.010) | -0.056 (p < .001) | -0.003 (p = 0.551) |
| ***right:AD:age*** | -0.000 (p = 0.993) | -0.001 (p = 0.662) | 0.009 (p = 0.264) | 0.072 (p < .001) |
| ***n:age*** | 0.000 (p = 0.628) | 0.002 (p = 0.003) | 0.002 (p = 0.599) | 0.014 (p < .001) |
| ***n:left:age*** | -0.000 (p = 0.857) | -0.000 (p = 0.778) | -0.005 (p = 0.444) | -0.002 (p = 0.527) |
| ***n:right:age*** | 0.000 (p = 0.994) | -0.001 (p = 0.161) | 0.004 (p = 0.454) | -0.001 (p = 0.887) |
| ***n:AD:age*** | -0.000 (p = 0.822) | -0.002 (p = 0.127) | -0.002 (p = 0.735) | -0.012 (p = 0.002) |
| ***n:left:AD:age*** | -0.000 (p = 0.960) | -0.001 (p = 0.665) | -0.000 (p = 0.959) | 0.001 (p = 0.810) |
| ***n:right:AD:age*** | -0.000 (p = 0.928) | -0.000 (p = 0.968) | -0.008 (p = 0.360) | 0.001 (p = 0.921) |
| ***sex*** | -0.013 (p = 0.708) | -0.983 (p = 0.377) | -0.088 (p = 0.773) | -3.181 (p = 0.197) |
| ***left:sex*** | -0.025 (p = 0.611) | -0.150 (p < .001) | 0.386 (p = 0.142) | 0.166 (p = 0.335) |
| ***right:sex*** | 0.001 (p = 0.988) | -0.152 (p < .001) | -0.102 (p = 0.697) | -1.733 (p < .001) |
| ***AD:sex*** | 0.017 (p = 0.775) | 1.334 (p = 0.488) | 0.125 (p = 0.768) | 10.016 (p = 0.004) |
| ***left:AD:sex*** | -0.038 (p = 0.654) | 0.320 (p < .001) | -1.816 (p < .001) | -0.124 (p = 0.605) |
| ***right:AD:sex*** | 0.002 (p = 0.982) | 0.079 (p = 0.270) | 0.351 (p = 0.338) | 3.313 (p < .001) |
| ***n:sex*** | 0.019 (p = 0.592) | 0.094 (p = 0.001) | 0.119 (p = 0.521) | 0.170 (p = 0.161) |
| ***n:left:sex*** | 0.015 (p = 0.759) | 0.009 (p = 0.835) | 0.170 (p = 0.520) | 0.042 (p = 0.808) |
| ***n:right:sex*** | 0.002 (p = 0.960) | -0.064 (p = 0.120) | 0.130 (p = 0.622) | -0.178 (p = 0.300) |
| ***n:AD:sex*** | -0.022 (p = 0.710) | -0.181 (p < .001) | -0.186 (p = 0.474) | -0.402 (p = 0.018) |
| ***n:left:AD:sex*** | 0.007 (p = 0.933) | -0.005 (p = 0.938) | -0.263 (p = 0.472) | 0.020 (p = 0.934) |
| ***n:right:AD:sex*** | -0.013 (p = 0.881) | 0.032 (p = 0.654) | -0.333 (p = 0.364) | 0.257 (p = 0.284) |
| ***age:sex*** | 0.000 (p = 0.715) | 0.013 (p = 0.435) | 0.001 (p = 0.774) | 0.042 (p = 0.216) |
| ***left:age:sex*** | 0.001 (p = 0.484) | 0.002 (p = 0.005) | -0.005 (p = 0.138) | -0.002 (p = 0.403) |
| ***right:age:sex*** | -0.000 (p = 0.975) | 0.002 (p < .001) | 0.001 (p = 0.725) | 0.023 (p < .001) |
| ***AD:age:sex*** | -0.000 (p = 0.761) | -0.017 (p = 0.511) | -0.002 (p = 0.778) | -0.129 (p = 0.006) |
| ***left:AD:age:sex*** | 0.000 (p = 0.814) | -0.004 (p < .001) | 0.024 (p < .001) | 0.001 (p = 0.687) |
| ***right:AD:age:sex*** | -0.000 (p = 0.996) | -0.001 (p = 0.345) | -0.005 (p = 0.367) | -0.043 (p < .001) |
| ***n:age:sex*** | -0.000 (p = 0.608) | -0.001 (p = 0.020) | -0.002 (p = 0.522) | -0.002 (p = 0.174) |
| ***n:left:age:sex*** | -0.000 (p = 0.693) | -0.000 (p = 0.795) | -0.002 (p = 0.521) | -0.001 (p = 0.799) |
| ***n:right:age:sex*** | -0.000 (p = 0.971) | 0.001 (p = 0.157) | -0.002 (p = 0.637) | 0.002 (p = 0.335) |

| | | | | |
|---|---|---|---|---|
| ***n:AD:age:sex*** | 0.000 (p = 0.688) | 0.002 (p = 0.002) | 0.002 (p = 0.493) | 0.005 (p = 0.024) |
| ***n:left:AD:age:sex*** | -0.000 (p = 0.991) | 0.000 (p = 0.891) | 0.004 (p = 0.471) | -0.000 (p = 0.956) |
| ***n:right:AD:age:sex*** | 0.000 (p = 0.897) | -0.000 (p = 0.666) | 0.004 (p = 0.379) | -0.003 (p = 0.311) |

*Table S18. Linear mixed model results for MSE and total outlier count (tOC) under age-skewed sampling for models trained with UKB and transferred to OASIS-3 or AIBL. Models evaluate the influence of diagnosis (HC, AD), log-transformed and standardized sample size (n), age sampling strategy (Representative, Left-skewed, Right-skewed), age, sex, and their interactions on standardized deviation score outcomes. Continuous variables were standardized to allow comparison of effect sizes. Representative sampling, HC, and female sex are used as reference levels. Reported β coefficients and corresponding p-values indicate the direction and significance of the effects.*

| | ***OASIS-3*** | | ***AIBL*** | |
|---|---|---|---|---|
| | **MSE β (p-value)** | **tOC β (p-value)** | **MSE β (p-value)** | **tOC β (p-value)** |
| ***Intercept*** | NA | 1.421 (p = 0.433) | -0.475 (p = 0.022) | 7.329 (p = 0.062) |
| ***n*** | -0.172 (p < .001) | -0.261 (p < .001) | -0.439 (p = 0.034) | -1.094 (p < .001) |
| ***AD*** | 0.000 (p = 1.000) | 0.355 (p = 0.904) | -0.061 (p = 0.831) | -8.922 (p = 0.103) |
| ***10F1M*** | 0.014 (p = 0.822) | 0.323 (p < .001) | -0.026 (p = 0.929) | -0.038 (p = 0.844) |
| ***4F1M*** | 0.006 (p = 0.929) | 0.216 (p < .001) | -0.027 (p = 0.925) | 0.033 (p = 0.864) |
| ***1F4M*** | 0.015 (p = 0.821) | -0.009 (p = 0.880) | 0.088 (p = 0.763) | 0.435 (p = 0.025) |
| ***1F10M*** | 0.016 (p = 0.809) | 0.027 (p = 0.647) | 0.072 (p = 0.805) | 0.425 (p = 0.029) |
| ***n:AD*** | 0.013 (p = 0.856) | 0.110 (p = 0.108) | 0.146 (p = 0.611) | 0.843 (p < .001) |
| ***n:10F1M*** | 0.006 (p = 0.920) | 0.013 (p = 0.821) | 0.091 (p = 0.756) | 0.130 (p = 0.504) |
| ***n:4F1M*** | 0.007 (p = 0.910) | -0.002 (p = 0.968) | 0.072 (p = 0.805) | 0.009 (p = 0.963) |
| ***n:1F4M*** | -0.011 (p = 0.858) | -0.002 (p = 0.969) | -0.050 (p = 0.865) | -0.177 (p = 0.361) |
| ***n:1F10M*** | 0.003 (p = 0.964) | -0.009 (p = 0.878) | 0.130 (p = 0.656) | 0.029 (p = 0.883) |
| ***10F1M:AD*** | 0.004 (p = 0.969) | -0.409 (p < .001) | 0.050 (p = 0.902) | 0.424 (p = 0.117) |
| ***4F1M:AD*** | 0.000 (p = 0.999) | -0.267 (p = 0.006) | 0.021 (p = 0.959) | 0.347 (p = 0.200) |
| ***1F4M:AD*** | -0.003 (p = 0.974) | -0.002 (p = 0.983) | -0.135 (p = 0.741) | -0.537 (p = 0.047) |
| ***1F10M:AD*** | -0.000 (p = 0.997) | -0.015 (p = 0.873) | -0.128 (p = 0.754) | -0.756 (p = 0.005) |
| ***n:10F1M:AD*** | -0.008 (p = 0.937) | 0.015 (p = 0.878) | -0.105 (p = 0.796) | -0.513 (p = 0.058) |
| ***n:4F1M:AD*** | -0.004 (p = 0.969) | 0.049 (p = 0.611) | -0.047 (p = 0.909) | -0.301 (p = 0.265) |
| ***n:1F4M:AD*** | 0.014 (p = 0.893) | 0.019 (p = 0.847) | 0.130 (p = 0.750) | 0.198 (p = 0.465) |
| ***n:1F10M:AD*** | 0.004 (p = 0.968) | 0.037 (p = 0.698) | -0.108 (p = 0.791) | 0.127 (p = 0.640) |
| ***age*** | 0.000 (p = 1.000) | -0.024 (p = 0.350) | -0.001 (p = 0.641) | -0.104 (p = 0.055) |
| ***10F1M:age*** | -0.000 (p = 0.943) | -0.004 (p < .001) | 0.001 (p = 0.849) | 0.001 (p = 0.677) |
| ***4F1M:age*** | -0.000 (p = 0.963) | -0.003 (p = 0.003) | 0.001 (p = 0.895) | -0.000 (p = 0.994) |
| ***1F4M:age*** | -0.000 (p = 0.932) | -0.000 (p = 0.968) | -0.001 (p = 0.826) | -0.006 (p = 0.027) |
| ***1F10M:age*** | -0.000 (p = 0.977) | -0.000 (p = 0.567) | -0.000 (p = 0.921) | -0.006 (p = 0.028) |
| ***AD:age*** | -0.000 (p = 1.000) | 0.002 (p = 0.954) | 0.001 (p = 0.831) | 0.122 (p = 0.104) |
| ***10F1M:AD:age*** | -0.000 (p = 0.988) | 0.006 (p < .001) | -0.001 (p = 0.905) | -0.005 (p = 0.156) |
| ***4F1M:AD:age*** | 0.000 (p = 0.992) | 0.004 (p = 0.002) | -0.000 (p = 0.963) | -0.004 (p = 0.236) |
| ***1F4M:AD:age*** | 0.000 (p = 0.967) | -0.000 (p = 0.895) | 0.002 (p = 0.738) | 0.007 (p = 0.050) |
| ***1F10M:AD:age*** | 0.000 (p = 0.989) | 0.000 (p = 0.928) | 0.002 (p = 0.752) | 0.010 (p = 0.006) |
| ***n:age*** | 0.000 (p = 0.529) | 0.002 (p < .001) | 0.002 (p = 0.434) | 0.014 (p < .001) |
| ***n:10F1M:age*** | -0.000 (p = 0.980) | -0.000 (p = 0.831) | -0.001 (p = 0.747) | -0.002 (p = 0.475) |
| ***n:4F1M:age*** | -0.000 (p = 0.991) | 0.000 (p = 0.939) | -0.001 (p = 0.802) | -0.000 (p = 0.893) |
| ***n:1F4M:age*** | 0.000 (p = 0.862) | 0.000 (p = 0.887) | 0.001 (p = 0.852) | 0.002 (p = 0.378) |
| ***n:1F10M:age*** | -0.000 (p = 0.993) | 0.000 (p = 0.806) | -0.002 (p = 0.662) | -0.000 (p = 0.888) |
| ***n:AD:age*** | -0.000 (p = 0.770) | -0.002 (p = 0.076) | -0.002 (p = 0.614) | -0.012 (p < .001) |
| ***n:10F1M:AD:age*** | 0.000 (p = 0.949) | -0.000 (p = 0.894) | 0.001 (p = 0.800) | 0.007 (p = 0.062) |
| ***n:4F1M:AD:age*** | 0.000 (p = 0.978) | -0.001 (p = 0.619) | 0.001 (p = 0.913) | 0.004 (p = 0.271) |
| ***n:1F4M:AD:age*** | -0.000 (p = 0.886) | -0.000 (p = 0.859) | -0.002 (p = 0.746) | -0.003 (p = 0.463) |
| ***n:1F10M:AD:age*** | -0.000 (p = 0.969) | -0.001 (p = 0.648) | 0.001 (p = 0.798) | -0.002 (p = 0.660) |
| ***sex*** | NA | -0.983 (p = 0.361) | -0.088 (p = 0.482) | -3.181 (p = 0.181) |
| ***10F1M:sex*** | -0.004 (p = 0.907) | -0.165 (p < .001) | 0.016 (p = 0.926) | 0.133 (p = 0.259) |
| ***4F1M:sex*** | -0.003 (p = 0.939) | -0.114 (p = 0.001) | 0.002 (p = 0.990) | 0.040 (p = 0.734) |
| ***1F4M:sex*** | -0.003 (p = 0.943) | 0.016 (p = 0.650) | -0.045 (p = 0.801) | -0.157 (p = 0.181) |
| ***1F10M:sex*** | -0.000 (p = 0.995) | 0.008 (p = 0.822) | -0.055 (p = 0.755) | -0.148 (p = 0.207) |
| ***AD:sex*** | NA | 1.334 (p = 0.474) | 0.125 (p = 0.473) | 10.016 (p = 0.002) |
| ***10F1M:AD:sex*** | 0.001 (p = 0.988) | 0.335 (p < .001) | -0.003 (p = 0.989) | -0.107 (p = 0.514) |
| ***4F1M:AD:sex*** | 0.001 (p = 0.989) | 0.213 (p < .001) | 0.008 (p = 0.975) | -0.105 (p = 0.520) |
| ***1F4M:AD:sex*** | 0.001 (p = 0.989) | -0.022 (p = 0.715) | 0.086 (p = 0.728) | 0.308 (p = 0.060) |
| ***1F10M:AD:sex*** | 0.003 (p = 0.970) | -0.002 (p = 0.968) | 0.111 (p = 0.652) | 0.472 (p = 0.004) |
| ***n:sex*** | 0.019 (p = 0.486) | 0.094 (p < .001) | 0.119 (p = 0.340) | 0.170 (p = 0.040) |
| ***n:10F1M:sex*** | 0.002 (p = 0.958) | -0.006 (p = 0.873) | -0.027 (p = 0.879) | -0.064 (p = 0.588) |
| ***n:4F1M:sex*** | 0.002 (p = 0.957) | 0.002 (p = 0.946) | -0.005 (p = 0.978) | 0.002 (p = 0.986) |
| ***n:1F4M:sex*** | 0.006 (p = 0.879) | 0.003 (p = 0.928) | 0.030 (p = 0.867) | 0.097 (p = 0.408) |
| ***n:1F10M:sex*** | -0.003 (p = 0.944) | 0.006 (p = 0.864) | -0.011 (p = 0.952) | 0.044 (p = 0.705) |
| ***n:AD:sex*** | -0.022 (p = 0.629) | -0.181 (p < .001) | -0.186 (p = 0.286) | -0.402 (p < .001) |
| ***n:10F1M:AD:sex*** | 0.001 (p = 0.992) | 0.005 (p = 0.933) | 0.036 (p = 0.885) | 0.288 (p = 0.079) |
| ***n:4F1M:AD:sex*** | 0.001 (p = 0.992) | -0.003 (p = 0.957) | -0.003 (p = 0.989) | 0.135 (p = 0.408) |
| ***n:1F4M:AD:sex*** | -0.002 (p = 0.977) | 0.016 (p = 0.794) | -0.067 (p = 0.784) | -0.085 (p = 0.604) |
| ***n:1F10M:AD:sex*** | -0.002 (p = 0.971) | 0.001 (p = 0.985) | 0.010 (p = 0.969) | -0.100 (p = 0.541) |
| ***age:sex*** | 0.000 (p = 1.000) | 0.013 (p = 0.421) | 0.001 (p = 0.485) | 0.042 (p = 0.199) |

| | | | | |
|---|---|---|---|---|
| ***10F1M:age:sex*** | 0.000 (p = 0.910) | 0.002 (p < .001) | -0.000 (p = 0.920) | -0.002 (p = 0.221) |
| ***4F1M:age:sex*** | 0.000 (p = 0.937) | 0.001 (p = 0.006) | -0.000 (p = 0.986) | -0.001 (p = 0.666) |
| ***1F4M:age:sex*** | 0.000 (p = 0.930) | -0.000 (p = 0.773) | 0.001 (p = 0.797) | 0.002 (p = 0.166) |
| ***1F10M:age:sex*** | 0.000 (p = 0.989) | -0.000 (p = 0.962) | 0.001 (p = 0.748) | 0.002 (p = 0.175) |
| ***AD:age:sex*** | 0.000 (p = 1.000) | -0.017 (p = 0.497) | -0.002 (p = 0.494) | -0.129 (p = 0.004) |
| ***10F1M:AD:age:sex*** | -0.000 (p = 0.981) | -0.005 (p < .001) | 0.000 (p = 0.987) | 0.002 (p = 0.488) |
| ***4F1M:AD:age:sex*** | -0.000 (p = 0.984) | -0.003 (p < .001) | -0.000 (p = 0.974) | 0.001 (p = 0.505) |
| ***1F4M:AD:age:sex*** | -0.000 (p = 0.980) | 0.000 (p = 0.682) | -0.001 (p = 0.729) | -0.004 (p = 0.070) |
| ***1F10M:AD:age:sex*** | -0.000 (p = 0.973) | 0.000 (p = 0.945) | -0.002 (p = 0.653) | -0.006 (p = 0.006) |
| ***n:age:sex*** | -0.000 (p = 0.505) | -0.001 (p = 0.007) | -0.002 (p = 0.341) | -0.002 (p = 0.047) |
| ***n:10F1M:age:sex*** | -0.000 (p = 0.948) | 0.000 (p = 0.858) | 0.000 (p = 0.871) | 0.001 (p = 0.568) |
| ***n:4F1M:age:sex*** | -0.000 (p = 0.941) | -0.000 (p = 0.953) | 0.000 (p = 0.970) | 0.000 (p = 0.966) |
| ***n:1F4M:age:sex*** | -0.000 (p = 0.839) | -0.000 (p = 0.872) | -0.000 (p = 0.871) | -0.001 (p = 0.414) |
| ***n:1F10M:age:sex*** | 0.000 (p = 0.960) | -0.000 (p = 0.794) | 0.000 (p = 0.953) | -0.001 (p = 0.706) |
| ***n:AD:age:sex*** | 0.000 (p = 0.601) | 0.002 (p < .001) | 0.002 (p = 0.308) | 0.005 (p < .001) |
| ***n:10F1M:AD:age:sex*** | -0.000 (p = 0.995) | -0.000 (p = 0.940) | -0.000 (p = 0.886) | -0.004 (p = 0.084) |
| ***n:4F1M:AD:age:sex*** | -0.000 (p = 0.996) | 0.000 (p = 0.928) | 0.000 (p = 0.989) | -0.002 (p = 0.400) |
| ***n:1F4M:AD:age:sex*** | 0.000 (p = 0.955) | -0.000 (p = 0.847) | 0.001 (p = 0.782) | 0.001 (p = 0.620) |
| ***n:1F10M:AD:age:sex*** | 0.000 (p = 0.981) | 0.000 (p = 0.930) | -0.000 (p = 0.979) | 0.001 (p = 0.573) |

*Table S19. Linear mixed model results for MSE and total outlier count (tOC) under sex-imbalanced sampling for models trained with UKB and transferred to OASIS-3 or AIBL. Models evaluate the influence of diagnosis (HC, AD), log-transformed and standardized sample size (n), sex ratio in the training set (1:1, 1F:4M, 1F:10M, 4F:1M, 10F:1M), age, sex, and their interactions on standardized deviation score outcomes. Continuous variables were standardized to allow comparison of effect sizes. The 1:1 ratio, HC, and female sex are used as reference levels. Reported β coefficients and corresponding p-values indicate the direction and significance of the effects.*

#### 5.3. Control analysis for instability around n ≈ 300

To determine whether the deviation observed at *n* = 300 under left-skewed sampling reflected a systematic modeling issue or stochastic sampling variability, we repeated the model-fitting procedure at sample size 300 using 20 independent random seeds. For each seed, training sets were newly generated following the same sampling schemes and models were re-estimated using the standard pipeline. Model performance in the HC test set was evaluated using the same metrics as in Figure 3 (MSLL, SMSE, EV, Rho, lower-tail HC%, upper-tail HC%, and ICC), and variability was quantified across seeds (or across ROIs for ICC). The original sampling results were retained in the main manuscript for transparency and to reflect the variability inherent in individual sampling draws.

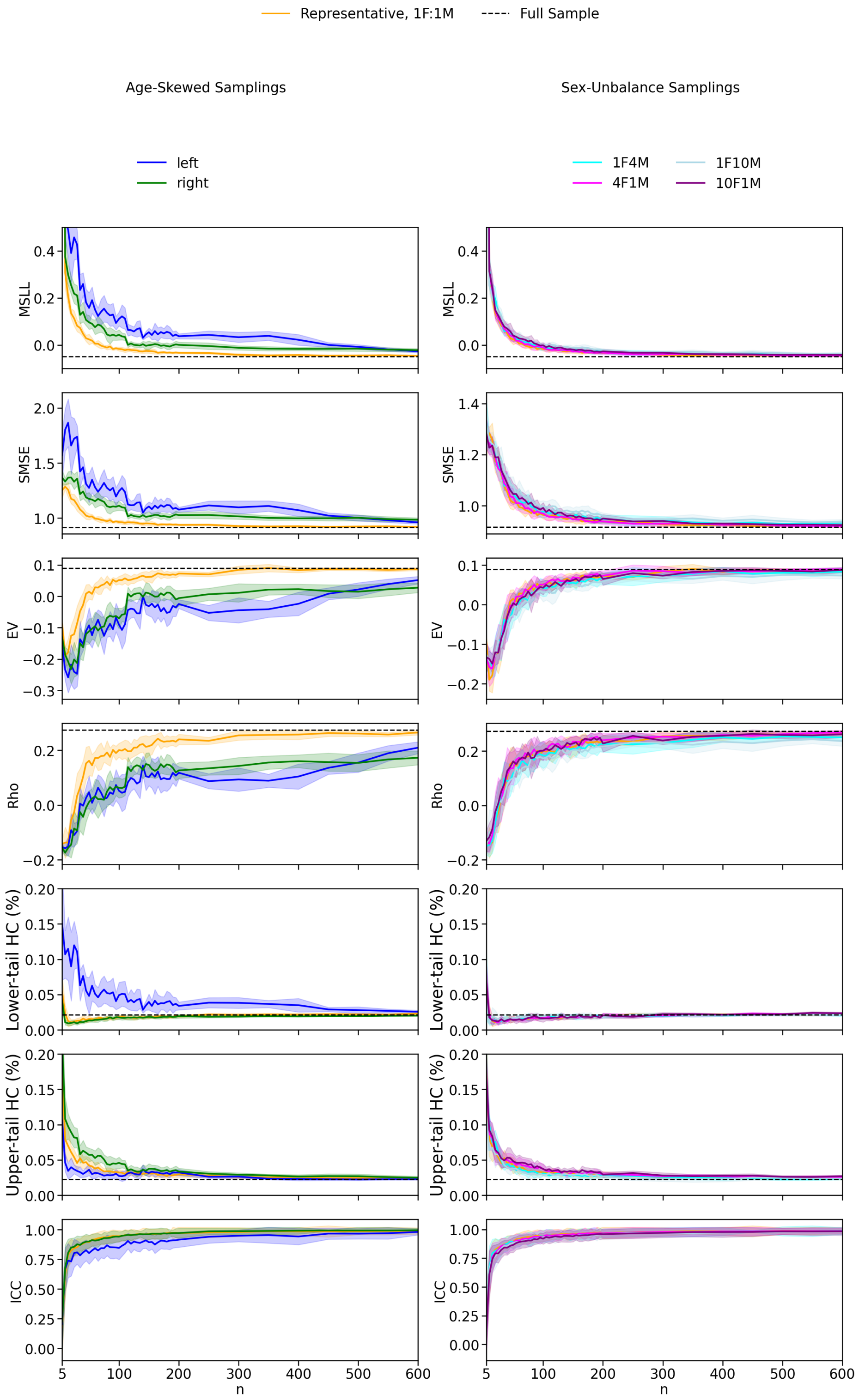

*Figure S36. Evaluation of model fits in the HC test set across different sampling strategies of the training set for varying sample sizes (n), following re-estimation with 20 independent random seeds to assess the robustness of an apparent artifact observed at n = 300 in left skewed sampling in the original analysis. Sampling strategies and metrics are identical to those shown in Figure 3. Solid lines represent the mean metric values across ROIs and random seeds, with shaded areas indicating the standard deviation across seeds for MSLL, SMSE, EV, Rho, the lower and upper tail HC%. For ICC, shaded areas represent variability across ROIs. Dashed lines indicate the mean metric values obtained with the full sample. The absence of a systematic deviation at n = 300 across random seeds indicates that the previously observed effect was driven by stochastic sampling variability rather than a stable modeling artifact.*

##### 5.4. Age-distribution coverage between sampled training sets and test cohorts

To quantify how well each sampling strategy aligned the age distribution of the sampled training sets with the empirical age distributions of the HC test cohorts, we computed an age-bin coverage metric based on distribution intersection. Age was discretized into 20 quantile-based bins using the full training set of each dataset (OASIS-3 and AIBL) as reference.

For each sampling strategy (Representative, Left-skewed, Right-skewed), sample size, and dataset, we generated 1000 independent training samples using the same sampling procedures as in the main analyses. For each sampled training set, age-bin count distributions were computed and compared to the corresponding HC test-set age-bin counts.

Coverage was defined as:

$$\text{Coverage} = \frac{\sum_i \min\left(n_{\text{train}}(i),\, n_{\text{test}}(i)\right)}{\sum_i n_{\text{test}}(i)}$$

where, $i$ indexes age bins, $n_{\text{train}}$ and $n_{\text{test}}$ are the numbers of individuals in bin i in the sampled training set and HC test set, respectively. This metric quantifies the fraction of the test-set age distribution that is "covered" by the sampled training set and ranges from 0 (no test-set ages covered) to 1 (complete coverage of the test-set age distribution). For each condition, the mean and standard deviation of the coverage across repetitions were computed.

This analysis quantifies how the imposed sampling-induced age profiles align with the empirical age structure of the target populations and how this alignment evolves with increasing sample size.

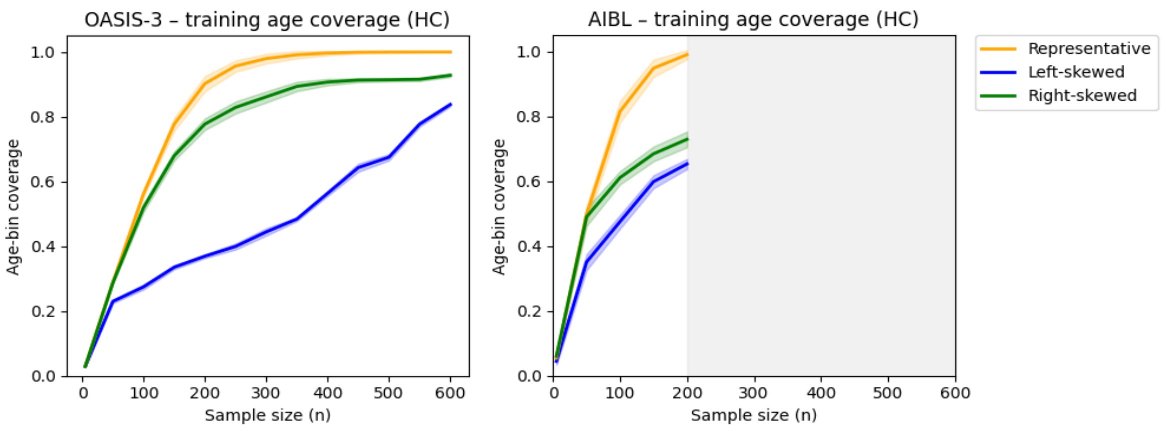

*Figure S37. Training age-distribution coverage with respect to the HC test cohorts in OASIS-3 (left) and AIBL (right). Mean age-distribution coverage (± SD across 1000 repetitions) between the sampled training sets and the HC test cohorts is shown for each age-sampling strategy as a function of training sample size (n). Coverage is defined as the distribution intersection between age-bin counts of the sampled training data and the HC test sets, using 20 quantile-based bins derived from the full training sets of each dataset. In OASIS-3, representative sampling yields the highest coverage, while left-skewed sampling remains consistently lower than right-skewed sampling across sample sizes, reflecting the older and skewed age distribution of the cohort. In AIBL, representative sampling also provides the highest coverage and left-skewed sampling remains slightly below right-skewed sampling but with reduced differences compared to OASIS-3. Sampling was only performed up to n = 200 due to sample availability in AIBL; the grey region marks larger sample sizes not evaluated, while the x-axis is extended to n = 600 for comparability across datasets.*
